# Optimization and Characterization of SHIP1 Ligands for Cellular Target Engagement and Activity in Alzheimer’s Disease Models

**DOI:** 10.64898/2025.12.31.697127

**Authors:** Cynthia D. Jesudason, Claudia Rangel-Barajas, Colin J. Beach, Daniel E. Beck, Isaac H. Caballero-Floran, W. Brent Clayton, Lais Da Silva, Juliane C. David, Suzanne Doolen, Alexandra N. Faulkner, Adam K. Hamdani, Hasi Huhe, Kevin Huynh, Ryan D. Imhoff, June Javens-Wolfe, Emily R. Mason, Mustapha Moussaif, Kratika Singhal, Disha M. Soni, Matilda van Buuren-Milne, Sean-Paul Williams, Steven P. Angus, Shaoyou Chu, Jeffrey L. Dage, Philip A. Hipskind, Travis S. Johnson, Rima Kaddurah-Daouk, Bruce T. Lamb, Peter J. Meikle, Andrew D. Mesecar, Alan D. Palkowitz, Sara K. Quinney, Stacey J. Sukoff Rizzo, Adrian L. Oblak, Timothy I. Richardson

**Author notes:** Corresponding Author: Timothy I. Richardson, Ph.D; 1210 Waterway Blvd., Ste. 2000, Indianapolis, IN 46202. These authors contributed equally to this work (Cynthia D. Jesudason, Claudia Rangel-Barajas).

## Abstract

Src homology 2 domain-containing inositol 5-phosphatase 1 (SHIP1), encoded by the gene INPP5D, is a lipid phosphatase that negatively regulates immune receptor signaling in hematopoietic cells and microglia. Here, we describe a pyridyl-pyrazole-piperidine scaffold and the lead compound 3-((2-chlorobenzyl)oxy)-5-(1-(piperidin-4-yl)-1H-pyrazol-4-yl)pyridine (**32**), which demonstrates SHIP1 target engagement, brain exposure, and evidence of a central pharmacodynamic response *in vivo*. Structure-activity relationship studies, guided by biochemical and cellular assays using multiple human and murine protein constructs and cells, identified SHIP1-active ligands. A thermal shift assay using full-length SHIP1 was used to assess compounds for cellular target engagement, while studies in IL-4 conditioned THP-1 cells was used to demonstrate changes in downstream AKT signaling. Targeted lipidomics revealed changes in the overall phosphoinositide pool consistent with SHIP1 target engagement and reduction of phospho-AKT levels. In a protein-lipid overlay assay, compound **32** induced changes in the relative association of SHIP1 with multiple phosphatidylinositols on a membrane surface. In high-content cellular imaging assays, compound **32** enhanced the uptake of myelin/membrane debris and fibrillar amyloid by primary murine microglia, phenocopying a genetic model with reduced SHIP1 expression. Finally, oral administration of compound **32** resulted in brain exposure sufficient to alter gene expression and reduce IL-1β levels as pharmacodynamic markers of microglial activation and neuroinflammation in an amyloidosis mouse model of Alzheimer’s disease. Collectively, these results define a scaffold with SHIP1 target engagement, CNS exposure, and *in vivo* activity, providing a foundation for the optimization of brain-penetrant SHIP1 ligands suitable for further mechanistic studies and therapeutic development for the treatment of Alzheimer’s disease.

## INTRODUCTION

Src homology 2 domain containing inositol polyphosphate 5-phosphatase 1 (SHIP1) is a member of the inositol 5-phosphatase subclass of phosphatidylinositol phosphatases. SHIP1, encoded by the gene *INPP5D*, is predominantly expressed in hematopoietic cells, especially within lymphoid tissues, where it plays an important role as a negative regulator of immune receptor signaling regulating cellular proliferation, survival, and chemotaxis. Its closest homolog, SHIP2, encoded by *INPPL1*, is expressed more widely across tissues, including liver, kidney, and muscle, where it plays important roles in metabolic pathways and insulin signaling. Both homologs have been studied as oncogenes, although the roles they play as tumor suppressors are highly dependent on cancer type and cellular context. In hematologic malignancies, SHIP1 generally acts as a tumor suppressor by negatively regulating the PI3K/AKT signaling pathway^1^. More recently *INPP5D* has been identified as a risk gene for Alzheimer’s Disease (AD) and multiple isoforms of the transcript and the protein have been reported that are differentially expressed in AD compared to normal controls^2-4^. The functions of *INPP5D*/SHIP1 in neurodegeneration, particularly its role in regulating microglial states, are a major focus of several research groups, including our own^3, 5-10^.

SHIP1 is a multi-domain, multi-functional protein that associates with cellular membranes where it is involved in the inositol phospholipid signaling pathway. The catalytic phosphatase domain (Ptase) is flanked by a pleckstrin-homology (PH) domain that binds to phosphatidylinositol (3,4,5)-trisphosphate [PI(3,4,5)P_3_] and a C2 domain that binds to phosphatidylinositol (3,4)-bisphosphate [PI(3,4)P_2_] (**Figure 1**)^11, 12^. The catalytic site of the Ptase domain converts PI(3,4,5)P_3_ to PI(3,4)P_2_, each of which recruit PH-containing enzymes and kinases (e.g. PLCγ2, BTK, PDK1, and AKT) involved in signal transduction. Interactions between the Ptase and C2 domains modulate enzymatic activity^13, 14^. Membrane localization of SHIP1 is facilitated by an N-terminal SH2 domain, which binds immunoreceptor tyrosine-based activation and inhibition motifs (ITAMs and ITIMs) of immune receptors (e.g. BCR, FcγRIIB)^15^. Studies of immune receptor activation (e.g. FcεRI in mast cells) have shown that SHIP1 undergoes cycles of association and dissociation with the membrane surface, inducing cortical oscillations of the plasma membrane^16^. The disordered C-terminal proline-rich region contains NPXY and PXXP motifs that also aid in membrane localization and regulation. NPXY motifs are critical for receptor-mediated endocytosis and intracellular trafficking while PXXP motifs mediate protein-protein interactions in signaling pathways and cytoskeletal dynamics. The SHIP1 NPXY motif undergoes phosphorylation and has been reported to interact with the phosphotyrosine-binding (PTB) domains of adaptor proteins (e.g. DOK1, DOK2)^17, 18^. The PXXP motif has been shown to engage with SH3-containing proteins (e.g. GRB2)^19^.

**Figure 1.**
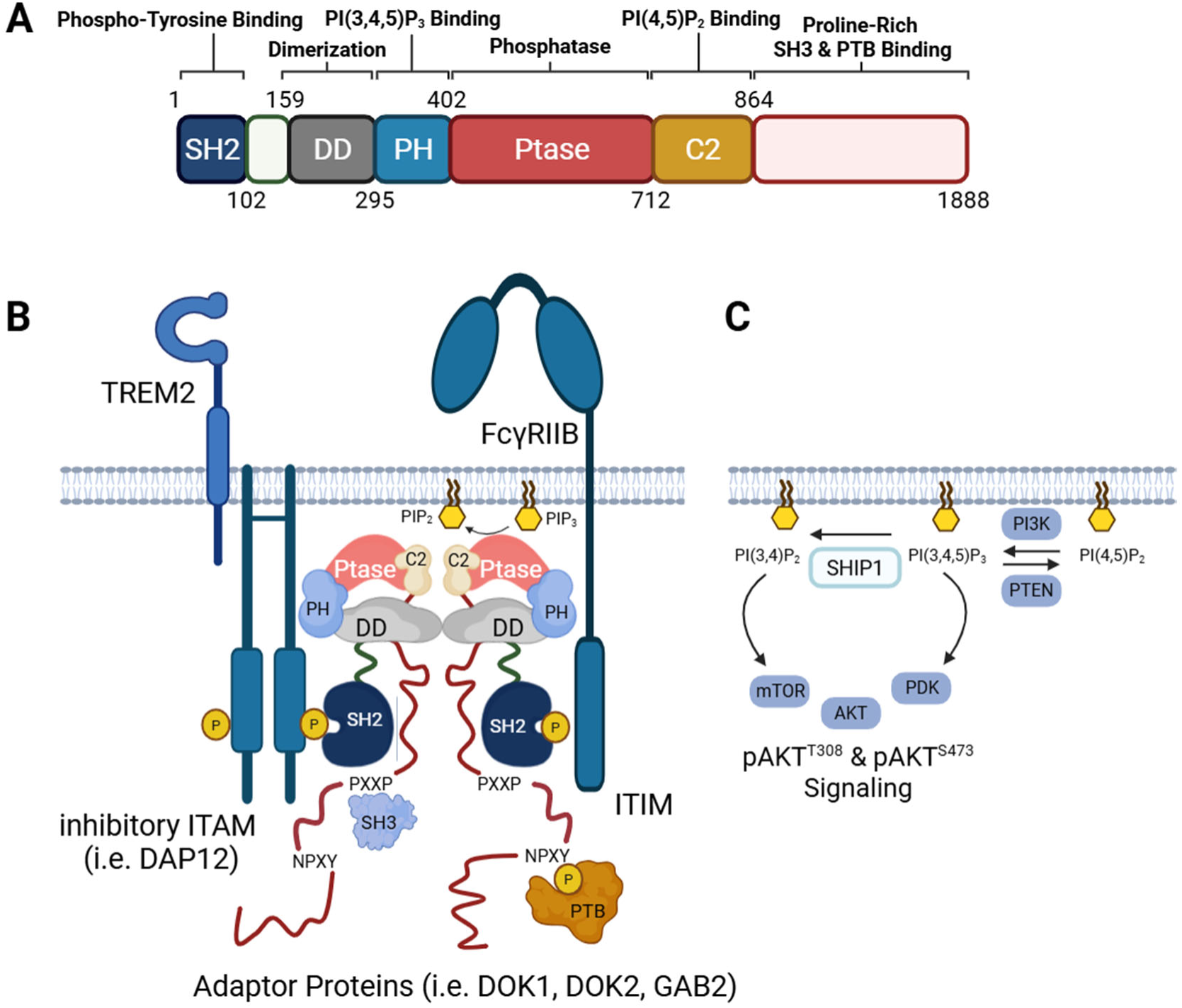
Structure and Function of SHIP1. **A.** Functional domains of SHIP1 with their approximate amino acid boundaries. **B.** Model illustrating SHIP1 dimerization and interactions with immunoreceptor tyrosine-based activation motifs (ITAMs) and inhibition motifs (ITIMs). The catalytic phosphatase (Ptase) converts PI(3,4,5)P_3_ to PI(3,4)P_2_. The pleckstrin-homology (PH) domain interacts with PI(3,4,5)P_3_. The C2 domain associates with PI(3,4)P_2_. The SH2 domain at the N-terminus interacts with inhibitory ITAMs and ITIMs of immune receptors. The disordered C-terminal region includes a proline-rich PXXP motif that interacts with SH3 domain containing proteins and an NPXY motif, which can be phosphorylated and interact with the phosphotyrosine-binding (PTB) domains of adaptor proteins. **C.** SHIP1 negatively regulates the PI3K/AKT pathway. The conversion of PI(3,4,5)P_3_ to PI(3,4)P_2_ by SHIP1 changes the overall phosphoinositide pool of lipids, which affects membrane dynamics, and modulates AKT signaling by influencing the activity of PH domain containing proteins such as PDK and mTORC2 (through SIN1) as well as AKT itself. Created in BioRender.com.

The dimerization and oligomerization of immune receptors on the surface of leukocytes is a well-studied phenomenon^20^. As a negative regulator downstream, SHIP1 has been reported to dimerize and form oligomers through interactions between the SH2 and C-terminal domains in the context of B cell receptor signaling, which may impact its function and interactions with other proteins^21^. Recently, we identified an internal dimerization domain (DD) that may assist in bridging immunoreceptor tyrosine-based inhibitory motifs (ITIMs) and/or inhibitory immunoreceptor tyrosine-based activation motifs (ITAMs)^22^. Others have functionally characterized this domain and shown that the N-terminal SH2 domain suppresses phosphatase activity through intramolecular contacts between the SH2 and C2 domains that mediate autoinhibition^23, 24^. For illustrative purposes, the model depicted in **Figure 1** shows the multifunctional nature of SHIP1 in its activated form in which it negatively regulates signals from various immune receptors clustered on the cell surface, by showing binding to an ITIM and an inhibitory ITAM. The membrane binding of its PH and C2 domains are shown along with its functional Ptase domain, which converts PI(3,4,5)P_3_ to PI(3,4)P_2_. Other potential protein interactions with SHIP1 are not shown for simplicity. This model describes the modular and adaptable nature of SHIP1-mediated signaling that may be modulated by small molecules potentially interacting with any of the SHIP1 domains, other protein binding partners, or within the cell membrane to which SHIP1 associates.

The Kerr lab has identified several inhibitors targeting SHIP1 and SHIP2 (**Figure 2**)^25^. Among these, the aminosteroid **1**, known as 3α-aminocholestane (3AC)^26^, has been the most extensively utilized^27-38^. In our hands the free base of 3AC was insoluble. As a result, we prepared the HCl salt, which was soluble in a DMSO stock solution. K116 (**2**) is a more soluble aminosteroid analog of 3AC^37, 39^. An *in vivo* study in 3-month-old C57BL/6 mice has been reported using the aminosteroid analog K161 (**3**), a pan-SHIP1/2 inhibitor that did not have sufficient exposure to demonstrate a meaningful pharmacodynamic response, failing to alter the frequency of major cell populations in the cerebral cortex or the surface density of receptors on microglia^37^. Several compounds based on a tryptamine scaffold (**4**) have also been reported^27^. A thiophene scaffold represented by AS1949490 (**5**) was described by Astellas Pharma^40, 41^ as a selective SHIP2 inhibitor with some activity against SHIP1. The pyrazole NGD-61338 (**6**), discovered by NeoGenesis Pharmaceuticals using affinity-based screening of a combinatorial library against SHIP2^42^, had some activity against the SHIP1 enzyme in our hands (*see below*). Researchers at the University of Toyama noted similarities between inhibitors reported by Astellas and NeoGenesis and used them as a starting point for a ligand-based design effort that culminated in the synthesis and evaluation of a pyridyl-based scaffold best represented by CPDA (**7**)^43^. Similar scaffolds represented by compounds **8** & **9** were reported by Lim *et al.* based on the observation that crizotinib (**10**), a multi-targeted kinase inhibitor, is also an inhibitor of SHIP2^44^. Most recently, the competitive SHIP1 inhibitors SP1-3 (**11**-**13**) were discovered using a high throughput screen of the SHIP1^397-715^ Ptase domain^45^. Previously, we described the evaluation reported ligands for SHIP1 and concluded that compound **9** engages SHIP1 without cytotoxicity^7^. Here we describe structure activity relationship studies around **9** (**Figure 3**).

**Figure 2.**
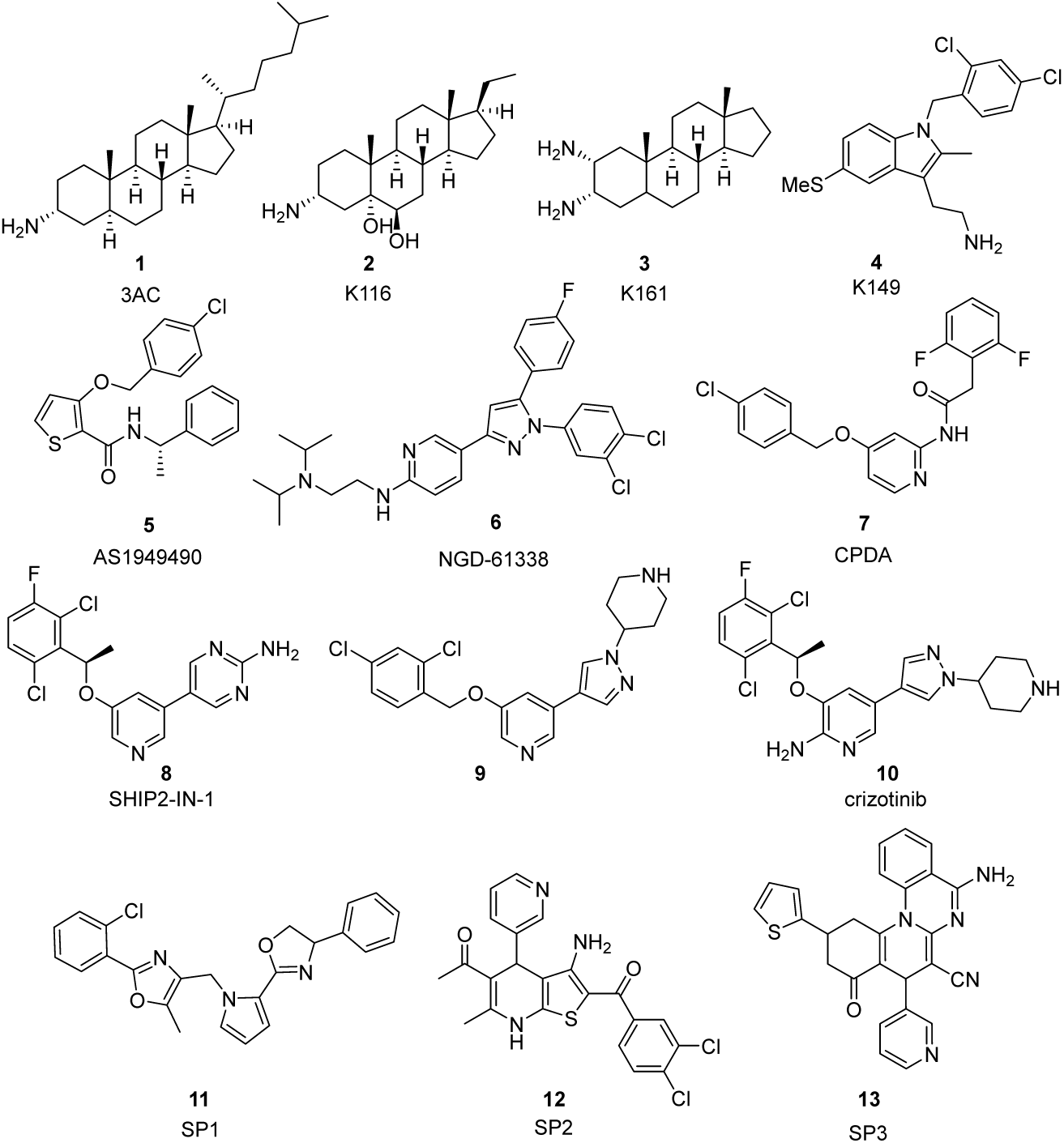
SHIP1 and SHIP2 reference compounds.

**Figure 3.**
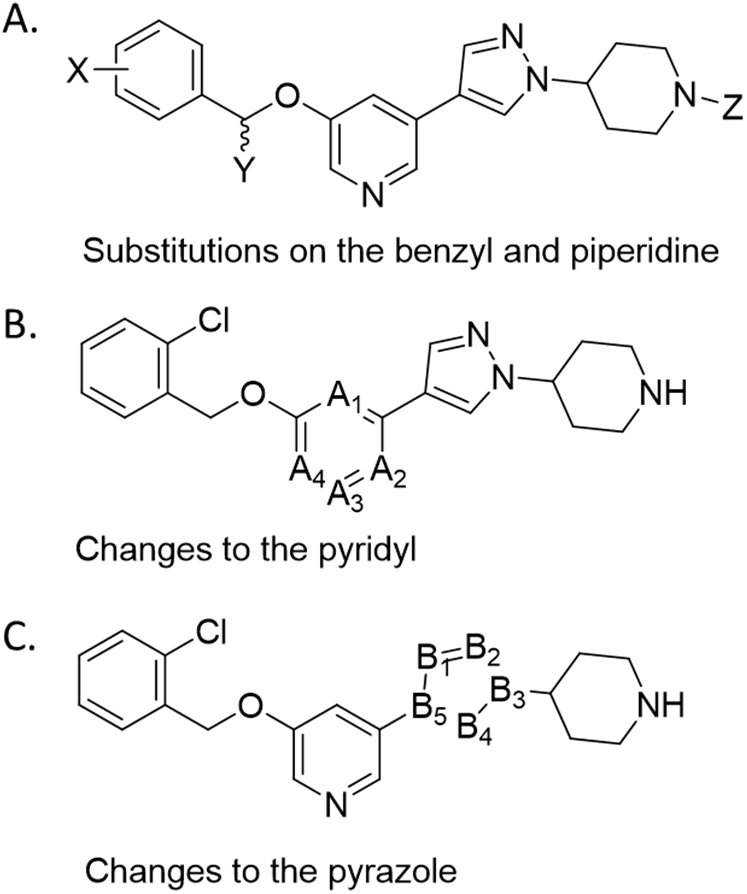
Structure activity relationship (SAR) studies. (A) Substitutions X on the benzyl ring, Y at the α-carbon, and Z on piperidine ring nitrogen atom. (B) Modifications to the pyridyl ring removing and shifting the position (A1-A4) of the nitrogen atom. (C) Modifications of the pyrazole ring varying the number and positions (B1-B5) of the nitrogen ring atoms.

## RESULTS AND DISCUSSION

### Synthesis

The syntheses of new compounds are described in **Schemes 1-4**. Suzuki Coupling of 5-bromopyridin-3-ol (**14**) with commercially available tert-butyl 4-(4-(4,4,5,5-tetramethyl-1,3,2-dioxaborolan-2-yl)-1H-pyrazol-1-yl)piperidine-1-carboxylate (**15**) provided a common intermediate (**16**) that could be used to explore substitution on the benzyl ring (**Scheme 1**). Phenol **16** was alkylated with various benzyl bromides to give the Boc-protected intermediates **17-24**, which were deprotected using trifluoroacetic acid or 4M HCl in dioxane to provide the trifluoroacetate or hydrochloride salts of final compounds **25-32**. Occasionally, materials had to be repurified and were isolated as free bases. Alternately, compound **16** could undergo a Mitsunobu reaction with (*R*) or (*S*)-1-(2-chlorophenyl)ethan-1-ol, and the resulting Boc-protected intermediates **33** & **34** could be deprotected to provide (*S*) or (*R*)-3-(1-(2-chlorophenyl)ethoxy)-5-(1-(piperidin-4-yl)-1*H*-pyrazol-4-yl)pyridine (**37** & **38** respectively). The (*S*) and (*R*) isomers of 3-(1-(4-chlorophenyl)ethoxy)-5-(1-(piperidin-4-yl)-1H-pyrazol-4-yl)pyridine (**39** & **40** respectively) were prepared in a similar fashion.

**Scheme 1:**
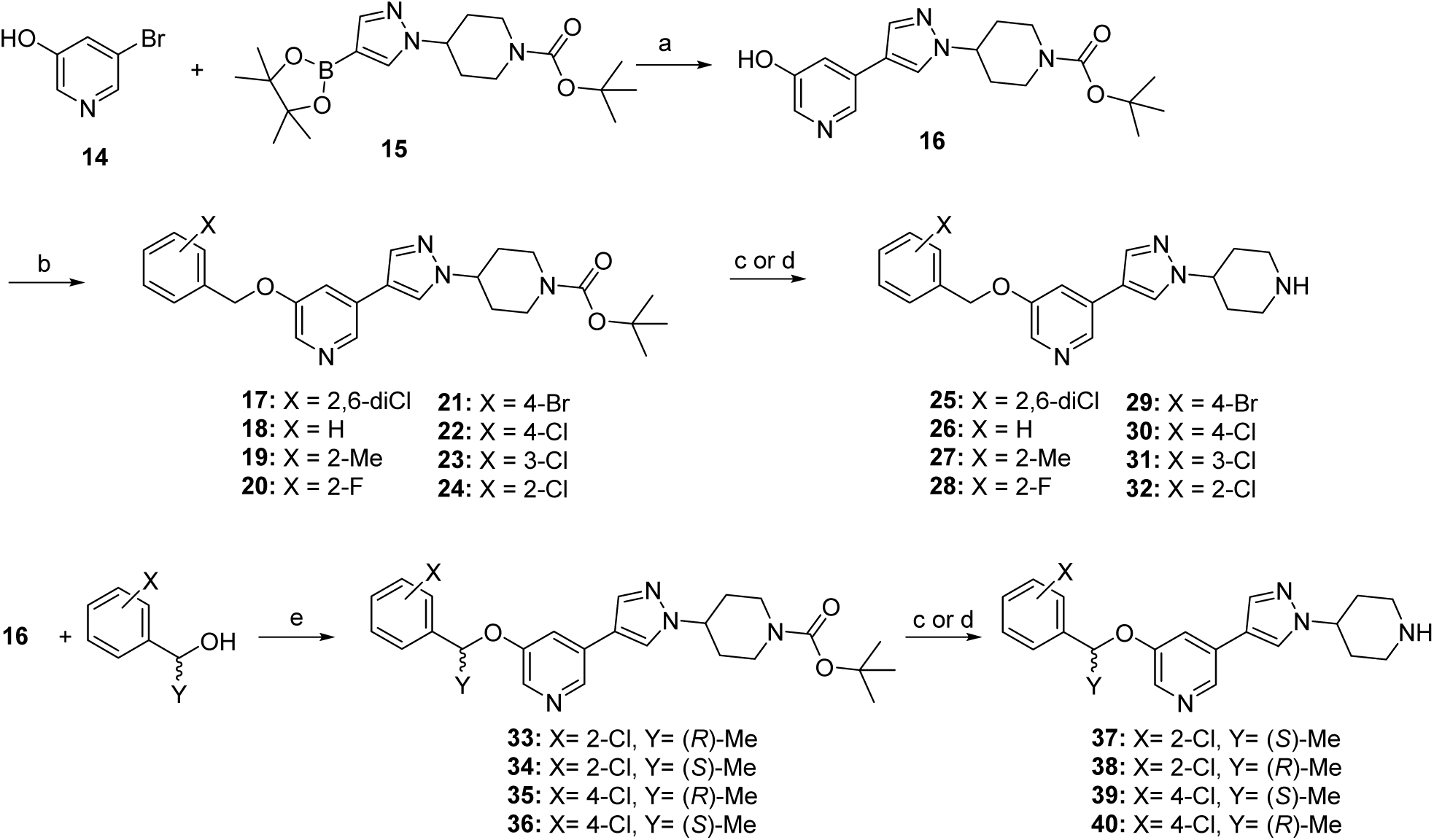
Synthesis of analogs with substitutions on the benzyl ring^a^ ^a^Reagents and Conditions: (a) Pd(dppf)Cl2, K2CO3, Toluene/EtOH/H2O, 100°C; (b) benzyl bromide, K2CO3, DMF, 100 °C; (c) 4M HCl in dioxane, CH2Cl2, 0 °C to RT; (d) TFA, CH2Cl2, 0 °C to RT; (e) Ph3P, DEAD, THF, RT.

To explore modifications on the piperidine nitrogen (**Scheme 2**), compound **24** was prepared according to **Scheme 1** and reduced with lithium aluminum hydride to provide the methylated piperidine **41**. Compound **24** was also deprotected to provide 3-((2-chlorobenzyl)oxy)-5-(1-(piperidin-4-yl)-1*H*-pyrazol-4-yl)pyridine (**32**). The nitrogen atom of the piperidine was alkylated or acylated to obtain compounds **42-46**.

**Scheme 2:**
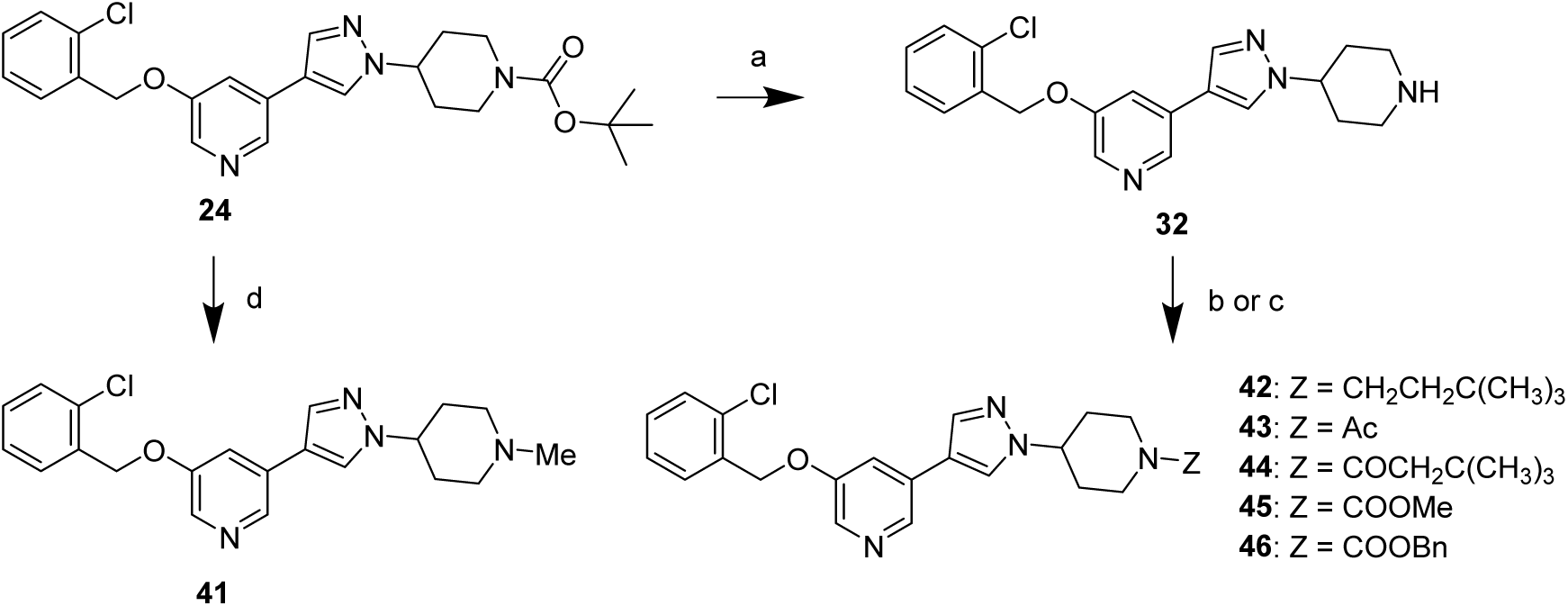
Synthesis of analogs with substitutions on the piperidine ring^a^ ^a^Reagents and Conditions: (a) 4M HCl in dioxane, CH2Cl2, 0 °C to RT; (b) 1-bromo-3,3-dimethylbutane, K2CO3, DMF, 100 °C (c) Electrophile (ROCl, RO2Cl, RCl, etc.), Et3N, CH2Cl2, 0 °C to RT; (d) LAH, THF, 0-50 °C.

To explore changes to the central pyridyl ring (**Scheme 3**), the synthesis described in **Scheme 1** was carried out using 3-bromophenol, 2-bromopyridin-4-ol, or 6-bromopyridin-2-ol without incident to furnish compounds **47**, **48** and **49**, respectively. However, while attempting to prepare compound **53** we observed exclusively N-alkylation of the pyridyl nitrogen atom of 4-bromopyridin-2-ol instead of the desired O-alkylation. Therefore, we prepared the iodo intermediate **52** from a nuclear aromatic substitution reaction of 2-(chlorophenyl)methanol (**50**) and 2-chloro-4-iodopyridine (**51**). Compound **52** was then subjected to Suzuki coupling and deprotected to afford the desired compound **53**.

**Scheme 3:**
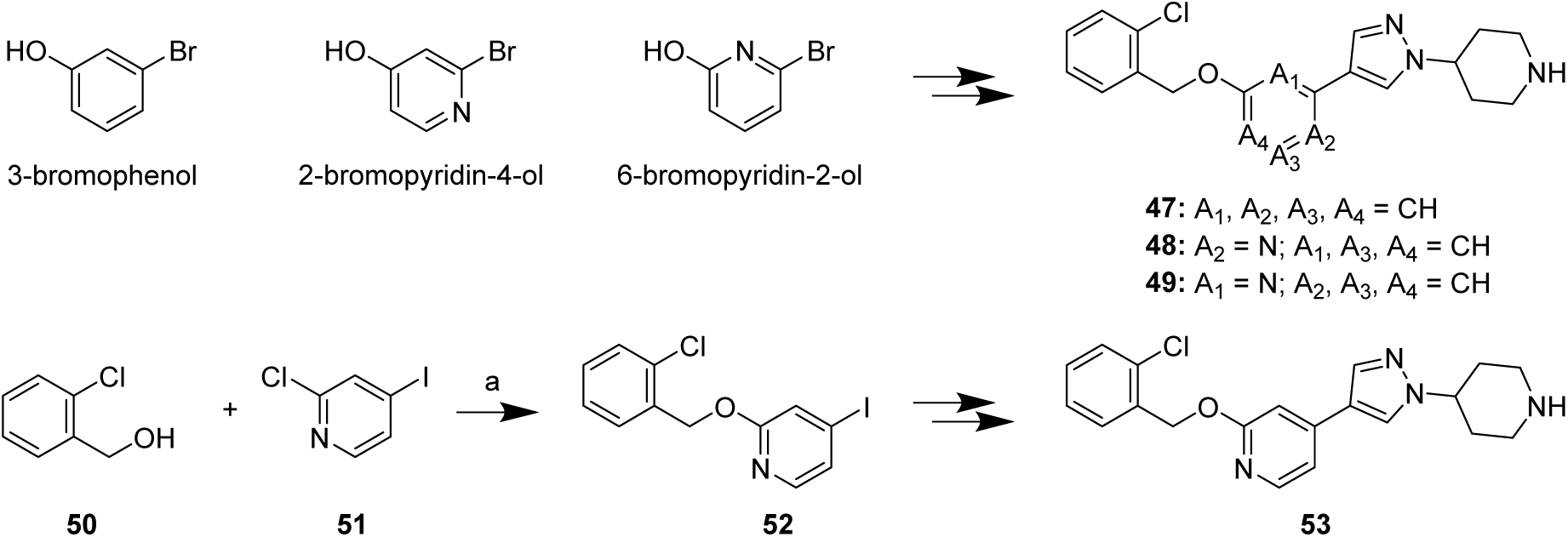
Synthesis of pyridyl ring analogs^a^ ^a^Reagents and Conditions: (a) NaH, DMF, 0 °C to RT.

We also explored modifications of the 5-membered heterocycle (**Scheme 4**). 3-Bromo-5-((2-chlorobenzyl)oxy)pyridine (**54**) was prepared by alkylation of 5-bromopyridin-3-ol (**14**) with 2-chlorobenzyl bromide. This bromide was coupled with *tert*-butyl 4-(1*H*-pyrazol-4-yl)piperidine-1-carboxylate (**55**) or *tert*-butyl 4-(1*H*-pyrazol-3-yl)piperidine-1-carboxylate (**56**), which after removal of the Boc protecting group, afforded 3-((2-chlorobenzyl)oxy)-5-(4-(piperidin-4-yl)-1*H*-pyrazol-1-yl)pyridine (**57**) and 3-((2-chlorobenzyl)oxy)-5-(4-(piperidin-4-yl)-1*H*-pyrazol-1-yl)pyridine (**58**), respectively. The alkyne **59** was prepared by a Sonogashira coupling to intermediate **54** followed by deprotection. *Tert*-butyl 4-azidopiperidine-1-carboxylate (**60**) was prepared in two steps from *tert*-butyl 4-hydroxypiperidine-1-carboxylate (**61**). A copper-catalyzed azide–alkyne cycloaddition of **59** and **60**, followed by deprotection of the Boc group yield the triazole **62**. Finally, the pyrazole regioisomer 3-((2-chlorobenzyl)oxy)-5-(1-(piperidin-4-yl)-1*H*-pyrazol-3-yl)pyridine (**63**) was prepared as described in **Scheme 1** starting from commercially available 4-(3-(4,4,5,5-tetramethyl-1,3,2-dioxaborolan-2-yl)-1*H*-pyrazol-1-yl)piperidine.

**Scheme 4:**
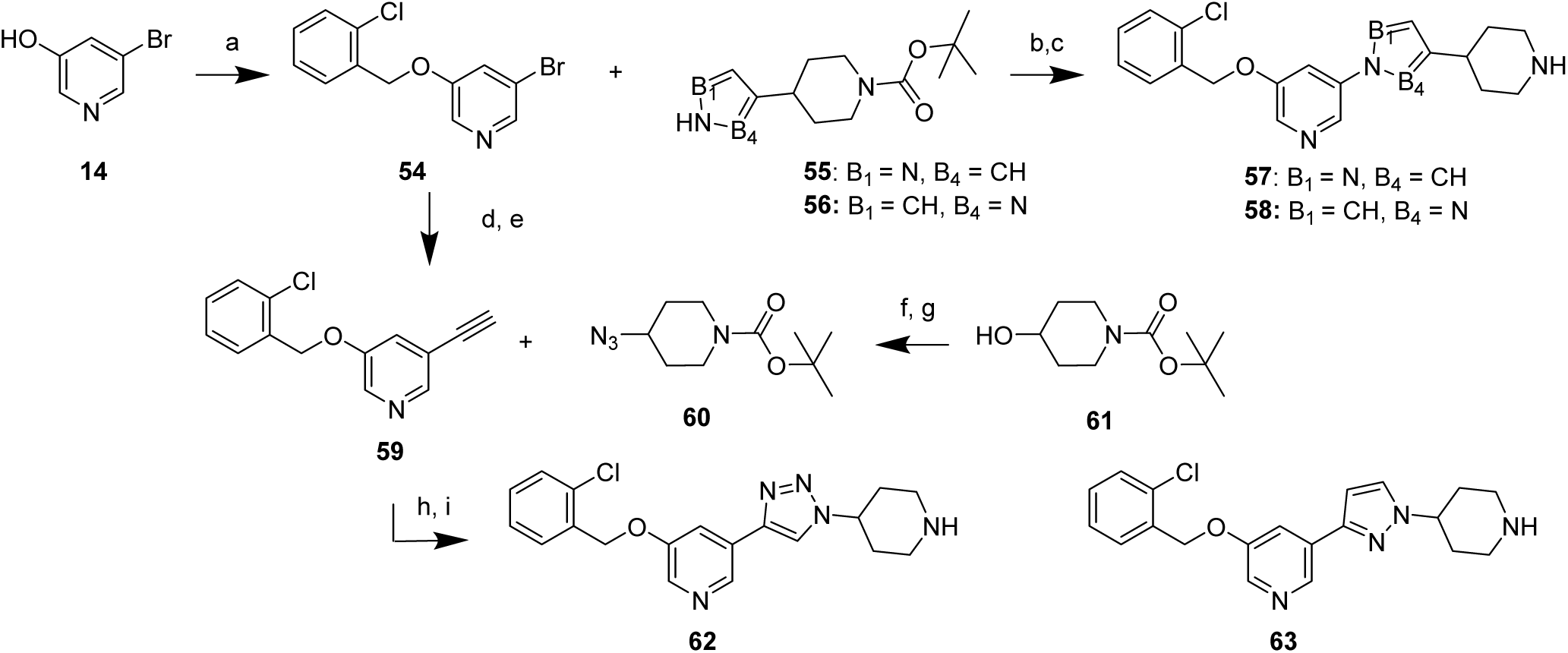
Synthesis of pyrazole ring analogs^a^ ^a^Reagents and Conditions: (a) 2-Cl benzyl bromide, Cs2CO3, DMF, RT (b) CuI, K2CO3, K3PO4, (1*R*,2*R*)-*N*^1^,*N*^2^-dimethylcyclohexane-1,2-diamine, ACN, 120 °C (c) 4M HCl in dioxane, CH2Cl2, 0 °C to RT; (d) Trimethylsilylacetylene (TMSA), Et3N, CuI, Pd(PPh3)2, 80 °C (e) K2CO3, MeOH, RT; (f) MsCl, Et3N, CH2Cl2, RT; (g) NaN3, DMF, 100 °C (h) Sodium ascorbate, CuSO4·5H2O, *n*BuOH/H2O, RT; (i) TFA, CH2Cl2, 0 °C to RT.

### Enzymatic inhibition

The SHIP1 phosphatase (Ptase) as an isolated domain was relatively inactive and unstable in our hands and as a result, we were concerned that the potency of inhibitors against a Ptase only protein may be inaccurate and fail to translate to desired cellular pharmacology. The C2 domain is known to enhance catalytic turnover^13^, and a potential allosteric binding site has been identified at the interface between the Ptase and C2 domains^46^. Therefore, we expressed multiple protein constructs containing the Ptase with its other domains, specifically the human two-domain (2D) constructs hSHIP1^395-898^ and hSHIP2^420-878^ and the human and murine five-domain (5D) hSHIP1^1-899^ and mSHIP1^1-861^ protein constructs, which were purified and used as previously reported^7^. Inhibition assays were conducted using the malachite green assay^47^. Initially, SHIP1 and SHIP2 inhibitory potencies were evaluated using the 2D (Ptase-C2) construct. We also sought to evaluate compounds against the full-length protein containing the disordered C-terminal region, but production, purification, and characterization proved difficult due to instability^7^. Therefore, we tested compounds against stable human and murine 5D (SH2+DD+PH+Ptase+C2) constructs without the disordered C-terminal domain. We evaluated compounds against the murine enzyme to test for potential species differences in anticipation of advanced studies in mice.

Enzymatic inhibitory potencies (IC_50_) were determined using soluble PI(3,4,5)P_3_-diC8 as a substrate at the K_m_ under initial velocity conditions. After a 20-minute preincubation of the 2D or 5D proteins (10 nM) with compound concentrations, the reaction was initiated by adding substrate. Reactions were quenched with Malachite BioMol Green at time points selected to ensure steady-state of product formation and adequate signal: 10 minutes for the 2D constructs and 2 minutes for the 5D constructs, which were much more active. Plates were quenched and incubated at room temperature for 30 minutes before analysis. Reaction rates were converted to percent inhibition based on enzyme controls without compound. Percent inhibition values, determined in triplicate, were plotted against compound concentration to obtain IC_50_ values. The results are summarized in **Tables 1-3**.

**Table 1.** Enzyme inhibitory activities of reference compounds^a^.

| Cmpd | Name | hSHIP1 2D<br>IC<sub>50</sub> (μM) | hSHIP2 2D<br>IC<sub>50</sub> (μM) | hSHIP1 5D<br>IC<sub>50</sub> (μM) | mSHIP1 5D<br>IC<sub>50</sub> (μM) |
| --- | --- | --- | --- | --- | --- |
| **1** | 3AC | 47.1 | 65.3 | 33.7 | 69.0 |
| **2** | K116 | 119 | 167 | 79.3 | 103 |
| **4** | K149 | 51.8 | 123 | 47.5 | 66.0 |
| **5** | AS1949490 | 52.6 | > 549 | 674 | > 952 |
| **6** | NGD-61338 | > 661 | 289 | 121 | 193 |
| **7** | CPDA | 29.7 | > 417 | > 100 | > 306 |
| **8** | SHIP2-IN-1 | 61.4 | > 230 | 82.1 | 95.6 |
| **9** |  | 279 | 266 | 191 | 256 |
| **10** | Crizotinib | 483 | 427 | 385 | 612 |
| **11** | SP1 | 9.36 | 113 | 28.0 | 35.8 |
| **12** | SP2 | 18.0 | > 303 | > 246 | >952 |
| **13** | SP3 | 52.1 | 806 | 63.4 | NT |

<sup>a</sup> Inhibitory potency values IC<sub>50</sub> (μM) are reported as the geometric mean. Number of repeats (N) > 1. NT = not tested.

Enzyme inhibitory potencies (IC_50_ values) are highly dependent on assay conditions that vary between laboratories, which confounds comparisons. To address this, our group previously measured and reported the potencies of inhibitors against both SHIP1 and SHIP2^7^. Some of these data are reproduced in **Table 1** for reference and are briefly reviewed here to highlight activity differences among scaffolds and for comparison to our SAR studies around **9**. The steroidal inhibitor 3AC (**1**), its more soluble analog K116 (**2**), and the tryptamine-based inhibitor K149 (**4**) each inhibited SHIP1 and SHIP2 in the 30–150 µM range, without distinction between the SHIP1 and SHIP2 homologs or the 2D and 5D constructs, *i.e.* the potency differences are within 3-fold, the minimum statistical ratio (MSR) of the assay^7^. AS1949490 (**5**), CPDA (**7**), and SP1-3 (**11-13**) were selective for SHIP1 over SHIP2. Interestingly, AS1949490 (**5**) and CPDA (**7**) were less effective against the 5D protein. Conversely, NGD-61338 (**6**) inhibited the 5D protein but failed to inhibit the 2D form. The recently reported SP1 (**11**) was the most potent inhibitor of the SHIP1 2D construct and also inhibited the 5D form of SHIP1. Crizotinib (**10**) exhibited weak potency, while the aminopyrimidine analog **8** was 8-fold more potent.

Although most of the reported inhibitors were active in the cell-free enzymatic assays described above, we did not observe target engagement with SHIP1 in cells, as determined by a cellular protein thermostability assay and/or they were cytotoxic^7^. However, one inhibitor, 3-((2,4-dichlorobenzyl)oxy)-5-(1-(piperidin-4-yl)-1H-pyrazol-4-yl) pyridine (**9**), engaged SHIP1 in cells, lowered pAKT levels, induced uptake of myelin by microglia, and exhibited a favorable pharmacokinetic profile with brain exposure^7^. Therefore, we executed the structure activity relationship (SAR) studies described in **Figure 3** around the pyridyl scaffold represented by compound **9**. Substitutions on the benzyl and piperidine rings were made. Modifications to the pyridyl ring changed the position of the nitrogen atom (A_1_-A_4_). Further modifications involved the pyrazole ring changing the number and positions of the nitrogen atoms (B_1_-B_5_). The enzymatic potencies of these analogs are summarized in **Tables 2**-**3**.

**Table 2.** Enzyme inhibitory activities of benzyl and piperidine ring substituted analogs of compound 9^a^.

| Cmpd | X | Y | Z | hSHIP1 2D<br>IC<sub>50</sub> (μM) | hSHIP2 2D<br>IC<sub>50</sub> (μM) | hSHIP1 5D<br>IC<sub>50</sub> (μM) | mSHIP1 5D<br>IC<sub>50</sub> (μM) |
| --- | --- | --- | --- | --- | --- | --- | --- |
| **9** | 2,4-diCl | H | H | 279 | 266 | 191 | 256 |
| **25** | 2,6-diCl | H | H | 147 | 499 | > 245 | > 572 |
| **26** | H | H | H | 196 | > 952 | 130 | 284 |
| **27** | 2-Me | H | H | > 838 | > 952 | > 840 | > 952 |
| **28** | 2-F | H | H | 695 | 391 | 908 | > 952 |
| **30** | 4-Cl | H | H | 105 | 222 | 93.5 | 121 |
| **31** | 3-Cl | H | H | 453 | 78.9 | 405 | 566 |
| **32** | 2-Cl | H | H | 384 | 683 | 177 | 379 |
| **37** | 2-Cl | (*S*)-Me | H | 344 | 196 | 294 | 578 |
| **38** | 2-Cl | (*R*)-Me | H | 238 | 74.5 | 197 | 418 |
| **39** | 4-Cl | (*S*)-Me | H | NT | NT | NT | 645 |
| **40** | 4-Cl | (*R*)-Me | H | 229 | 336 | 254 | 408 |
| **41** | 2-Cl | H | Me | > 952 | 747 | > 952 | > 952 |
| **42** | 2-Cl | H | CH<sub>2</sub>CH<sub>2</sub>C(CH<sub>3</sub>)<sub>3</sub> | 43.8 | 51.1 | 32.8 | 72.7 |
| **43** | 2-Cl | H | Ac | 233 | 165 | 156 | 405 |
| **44** | 2-Cl | H | COCH<sub>2</sub>C(CH<sub>3</sub>)<sub>3</sub> | 96.0 | > 840 | 56.4 | 102 |
| **24** | 2-Cl | H | Boc | 91.6 | > 854 | 98.2 | 50.0 |
| **45** | 2-Cl | H | COOMe | 131 | > 326 | 87.9 | 102 |
| **46** | 2-Cl | H | COOBn | 27.9 | > 109 | 41.9 | 29.0 |

<sup>a</sup> Substitutions X on the benzyl ring, Y at the α-carbon, and Z on piperidine ring nitrogen atom (**Figure 3**). Inhibitory potency values IC<sub>50</sub> (μM) are reported as the geometric mean. Number of repeats (N) > 1. NT = not tested.

**Table 3.**
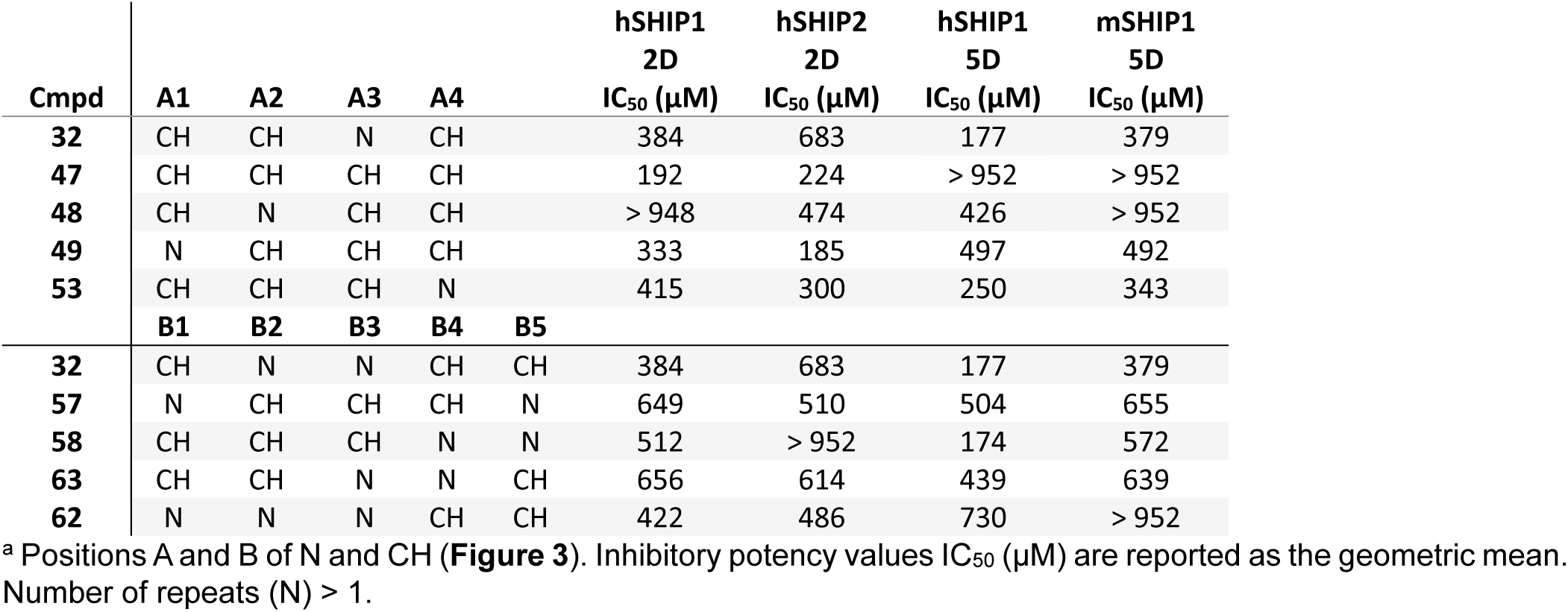
Enzyme inhibitory activities of pyridyl ring analogs of compound 32^a^.

Substitutions around the benzyl ring were generally well tolerated, including modifications at the alpha carbon (**Table 2**). A preference for 4-chloro substitution (**30**) was observed, along with a slight preference for the (*R*)-enantiomer of a methyl substitution at the alpha carbon (**37** vs. **38**). Notably, the 2-methyl (**27**) and 2-fluoro (**28**) substitutions led to a surprising reduction in potency compared to the unsubstituted analog (**26**), which retained activity. Modifications to the piperidine ring had a more pronounced impact. Capping the nitrogen with a methyl group (**41**) completely abolished activity while increasing the size of an alkyl substitution improved potency up to sixfold, as observed with compound **42**, which contains the highly hydrophobic 2,2-diimethylbutyl group. Capping the nitrogen with an acetyl group (**43**) preserved activity. In a similar fashion to alkyl substitution, further increasing hydrophobic substitution size enhanced potency (**44**). Among the modifications, carbamates (**24** & **45**) proved most effective, with the benzyl carbamate (**46**) emerging as the most potent inhibitor of this series in terms of enzymatic inhibition demonstrating that this scaffold can achieve potencies comparable to 3AC (**1**) while improving selectivity over SHIP2.

Changes to the pyridyl and pyrazole rings shown in **Tables 3** were tolerated in the enzyme assays in most cases. Since 3-((2-chlorobenzyl)oxy)-5-(1-(piperidin-4-yl)-1H-pyrazol-4-yl) pyridine (**32**) emerged as one of the more effective compounds in cellular assays, these analogs were prepared as matched pairs to this compound. Removing the nitrogen of the central pyridine ring to give compound **47** retained activity at the 2D protein but this analog failed to inhibit the 5D. Moving the substituents around the pyridine ring to the 4- and 2-positions (**48**) dramatically reduced activity, while moving these substituents to the 2- and 6-positions (**49**) retained activity. We also prepared the 2-, 4-substituted analog **53** and it retained activity. Moving the pyridyl and piperidine attachments around the pyrazole ring (**32** vs **57**, **58** & **63**) slightly reduced activity by only 2-fold while the triazole analog **62** retained activity. Overall, these SAR studies indicate that some variations in the position of the nitrogen atom within the pyridyl core are not tolerated while positional changes of nitrogens in the pyrazole had minimal impact on potency in cell-free enzyme assays.

### Cellular target engagement

We developed two complementary cellular assays to assess target engagement in a whole cell context. First, we implemented a Bioluminescence-based Thermal Shift Assay (BiTSA) as a measurement of SHIP1 thermal stability that can be modulated by the direct binding of compounds to the protein in intact cells. Second, we established a phospho-AKT to total AKT ratio (pAKT/tAKT) assay to evaluate the functional consequences of SHIP1 modulation on proximal downstream signaling.

The Cellular Thermal Shift Assay (CETSA^®^) has become well-established method for assessing ligand-induced protein thermostability changes^48^. We implemented a split NanoLuciferase format to assess SHIP1 thermostability in a cell-based assay with sufficient throughput^49-51^ for the SAR studies described in **Tables 4**-**6**. HMC3 cells stably overexpressing HiBiT-*INPP5D* were used to assess the thermal stability of full-length SHIP1 in two modes. In screening mode, cells were treated with 100 μM of each compound at 37 °C for 60 min, followed by a 3-min isothermal heating at 44.2 °C, the experimentally determined Tm (n = 28). In concentration-response mode, cells were treated with a serial dilution of compound, followed by heating, and then the luminescence measured and reported as a percentage of DMSO control. An AC_50_, the concentration required to induce a 50% change in the thermal stability of HiBiT-SHIP1, was calculated using a four-parameter logistic regression model if the compound reduced luminescence by at least three times the standard deviation (>3SD) of the Tm compared to control. Otherwise, an AC_50_ was not calculated, and the compound was reported as inactive. To confirm that observed protein thermal stability changes were not due to inhibition of complimenting with LgBiT to reconstitute the luminescent NanoBit enzyme or it’s luminescence activity, an assay was performed in parallel using the HiBiT control protein (N3010, Promega).

**Table 4.** SHIP1 cellular target engagement (BiTSA) and downstream signaling (pAKT/tAKT) modulation by reference compounds.

| Cmpd | Name | BiTSA |  | pAKT/tAKT (S473) | pAKT/tAKT (T308) |
| --- | --- | --- | --- | --- | --- |
|  |  | AC<sub>50</sub> (μM)<sup>a</sup> | % remain<sup>b</sup> | IC<sub>50</sub> (μM)<sup>c</sup> | IC<sub>50</sub> (μM)<sup>c</sup> |
| 1 | 3AC (HCl) | NC | 98 | > 44.7 | > 100 |
| 2 | K116 | 94.3 | 34 | 20.8 | 22.8 |
| 4 | K149 | NT | 104* | > 100 | > 100 |
| 5 | AS1949490 | NC | 84 | > 100 | > 100 |
| 6 | NGD-61338 | NT | 89* | NT | NT |
| 7 | CPDA | NC | 92 | > 100 | > 50 |
| 8 | SHIP2-IN-1 | NC | 92 | > 40 | >36.0 |
| 9 |  | 68.2 | 29 | 13.1 | 18.1 |
| 10 | Crizotinib | 23.2 | 13 | NT | 2.5 |
| 11 | SP1 | NC | 92* | NT | NT |

<sup>a</sup> Concentration (μM) that induced a half-maximum loss of luminescence (AC<sub>50</sub>) compared to control. Number of repeats (N) > 1. NT = not tested.
<sup>b</sup> Minimum percent luminescence at any dose in the concentration-response or from the single point screen at 100 μM (\*). Target engagement was considered significant if ΔT<sub>m</sub> difference from control was >3SD, otherwise AC<sub>50</sub> was not calculated (NC) and the compound was reported as inactive.
<sup>c</sup> Reduction in the pAKT/tAKT ratio reported as the geometric mean of the IC<sub>50</sub> (μM). Number of repeats (N) = 2 biological replicates each tested in duplicate. NT = not tested.

Given the role of SHIP1 regulating phosphatidylinositol signaling, ligands that engage this protein would be expected to modulate the AKT pathway (**Figure 1**). Therefore, to provide additional evidence for on-target activity and assess SAR, we implemented an AKT signaling assay in THP-1 cells. Western blot analysis confirmed SHIP1 expression in THP-1 cells (Supplementary **Figure S1**). IL-4 treatment of THP-1 cells upregulates the low-affinity inhibitory FcγRIIB receptor, producing a cellular context in which FcγRIIB can recruit SHIP1 to regulate PI3K/Akt signaling^52, 53^. To evaluate SAR, THP-1 cells were conditioned with IL-4 for 24 hours followed by 24-hour recovery and were then treated for 15 minutes with a serial dilution of each compound. pAKT and tAKT levels were quantified using the Revity Alpha SureFire Ultra Multiplex PhosphoAKT (S473 or T308) assay kit. The pAKT/tAKT ratios were calculated from the average of two technical replicates per wavelength and the ratio used to generate IC_50_ values. To ensure that changes in AKT signaling were not attributable to cytotoxic effects, cellular viability was assessed using the CytoTox-Glo^®^ luminescent assay. A summary of BiTSA and pAKT/tAKT ratio data is provided in **Tables 4**-**6**.

Although the HCl salt of 3AC (**1**) was inactive, the more soluble analog K116 (**2**) did show activity in both assays suggesting this scaffold does engage SHIP1 (**Table 4**). None of the other previously reported SHIP inhibitors were active in these assays except crizotinib and compound **9**. Because the pyridyl scaffold represented by compound **9** successfully engaged SHIP1 as determined by the BiTSA and reduced pAKT levels, we pursued SAR studies around this scaffold (**Tables 5**-**6**). None of these compounds showed appreciable activity against the HiBiT control protein or in the CytoTox-Glo toxicity assay performed in THP-1 cells under comparable conditions (data not shown). Similar to the SAR trends observed in the enzyme assays, substitutions around the benzyl ring were generally well tolerated, including modifications at the alpha carbon (**Table 5**). The 2-, 3-, and 4-chloro substitutions (**32**, **31**, **30**) were preferred in that order compared to the unsubstituted (**26**), 2-fluoro (**28**), and 2-methyl (**27**) substituted analogs with the 2-chloro analog **32** being the most potent. Similar activities were observed when a methyl group was added to the alpha carbon of the benzyl (**32** vs. **37**/**38** and **30** vs **39**/**40**) with a slight preference for the *S*-methyl enantiomers **37** and **39**. Substitution on the piperidine ring nitrogen typically reduced activity in these assays in stark contrast to enzyme assays for which these changes, especially increasing substitution size, increased activity. In fact, most of these compounds (**24**, **42-46**) failed to meet the % remaining threshold for AC_50_ calculation, and those that did were only weakly active with none more active than compound **9**, the starting point for these studies. None of these compounds were active in the pAKT assay except the methyl substituted compound **41**, which was substantially less potent than compound **9**. Changes to the pyridyl and pyrazole rings are shown in **Tables 6**. Removing the nitrogen of the central pyridine ring to give compound **47** reduced activity as did moving the substituents around the pyridine ring, revealing a preference for compound **32** with the pyridyl ring substituted at the 3- and 5-positions. In contrast, moving substitutions around the pyrazole ring was tolerated with only a small preference for compound **32**.

**Table 5.** SHIP1 cellular target engagement (BiTSA) and downstream signaling (pAKT/tAKT) modulation by benzyl and piperidine ring substituted analogs of compound 9^a^.

| Cmpd | X | Y | Z | BiTSA |  | pAKT/tAKT<br>(S473) | pAKT/tAKT<br>(T308) |
| --- | --- | --- | --- | --- | --- | --- | --- |
|  |  |  |  | AC<sub>50</sub> (μM)<sup>b</sup> | % remain<sup>c</sup> | IC<sub>50</sub> (μM)<sup>d</sup> | IC<sub>50</sub> (μM)<sup>d</sup> |
| **9** | 2,4-diCl | H | H | 68.2 | 29 | 13.1 | 18.1 |
| **25** | 2,6-diCl | H | H | 50.5 | 13 | > 28.5 | >38.7 |
| **26** | H | H | H | 101 | 37 | 10.0 | 12.7 |
| **27** | 2-Me | H | H | 100 | 38 | 13.3 | 11.5 |
| **28** | 2-F | H | H | 139 | 68 | 11.6 | 14.8 |
| **29** | 4-Br | H | H | 67.6 | 28 | NT | NT |
| **30** | 4-Cl | H | H | 56.5 | 19 | 10.1 | 9.4 |
| **31** | 3-Cl | H | H | 61 | 33 | 6.9 | 8.2 |
| **32** | 2-Cl | H | H | 40.4 | 25 | 4.0 | 5.3 |
| **37** | 2-Cl | (S)-Me | H | 37.4 | 24 | 9.8 | 7.5 |
| **38** | 2-Cl | (R)-Me | H | 31.4 | 23 | 15.7 | 16.2 |
| **39** | 4-Cl | (S)-Me | H | 41.0 | 22 | 8.0 | 7.5 |
| **40** | 4-Cl | (R)-Me | H | 46.5 | 19 | 14.2 | 12.6 |
| **41** | 2-Cl | H | Me | > 92.7 | 62 | >26.5 | 23.2 |
| **42** | 2-Cl | H | CH<sub>2</sub>CH<sub>2</sub>C(CH<sub>3</sub>)<sub>3</sub> | NC | 72 | > 100 | > 100 |
| **43** | 2-Cl | H | Ac | 236 | 87 | > 25 | > 50 |
| **44** | 2-Cl | H | COCH<sub>2</sub>C(CH<sub>3</sub>)<sub>3</sub> | > 136 | 69 | > 50 | > 100 |
| **24** | 2-Cl | H | Boc | NC | 67 | > 100 | > 100 |
| **45** | 2-Cl | H | COOMe | > 90.3 | 75 | > 100 | > 100 |
| **46** | 2-Cl | H | COOBn | NC | 80 | > 100 | > 100 |

<sup>a</sup> Substitutions X on the benzyl ring, Y at the α-carbon, and Z on piperidine ring nitrogen atom (**Figure 3**).
355 <sup>b</sup> Concentration (μM) that induced a half-maximum loss of luminescence (AC<sub>50</sub>) compared to control. Number of repeats (N) > 1. NT = not tested.
<sup>c</sup> Minimum percent luminescence at any dose in the concentration-response. Target engagement was considered significant if ΔT<sub>m</sub> difference from control was >3SD, otherwise AC<sub>50</sub> was not calculated (NC) and the compound was reported as inactive.
360 <sup>d</sup> Reduction in the pAKT/tAKT ratio reported as the geometric mean of the IC<sub>50</sub> (μM). Number of repeats (N) = 2 biological replicates each tested in duplicate. NT = not tested.

**Table 6.**
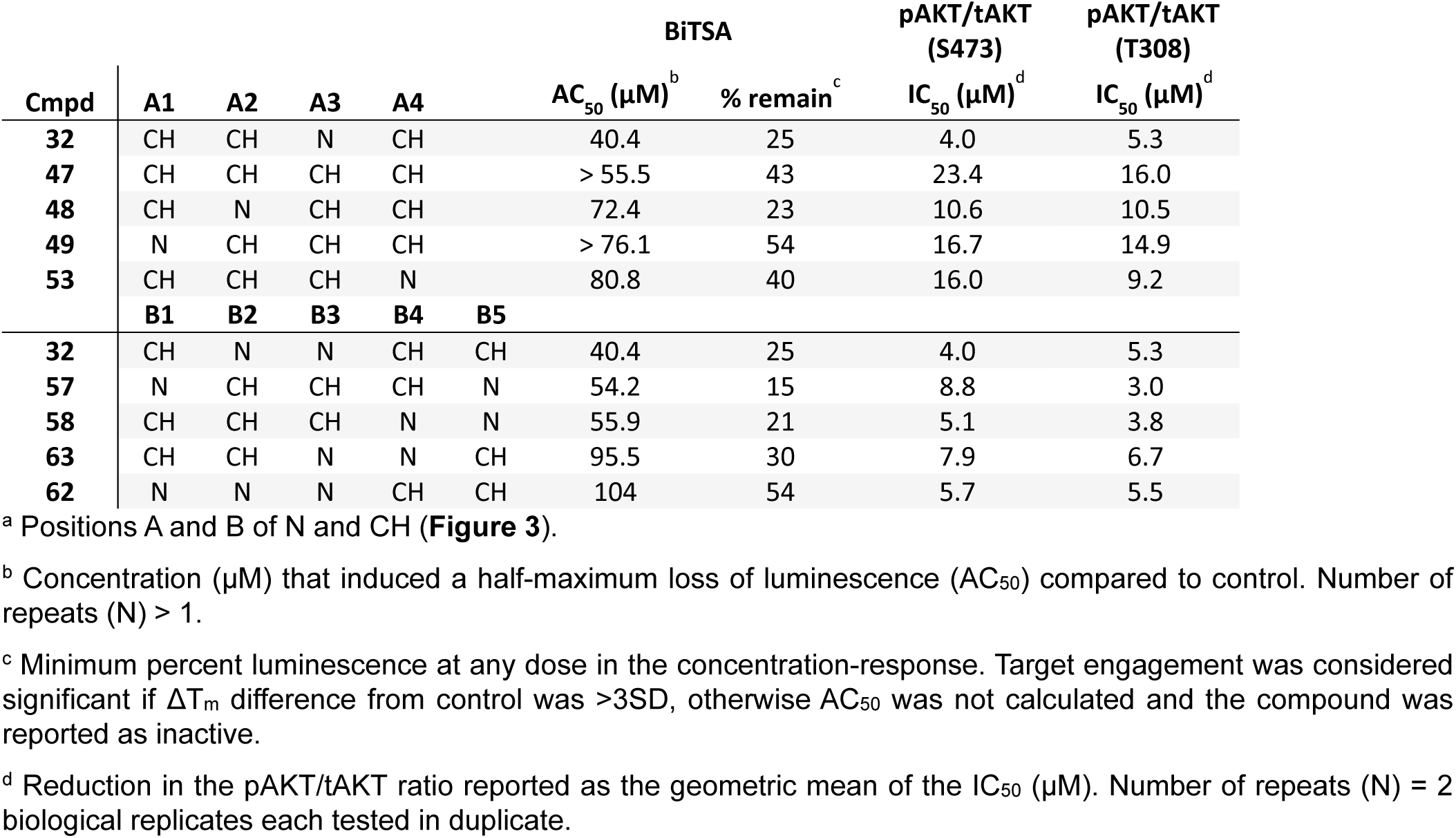
SHIP1 cellular target engagement (BiTSA) and downstream signaling (pAKT/tAKT) modulation by pyridyl ring analogs of compound 32^a^.

### Lipidomic profiling

The phosphoinositides (PIPs) are low-abundant lipids in cell membranes that influence cellular processes such as endocytosis, exocytosis, and organelle maturation. Their levels are tightly regulated by a complex and dynamic metabolic network governed by lipid metabolism and the coordinated exchange of lipids between membranes^54, 55^. Because SHIP1 is known to bind PIPs that are phosphorylated at the 3-position, and primarily converts PI(3,4,5)P_3_ to PI(3,4)P_2_, target engagement would be expected to shift the overall pool of PIP species in a cell. Having established target engagement of the HiBiT-SHIP1 protein in HMC3 cells and modulation of AKT signaling in THP-1 cells, we next examined the effect of a subset of compounds on the overall phosphoinositide pool in THP-1 cells. We reasoned that if these compounds modulate SHIP1 activity, corresponding changes in phosphatidylinositol (PI) and phosphatidylinositol phosphate (PIP) levels would occur in a manner consistent with activities in the BiTSA and pAKT/tAKT assay. Therefore, we quantified the levels of PI and PIP species in THP-1 cells expressing endogenous levels of SHIP1. In this study, we did not expect to directly observe changes in the levels of PI(3,4,5)P_3_ and PI(3,4)P_2_ because of the very low abundance of these PIPs. In most mammalian cells including monocytes, the dominant molecular species is PI 38:4 with stearoyl and arachidonoyl side chains (18:0/20:4). We compared the ability of compound **32** to shift the overall PI and PIP pool with two analogs substituted on the piperidine ring, the acetamide **43**, which failed to demonstrate target engagement as assessed by the BiTSA and pAKT/tAKT assay, and the methyl substituted analog **41**, which lost substantial activity. The experiment was performed in serum-free RPMI media using THP-1 cells treated with compound (n = 3 per condition at 2, 10, 20 and 40 µM) compared to vehicle control (0.2% DMSO, n = 2), each with a final DMSO concentration of 0.2% across all conditions. Cells were incubated at 37 °C for 5 hours before lipid extraction and analysis using mass-spectrometry–based lipidomics. Phosphatidylinositol phosphate signals were combined (PIP1 and PIP2 combined separately), and data normalized to total PI. Compound **32** increased the overall levels of PIP1 species and decreased the levels of PIP2, consistent with target engagement as assessed by the BiTSA and pAKT/tAKT assay (**Figures 4A-D**). Compounds **43** and **41** were significantly less active as expected. Intriguingly, the effect of compound **32** on phosphatidylinositols was dependent on the acyl chains of the detected PI species (Supplemental **Figure S2**), suggesting that membrane composition may influence the activity of the compound and/or SHIP1 itself, as has been observed with diacylglycerols (DAGs), where different acyl chain compositions influence the activity of protein kinase C family proteins and downstream signaling^56, 57^. Changes in PIP2 levels were observed to a lesser extent due to their lower abundance and the dynamic flux between different PIP2 species to the more abundant PIP1. Changes in PIP3 were not detected because of very low abundance. These results demonstrate that compound **32** modulates phosphatidylinositol metabolism in a concentration range consistent with cellular activity and the inactive and less active analogs, compounds **43** and **41**, have no and lower propensity to shift the overall phosphoinositide pool, respectively.

**Figure 4.**
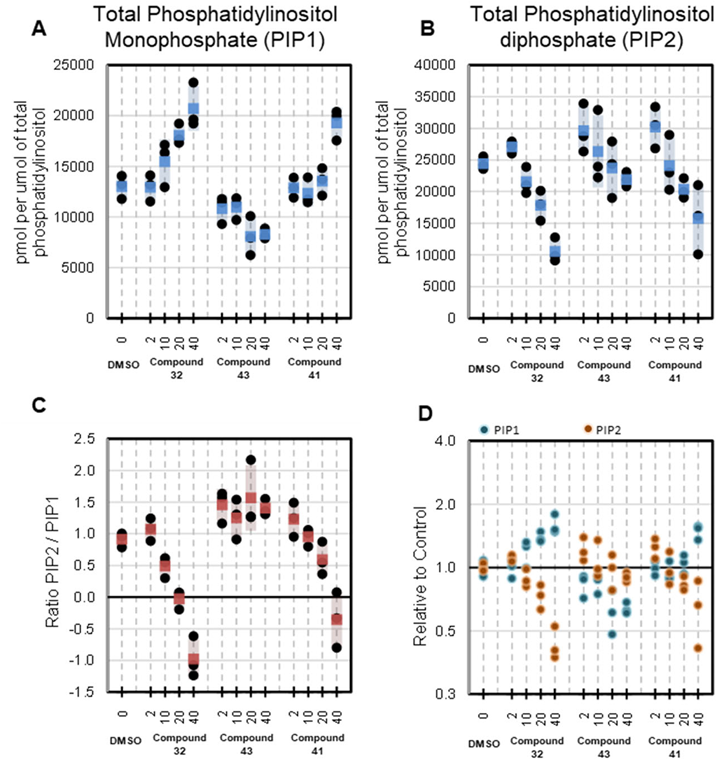
Quantification of Phosphoinositide Lipid Species Following Compound Treatment in THP-1 Cells. THP-1 cells were cultured in serum-free RPMI media and treated with compounds **32**, **43** and **41** at 2, 10, 20, and 40 µM (n = 3 per condition) including a vehicle control (0.2% DMSO, n = 3) with all conditions having a final DMSO concentration of 0.2%. Cells were incubated at room temperature for 5 hours prior to lipid extraction and analysis by mass-spectrometry-based targeted lipidomics. **A–B.** Total levels of mono- (PIP1) and bis- (PIP2) phosphoinositides (pmol), normalized to total phosphatidylinositol (PI). **C.** Ratio log2 transformed of PIP2 to PIP1 following compound treatment. **D.** Relative abundance of PIP1 and PIP2 compared to DMSO control with individual replicates for each compound treatment.

### Protein Lipid Overlay (PLO) assay

Because we observed shifts in the levels of PI, PIP1 and PIP2 in THP-1 cells, we tested the activity of compound **32** in a protein-lipid overlay (PLO) assay to assess its ability to alter SHIP1 interactions with phosphoinositides on a membrane surface. The PLO assay is a semi-quantitative technique used to evaluate protein-lipid associations and to assess the relative binding preference of a protein for various lipids^58, 59^. A nitrocellulose membrane pre-spotted with a concentration gradient of phosphatidylinositol (PI) and all 7 phosphoinositides (PIPs) was purchased from Echelon Biosciences (PIP Array P-6100). Following an initial blocking step with 5% fatty acid–free BSA to inhibit nonspecific binding, the membrane was incubated with the 5D SHIP1 protein in the presence or absence of compound **32** at 1.0, 3.3, and 10 µM. After incubation, unbound protein was removed by washing and to detect bound SHIP1, the membrane was incubated with a primary antibody (Invitrogen MA1-10450) that recognizes the N-terminal SH2 domain of SHIP1, followed by incubation with a secondary antibody (Li-Cor 926-32210) for visualization. The membrane was imaged using chemiluminescence (800 nm) to assess protein-lipid interaction profiles across the membrane. The results are shown in **Figure 5**. The 5D SHIP1 protein had no association with unphosphorylated PI and showed a clear preference for di- and tri-phosphorylated phosphoinositides over the monophosphorylated forms, exhibiting a relative overall preference for PI(4,5)P_2_. Compound **32** distinctly altered lipid association preferences among the various phosphoinositides at concentrations between 1.0 and 3.3 µM, shifting the protein’s preference away from monophosphorylated PIPs toward di- and tri-phosphorylated forms. Notably, the relative association with the product of the phosphatase activity of SHIP1, PI(3,4)P_2_, increased dramatically, suggesting a possible mode of action.

**Figure 5.**
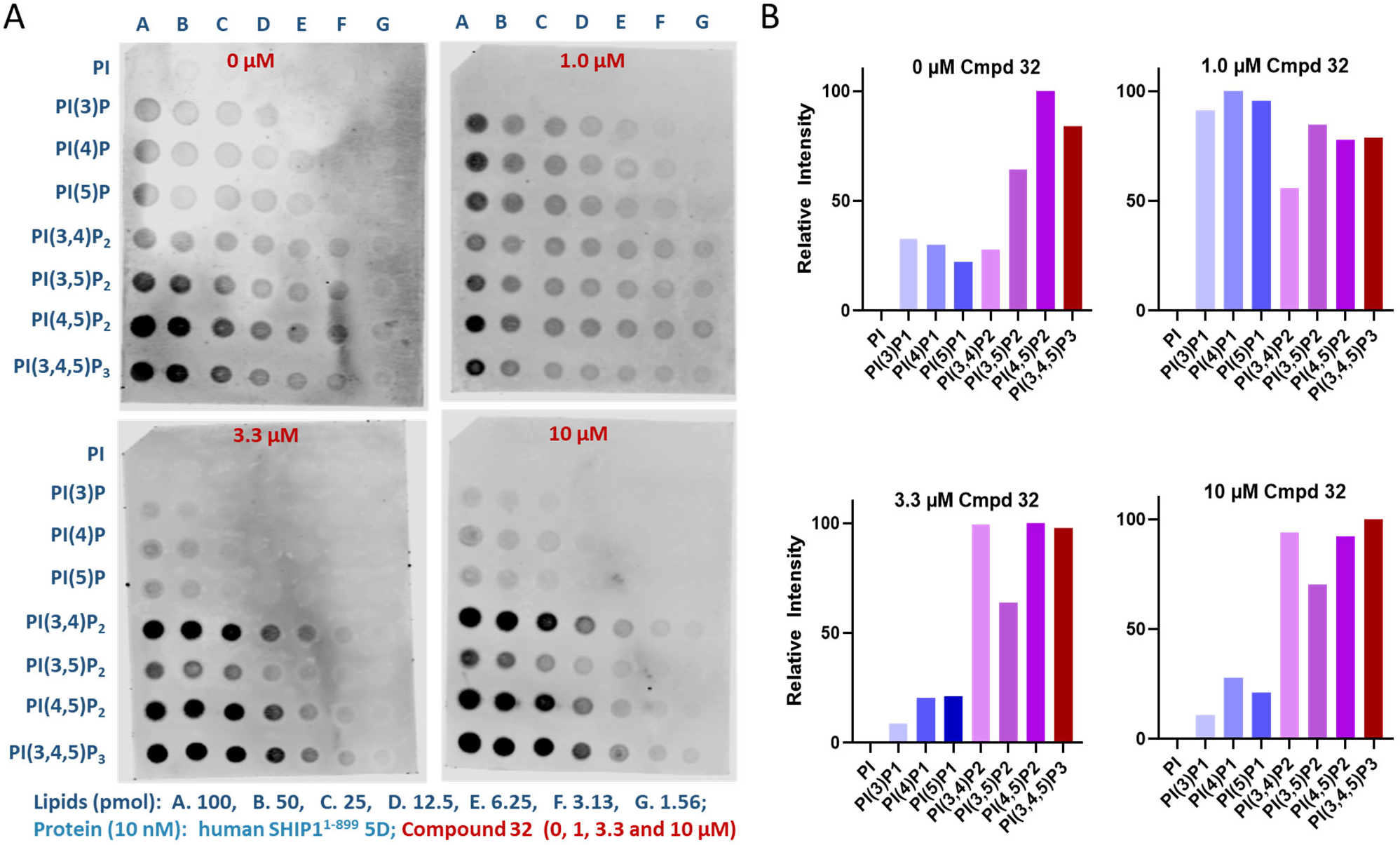
Protein Lipid Overlay (PLO) Assay. **A.** Pre-spotted nitrocellulose membranes containing a panel of phospholipids (Echelon Biosciences) were used to assess interactions with the human SHIP1^1-899^ five domain protein in the presence of increasing concentrations of compound **32**. SHIP1 5D protein (10 nM) was incubated with membranes containing a serial array of all eight phosphoinositides. Each membrane is spotted with descending quantities (100 to 1.56 pmol, columns A–G) of PI, PI(3)P, PI(4)P, PI(5)P, PI(3,4)P₂, PI(3,5)P₂, PI(4,5)P₂, and PI(3,4,5)P₃. Association of the protein was detected using an antibody against SHIP1 (Invitrogen MA1-10450) followed by incubation with a secondary antibody (Li-Cor 926-32210) for visualization. **B.** Intensities of the 100 pmol spots were quantified using ImageJ and normalized to the maximum value arbitrarily set to 100% on each membrane to generate Relative Intensity values between spots.

### Cellular pharmacology

Because our animal models of desired pharmacology are murine, we sought to confirm SHIP1 expression in BV2 cells, a reliable and reproducible murine cellular model for studying microglial function due to their high rate of proliferation and consistency compared to primary murine microglial cultures. Previous testing in the enzyme assays described above indicated that the pyridyl scaffold inhibits both the human and murine 5D SHIP1 proteins to a similar degree. Microglia isolated from murine brain were used to verify that results using BV2 cells are similar to those obtained with primary cells. Microglia were isolated as described previously^60-62^. Briefly, cortical and hippocampal tissue from neonate C57BL/6J mice (P0-P3) was obtained, dissociated it into a single-cell suspension, and cultured. Western blot analysis confirmed SHIP1 expression in BV2 cells and wild type primary microglia (Supplementary **Figure S3A**). Mice lacking the promoter and exon 1 of the *Inpp5d* gene (B6.129S6(C)-*Inpp5d*^tm1.1Wgk^/J; *Inpp5d*^HET^) were crossed to generate *Inpp5d^HET^*, *Inpp5d^HOM^* and *Inpp5d^WT^* mice. Primary microglia were isolated from *Inpp5d^HET^*and *Inpp5d^HOM^* mice (P0-P3) to obtain cells with reduced or ablated *Inpp5d* expression. Western blot analysis confirmed the expected reduction and loss of SHIP1 expression in these cells (Supplementary **Figure S3B**).

Given that SHIP1 is a negative regulator of TREM2, a key microglial receptor responsible for sensing cellular debris, especially lipids associated with misfolded proteins such as amyloid, we employed a pH-sensitive fluorescent probe, pHrodo, to assess microglial uptake of myelin/membrane debris. This approach enabled live-cell imaging and quantification of membrane debris uptake while simultaneously assessing cell count and nuclear intensity in a high-content imaging assay, providing an assessment of both desired cellular pharmacology and cell health. Myelin/membrane debris was isolated from mouse brain and labeled with pHrodo as previously reported^63^. BV2 cells were treated with test compounds for 24 hours, followed by the addition of pHrodo-myelin/membrane debris. After an additional 20-hour incubation, nuclear staining solution (Hoechst 33342) was added, and plates were incubated for 30 minutes before imaging with an ArrayScan XTI high-content analysis reader (Thermo Scientific). Since this assay follows a mix-and-read format without media exchange, an imaging-based cell count is reliable. Image data were subsequently analyzed using Thermo Scientific HCS Studio. Assay readouts included mean total spot intensity per cell to assess uptake of pHrodo-myelin/membrane debris, total cell count per well as a measure of inhibition of proliferation or cytotoxicity and mean average nuclear intensity per cell as another indicator of cell health. Compound potencies for each endpoint were determined using a four-parameter logistic curve regression model to derive EC_50_ values for uptake of myelin/membrane debris and IC_50_ values for cell count and nuclear intensity. An increase in pHrodo-myelin/membrane debris uptake is interpreted as microglial activation, whereas a decrease in cell count with nuclear intensity changes indicates cytotoxicity, either due to lower cell proliferation or increased cell death. Concentration response curves for all compounds that demonstrated target engagement as well as or better than compound **9** are shown in Supplemental **Figure S4A** including those that appear to be cytotoxic. In addition, any compound that increased pHrodo-myelin/membrane debris uptake at any concentration without reduced cell count or nuclear intensity are included regardless of target engagement. For a subset of compounds, we also assessed their ability to induce the uptake of pHrodo-myelin/membrane debris by wildtype primary murine microglia (Supplemental **Figure S4B**), which were isolated as described above. The EC_50_ and IC_50_ values of the results with these compounds are shown in **Table 7**.

**Table 7.** High content imaging assay results: uptake of pHrodo-myelin/membrane debris and cellular health^a^.

| Cmpd | BV2 |  |  | Primary Murine Microglia |  |  |
| --- | --- | --- | --- | --- | --- | --- |
|  | Cell Count | Nuclear Intensity | Uptake | Cell Count | Nuclear Intensity | Uptake |
|  | IC<sub>50</sub> (μM) | IC<sub>50</sub> (μM) | EC<sub>50</sub> (μM) | IC<sub>50</sub> (μM) | IC<sub>50</sub> (μM) | EC<sub>50</sub> (μM) |
| **2** | 1.00 | 1.59 | NC | 8.71 | 8.72 | NC |
| **10** | 0.769 | 1.17 | NC | NT | NT | NT |
| **9** | 2.52 | 2.95 | 0.260 | 2.12 | 7.03 | NC |
| **32** | 3.45 | 6.46 | 0.777 | 5.65 | 19.4 | 0.484 |
| **31** | 2.59 | 5.61 | NC | 3.78 | 2.02 | NC |
| **30** | 2.53 | 6.61 | 0.750 | 2.72 | 2.29 | NC |
| **25** | 7.24 | 1.71 | NC | 8.53 | 18.5 | NC |
| **26** | 7.41 | 30 | 21.00 | 7.48 | 26.3 | 4.09 |
| **37** | 6.60 | 7.33 | NC | 6.89 | 21.5 | NC |
| **38** | 5.52 | 4.97 | NC | 5.72 | 9.90 | 25.6 |
| **39** | 2.91 | 16.0 | NC | NT | NT | NT |
| **40** | 2.69 | 6.90 | 0.742 | 6.16 | 7.42 | NC |
| **44** | 19.1 | 4.99 | 2.46 | 55.5 | 59.1 | 4.94 |
| **45** | 57.3 | 99.7 | 19.1 | > 60 | 14.5 | 5.65 |
| **46** | > 60 | > 60 | 0.943 | 26.15 | >60 | 0.38 |
| **57** | 4.06 | 6.96 | NC | 2.39 | 2.40 | NC |
| **58** | 5.22 | 16.4 | NC | 7.14 | 6.93 | NC |

<sup>a</sup> Values reported as the geometric mean of the IC<sub>50</sub> or EC<sub>50</sub> (μM). Number of repeats (N) > 1. NC = not calculated. NT = not tested.

Previously reported SHIP inhibitors that engaged the target as assessed by the BiTSA and pAKT/tAKT assay were K116 (**2**), crizotinib (**10**), and compound **9**. In the pHrodo-myelin/membrane debris uptake assay described above, K116 (**2**) proved to be cytotoxic to both BV2 cells and primary murine microglia. Crizotinib (**10**) was potently cytotoxic to BV2 cells, which have a rapid proliferation rate, consistent with its activity against aurora kinase B (*see below*), and was therefore not tested in primary murine microglia. Fortunately, compound **9** promoted the uptake of pHrodo-myelin/membrane debris at concentrations that did not lead to reduced cell count or nuclear intensity changes. In contrast to the SAR studies describe above in the biochemical and cellular target engagement assays, changes to compound **9** were not well tolerated. The most notable improvement involved simply removing the 4-chloro substituent to give the 2-chloro substituted analog **32**, which markedly improved uptake at concentrations without cytotoxicity. The 2-chloro substitution (**32**) was also preferred over the 3-chloro (**31**), the 4-chloro (**30**), and the 2,6-di-chloro (**25**) substituted analogs as well as the unsubstituted analog **26**. Methyl substitution at the alpha carbon (compounds **37, 38, 39** and **40**) reduced activity. Substitution on the piperidine nitrogen was allowed in some cases, most notably compound **44**, but the increased myelin uptake induced by compounds **45** and **46** was dramatic and not well separated from cytotoxicity suggesting an off-target mechanism, which is consistent with their lack of target engagement as assessed by the BiTSA and pAKT/tAKT assay and in stark contrast to the biochemical SAR studies described above. Moving substituents around either the pyridyl or pyrazole rings completely obliterated activity suggesting that although these analogs engaged SHIP1, the arrangement of these hydrogen bond accepting nitrogen atoms is critical for activity in this assay.

Previously reported findings from our group found that reduced expression of *INPP5D*, which encodes SHIP1, in the 5xFAD murine amyloidosis model of AD promoted multiple protective responses of microglia that mitigate Aβ-mediated pathology including uptake of Aβ fibrils by microglia^60^. Therefore, for a subset of compounds, we sought to assess their ability to induce the uptake of aggregated Aβ fibrils by primary murine microglia. Wildtype microglia were isolated as described above. Aβ fibrils were prepared by incubating Alexa Fluor labeled Aβ peptide at 37 °C for five days (**Figure 6A**). Microglia were pretreated with test compound or vehicle control (DMSO) followed by the addition of Aβ fibrils. An initial time course study found that amyloid uptake plateaued after 6 hours of pretreatment with 1 µM of compound **32** (Supplementary **Figure S5**). Therefore, after 6 hours, the pre-formed Aβ fibrils were added to the culture medium, and cells were incubated for an additional 30 minutes. Cells were then washed to remove amyloid, and uptake was quantified using immunofluorescence. All experiments were performed in triplicate and compared to untreated cells exposed to Aβ. Microglial uptake of Aβ fibrils was assessed following treatment with compound **32**, the best performing analog in the pHrodo-myelin/membrane debris uptake assay, at 0.3 µM, 1 µM, and 3 µM. The activity of compound **32** was compared to the methyl **41** and acetamide **43** analogs. Gratifyingly, the results were consistent with activity in the BiTSA and pAKT/tAKT assay, as well as the PIP profiling activities described above (**Figure 6B**). Treatment with compound **32** significantly increased amyloid uptake compared to the control (p < 0.01) with a maximum 2-fold increase at 1 µM. The less active **41** showed a non-significant trend toward increased uptake at 0.3 µM and 1 µM, but uptake was nearly absent at 3 µM. Compound **43**, which completely failed to engage SHIP1, actually reduced amyloid uptake. In a separate experiment, the methyl-substituted enantiomers **37** and **38** were tested side-by-side, and their activities were consistent with their respective potencies in the pAKT/tAKT assay; the more potent enantiomer **37** was active in contrast to compound **38**. These results suggest that modulation of SHIP1 increases the uptake of Aβ fibrils by microglia, and that this activity is consistent with target engagement. Furthermore, compound **32** was not cytotoxic to primary microglia at concentrations up to 10 µM, and at 1–3 µM rescued these cells from the cytotoxic effects of fibrillar Aβ, as assessed by LDH release (**Figure 6C**).

**Figure 6.**
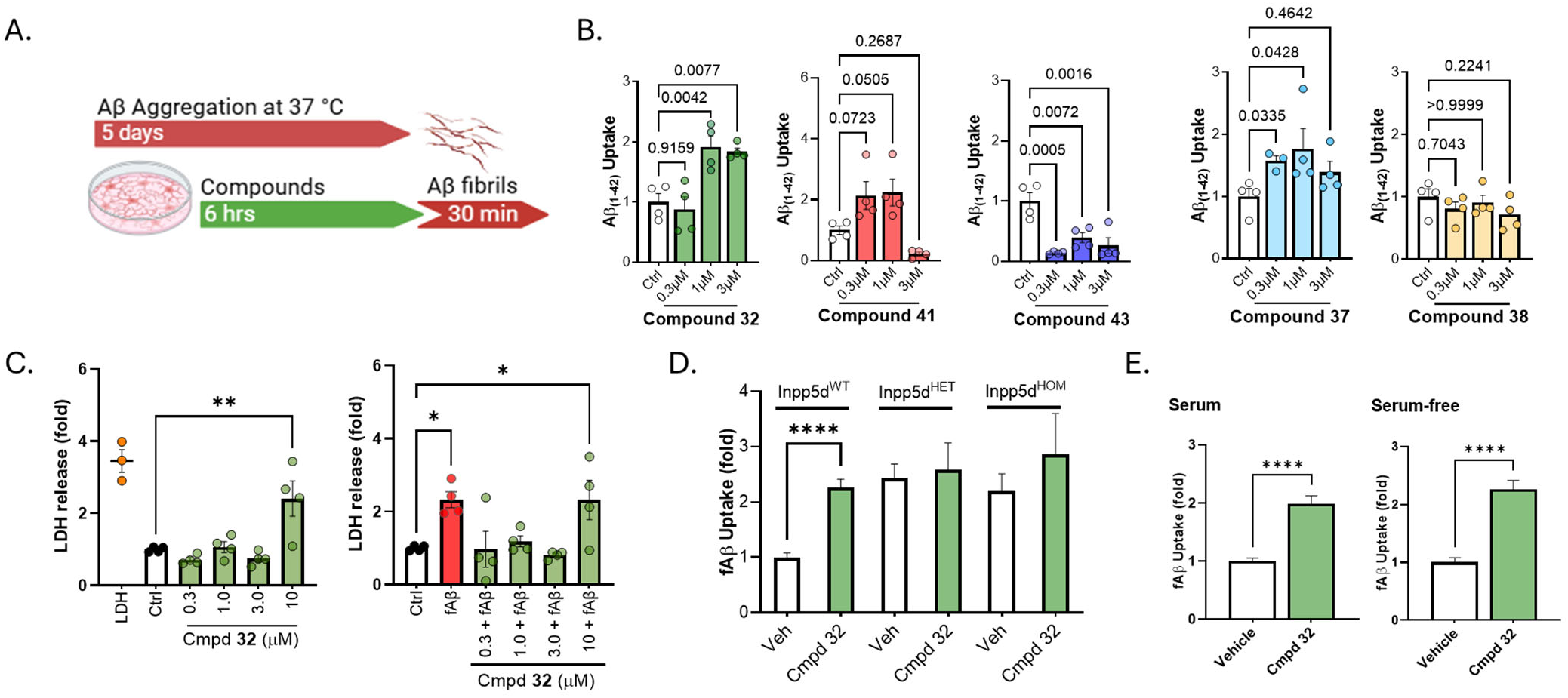
SHIP1-dependent microglial uptake of fibrillar Aβ1-42. **A.** Schematic of the experimental design for Aβ uptake assay. Primary microglial cultures were pretreated with SHIP1 compounds concentrations of 0.3, 1, and 3 µM for 6 hours, followed by a 30-minute incubation with fluorescently labeled fibrillar Aβ1-42. **B.** Bar graphs showing fold change in Aβ fibril uptake relative to vehicle control for each compound **32**, **41** and **43**; and in a separate experiment the enantiomers **37** and **38**. **C.** Compound **32** exhibited no cytotoxicity at concentrations up to 10 µM and mitigated fibrillar Aβ-induced cytotoxicity to primary microglia as assessed by LDH release. **D.** Treatment with compound **32** at 1 µM significantly enhanced Aβ uptake (∼2-fold) in *Inpp5d*^WT^ microglia compared to vehicle control (p < 0.0001). This effect phenocopied the increased uptake observed in *Inpp5d*^HET^ microglia. Notably, compound **32** did not alter Aβ fibril uptake in *Inpp5d*^HOM^ microglia. Data are presented as mean ± SEM; n = 4 biological replicates per group (eight scans analyzed per biological replicate). Statistical analysis was performed using one-way ANOVA followed by Tukey’s multiple comparisons test, or unpaired t test as appropriate (**** p < 0.0001). **E.** Assessment of Aβ fibril uptake by *Inpp5d*^WT^ microglia treated with 1 µM compound **32** in the presence or absence of serum. Data are presented as mean ± SEM; n = 4 biological replicates per group. Statistical analysis was performed using one-way ANOVA followed by Tukey’s multiple comparisons test, or unpaired t test as appropriate (**** p < 0.0001).

To provide evidence that the increased Aβ uptake observed with compound **32** is dependent on SHIP1, we repeated the study described above using primary microglia isolated from *Inpp5d^HET^* and *Inpp5d^HOM^* mice, which have reduced and loss of SHIP1, respectively, compared to *Inpp5d^WT^* mice. As shown in **Figure 6D**, treatment with compound **32** significantly increased Aβ uptake 2-fold by *Inpp5d^WT^* microglia at 1 µM compared to vehicle control (p < 0.0001). Importantly, compound **32** phenocopied the level of Aβ uptake by *Inpp5d^HET^* microglia. These results demonstrate that pharmacological modulation of SHIP1 by compound **32** achieves similar effects compared to decreased SHIP1 expression, which we have previously reported promotes multiple protective responses of microglia. Furthermore, compound **32** did not alter the uptake of Aβ fibrils by *Inpp5d^HOM^* knockout microglia, supporting that its activity is dependent on SHIP1, and providing additional evidence that this desired pharmacology is mediated by SHIP1. Since compound **32** is highly protein-bound (*see below*), we performed a serum shift assay (**Figure 6E**). We compared amyloid uptake by *Inpp5d*^WT^ microglia treated with 1 µM of compound **32** in the presence or absence of serum. As shown, this compound significantly increased Aβ fibril uptake equally under both serum and serum-free conditions (p < 0.0001). Because uptake remained similar and robust in both conditions, these results suggest that despite its high protein binding, compound **32** is likely to retain activity *in vivo*.

### Kinome profiling

Because compound **32** has structural features similar to crizotinib, a multi-targeted kinase inhibitor^64^, we sought to discharge the risk that the pharmacology of compound **32** is due to kinase inhibition. Results from broad biochemical kinase panels can be difficult to interpret with respect to activities in specific cellular contexts because they lack the biological complexity of intact cells, especially the relevant ATP concentrations and kinase activation states, which can lead to misleading conclusions that do not translate to observed pharmacology^65^. Therefore, we opted to explore this possibility using a targeted approach focused on observed cellular activities and a broad approach within the context of desired cellular pharmacology. Since we observed an increase in monophosphorylated PIPs such as PI(4)P when cells were treated with compound **32**, we tested the inhibitor in a panel of phosphatidylinositol-4-phosphate 5-kinases (PIP5K1A, PIP5K1B, and PIP5K1C), which are responsible for phosphorylating PI(4)P at the 5-position of the inositol ring, producing PI(4,5)P_2_, an outcome potentially similar to inhibiting SHIP1, which also produces PI(4,5)P_2_. Compound **32** was inactive at PIP5K1A and PIP5K1C up to 10 µM and resulted in only 50% inhibition of PIP5K1B at 10 µM. See supplementary **Table S1**. Notably, the SHIP1 inactive compounds **41** and **43** had similar levels of activity against these kinases suggesting that the observed SAR in the cellular assays described above is not related to PIP5K1B inhibition. We wanted to further ensure that the cellular activity of compound **32** is not due to kinase inhibition in general within the context of BV2 cells where we observed increased uptake of myelin/membrane debris. Therefore, we executed a kinome profiling study of compound **32** compared to crizotinib as a positive control using BV2 cell lysates. With crizotinib treatment both Aurkb and Mertk had decreased kinase binding consistent with a previous report using similar methodology^66^. See supplementary **Figure S6**. Importantly, compound **32** did not cause significant changes in binding to any kinases across the concentration range up to 3 µM that resulted in enhanced uptake of pHrodo-labeled myelin/membrane debris or fibrillar Aβ. These data provide evidence that the observed cellular pharmacology of compound **32** is not due to kinase inhibition.

### Pharmacokinetic (PK) studies

To assess the suitability of compound **32** for animal studies, we first determined physicochemical and ADME properties in the following assays: kinetic solubility in pH 7.4 phosphate buffer, microsomal stability/intrinsic clearance (CL_int_) in mouse liver microsomes, MDCK (Madin-Darby Canine Kidney) permeability (Papp), and protein binding (fu) in mouse plasma and brain (**Table 8**). Compound **32** demonstrated favorable properties with high solubility exceeding 100 µM, which combined with the high permeability observed in the MDCK assay (Papp of 49.4 x10^-6 cm/s), indicates good potential for absorption. The solubility of compound **32** was >5 mg/mL in hydroxypropylmethylcellulose (HPMC) (1%) with Tween 80 (0.25%) in purified water, which was the formulation used for *in vivo* studies. The metabolic stability of compound **32** was good with moderate intrinsic clearance observed (CL_int_ of 84.5 µL/min/mg). Protein binding studies revealed low plasma unbound fraction (f_u_) at 0.055, while brain unbound fraction was even lower at 0.0037, suggesting limited free drug availability, however the serum shift assay described above suggested this high protein binding may not limit desired pharmacology *in vivo*. Therefore, we conducted a single-dose, time course PK study in adult male C57BL/6J mice. Compound **32** (100 mg/kg) was formulated (5 mg/ml) and administered to non-fasted mice via oral gavage (20 ml/kg). Plasma exposures were obtained from two cohorts of mice following serial bleeds and a terminal blood collection. Cohort 1 had blood collected at 0.5, 2, 6, 8, and 24 hours post dose; while cohort 2 had blood collected at 1, 4, 8, 12, and 24 hours post dose. Brain exposures were obtained at terminal 24 hours post dose. Average exposures over time are depicted in **Figure 7** and the corresponding PK parameters are reported in **Table 9**. Compound **32** achieved a maximum concentration (C_max_) of 6 µM in plasma and maintained a mean brain concentration above 0.6 µM at 24 hours, achieving concentrations consistent with desired pharmacology in the cellular assays described above thereby supporting advancement to pharmacodynamic (PD) studies.

**Figure 7.**
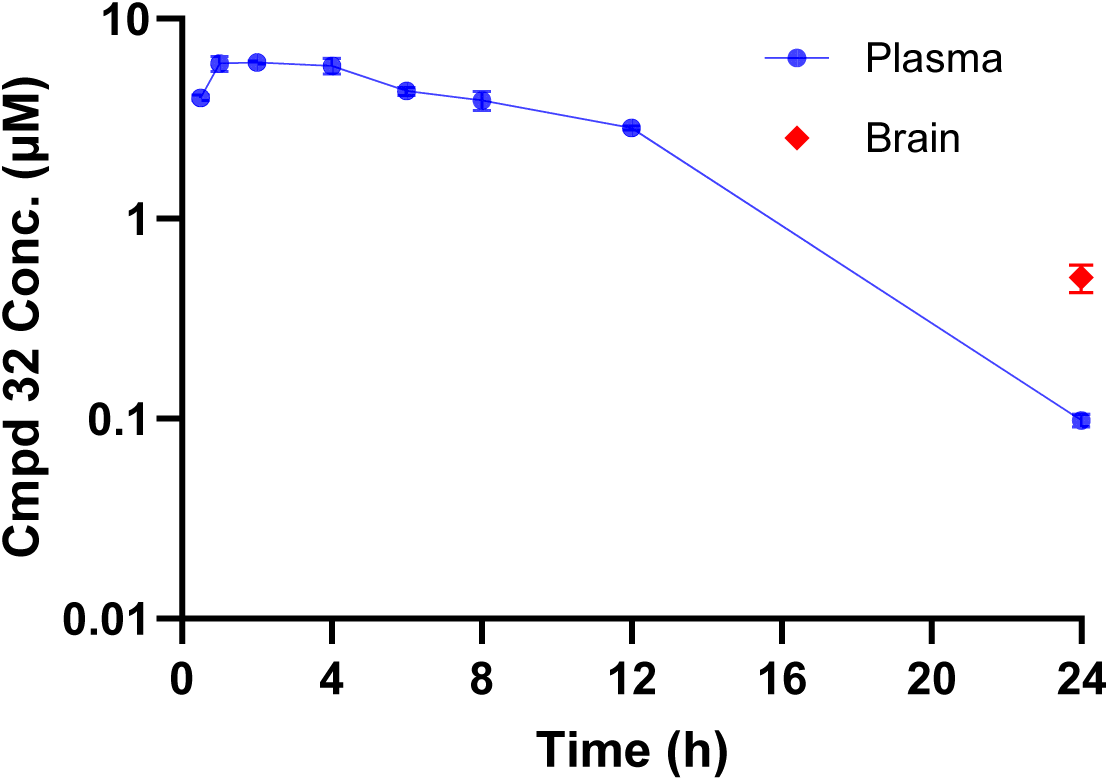
Pharmacokinetics (PK). Plot of time vs. plasma concentration for the pharmacokinetics of compound **32** illustrating the data shown in **Table 9**. Plasma (blue) and brain (red) concentrations are shown following an oral administration of 100 mg/kg to B6 mice. Compound **32** demonstrated significant exposures in both plasma and brain with a brain/plasma ratio of 0.5–3-fold that increased over time.

**Table 8.**
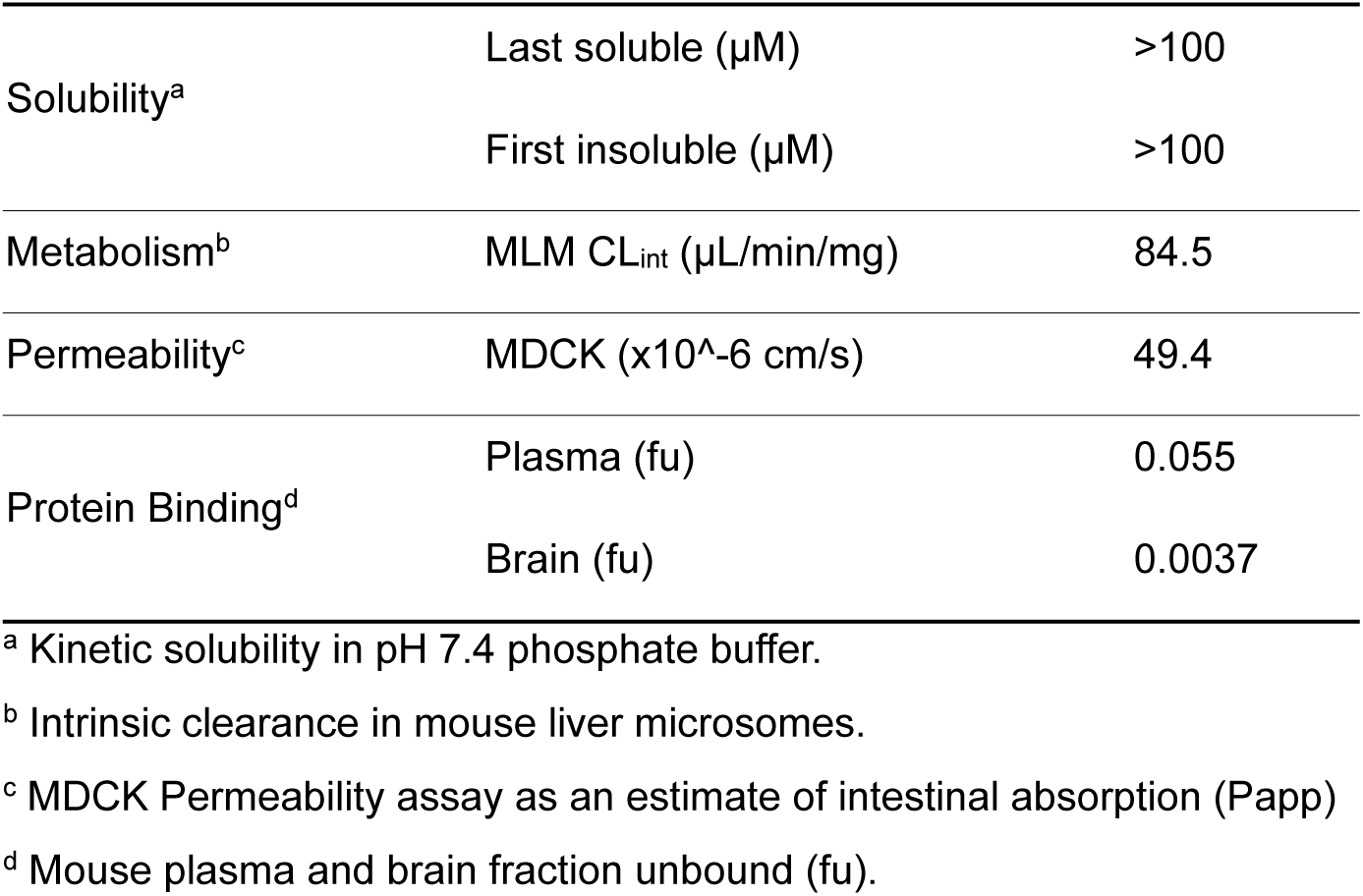
Physicochemical and ADME properties of compound 32.

**Table 9.**
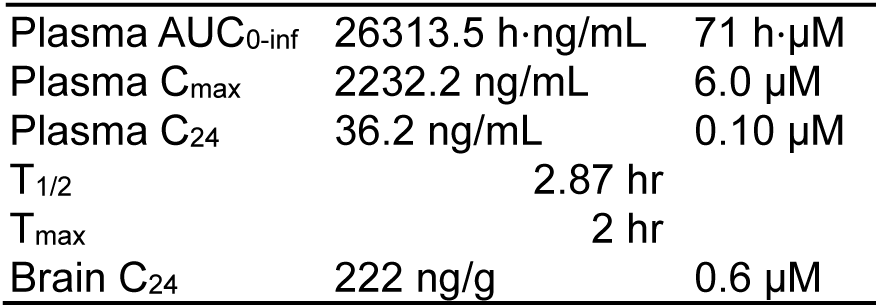
Pharmacokinetic parameters of compound 32 after a single *p.o.* dose (100 mg/kg) in mice.

### Pharmacodynamic (PD) Studies

The hAβ^SAA^ knock-in (SAA) mouse is a genetically engineered animal model in which the endogenous *App* locus encodes a humanized Aβ sequence containing three familial AD-linked mutations, the Swedish (KM670/671NL), Arctic (E693G), and Austrian (T714I), which accelerate amyloid deposition^67^. SAA mice exhibit age-dependent accumulation of Aβ plaques, detectable at 4-months of age, as well as cerebral amyloid angiopathy, dystrophic neurites, and microgliosis around plaques. We selected this model to evaluate the PD effects of SHIP1 modulation by compound **32** because it is openly available, avoids the overexpression artifacts and sexual dimorphism seen in other amyloidosis models such as the 5xFAD, and most importantly the transcriptomic profile of plaque-associated microglia in SAA mice resemble that of microglia from human AD brains. We conducted a study in 14-month-old SAA mice and compared results to age-matched C57BL/6J mice, the wild-type background strain for the SAA line. Mice were treated with compound **32**, formulated as described above, and dosed at 150 mg/kg at 0, 12, and 24 hours (**Figure 8A**). At the 1-hr post-dose timepoint plasma was collected and at the terminal 25-hr endpoint, both blood and brain tissues were collected. Compound **32** achieved similar exposure levels in both sexes and at both timepoints in plasma and exceeded levels in brain required for activity in the cellular assays described above (**Figure 8B**). For transcriptomic analyses, hemibrains were homogenized, RNA was isolated, and each sample was analyzed using murine neuroinflammation and glial NanoString nCounter panels. Differential gene expression analysis revealed sex-specific differences in neuroinflammatory and glial transcriptomic responses, which were normalized following treatment of SAA mice with compound **32** (**Figure 8C**). The sex differences observed suggest more neuroinflammation in female SAA mice, which had elevated gene expression compared to males. As a result, while gene expression levels were reduced in both sexes, differences with treatment were more apparent in male mice, which had increased levels of gene expression in both panels that were normalized to B6 wild type levels. Normalization in gene expression was also observed in females treated with compound **32** but they did not quite reach B6 levels.

**Figure 8.**
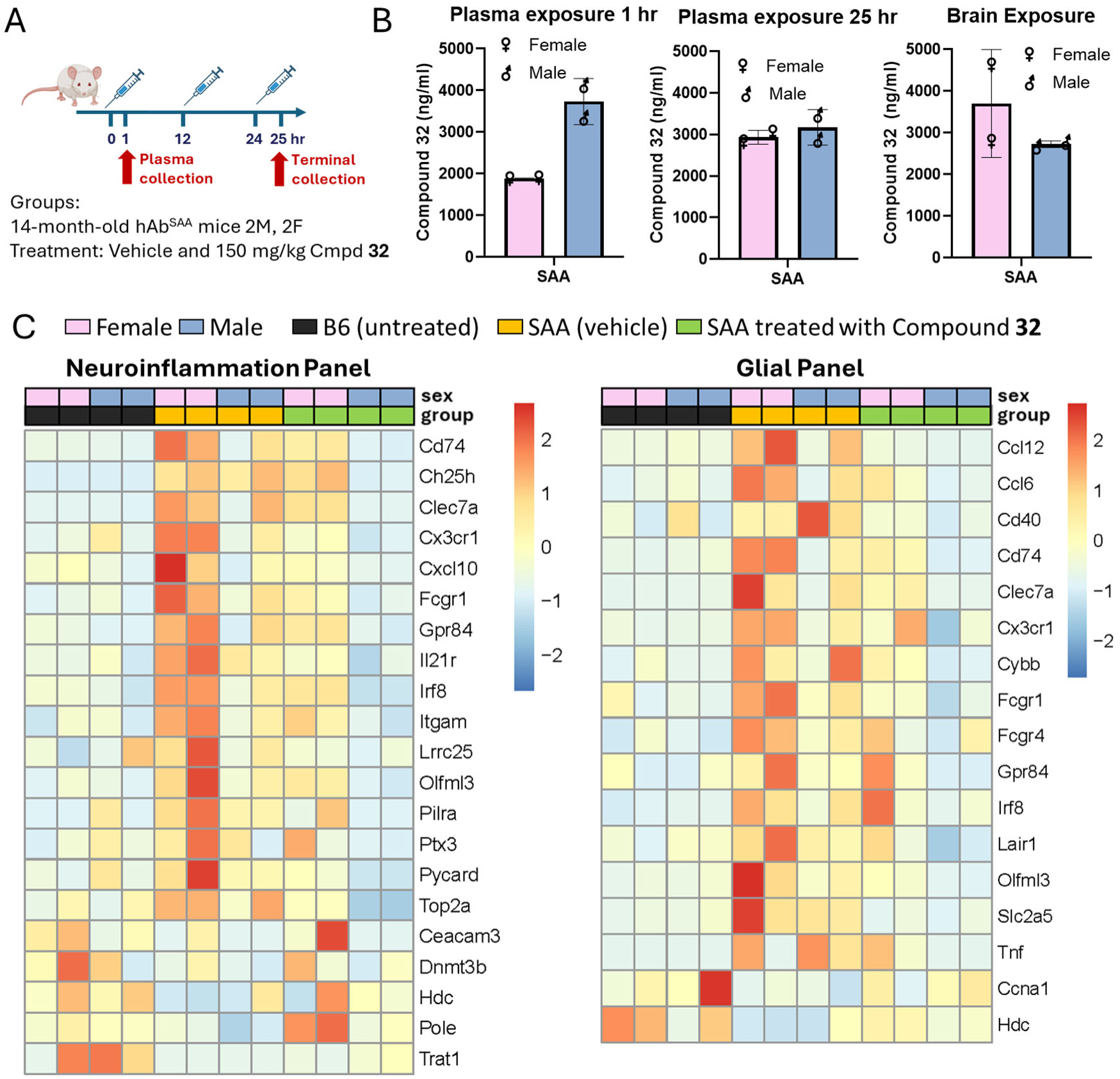
Acute PD study in wildtype (B6) and the human APP-SAA knock-in mouse model of AD (SAA). **A.** 14-month-old hAβ^SAA^ knock-in (SAA) mice (n = 2 males, 2 females per group) were treated with either vehicle or compound **32** formulated in 1% HPMC / 0.25% Tween 80 / purified water and administered at 150 mg/kg (10 mL/kg body weight) via oral gavage at 0, 12, and 24 hours. Plasma was collected 1 hr post dose and both plasma and brain were collected at the 25 hr terminal end of the study. **B.** Compound **32** concentrations quantified in plasma and brain homogenates using HPLC-MS/MS. **C.** For transcriptomic analysis, hemibrains were homogenized, RNA was isolated, and 200 ng of total RNA from each sample was profiled using the NanoString nCounter Mouse Neuroinflammation and Glial panels. The heatmaps display z-score normalized expression for differentially expressed genes across genotype and treatment conditions comparing untreated wildtype B6 mice to SAA mice treated with vehicle to SAA mice treated with compound **32**.

Cytokine protein levels were quantified from hemibrain homogenates using the Meso Scale Discovery (MSD) V-PLEX Mouse Proinflammatory Panel I (Meso Scale Diagnostics, K15048D). This multiplex immunoassay enables sensitive detection of multiple cytokines in parallel standardized to total protein content (pg/mg protein). Supplementary **Figure S7** shows results for IFN-γ, IL-1β, IL-2, IL-4, IL-5, IL-6, KC/GRO, TNF-α, and IL-12p70. Most of these cytokines were not significantly different between wild-type B6 and SAA mice, except for IL-1β, which was elevated in SAA mice. As shown in **Figure 9**, treatment with compound **32** reduced IL-1β expression in SAA mice to levels comparable to wild-type B6 controls. To confirm that amyloid drives this increased IL-1β expression, primary microglia from wild-type mice were treated with fAβ, which increased IL-1β protein expression, while co-treatment with compound **32** reversed the effect, consistent with the *in vivo* findings. Together, these results are consistent with a central pharmacodynamic response to SHIP1 modulation that normalizes amyloid-induced neuroinflammatory IL-1β signaling.

**Figure 9.**
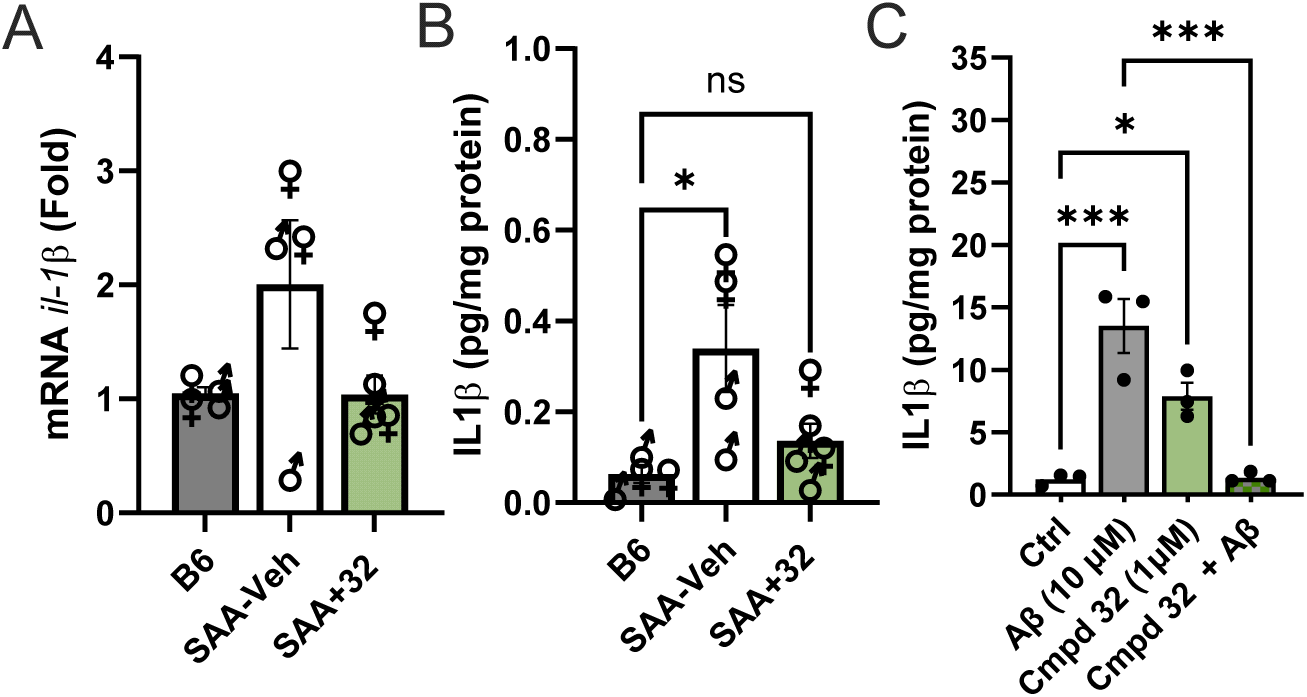
SHIP1 target engagement reduces brain and microglial expression of the pro-inflammatory cytokine IL-1β. **A.** *Il1β* mRNA expression measured by qPCR in the brains of SAA mice showed an increasing trend compared to B6 controls, while oral administration of compound **32** appeared to reduce expression. **B.** IL-1β protein levels were significantly elevated in the brains of SAA mice relative to B6 controls as measured by the MSD Cytokine Assay and are reported as pg protein per mg total protein. Oral treatment of SAA mice with compound **32** reduced IL-1β protein expression. **C.** Primary microglia treated with fAβ exhibited increased IL-1β protein expression, which was reversed by compound **32** treatment. Data are presented as mean ± SEM; n = 4–5 per group for *in vivo* studies and n=3 for primary microglial cultures. Statistical analysis was performed using the Kruskal–Wallis test followed by Dunn’s multiple comparisons test (* p< 0.05, *** p< 0.001).

## CONCLUSIONS

Phosphatases have long been considered challenging “undruggable” targets^68^. More recently, the protein phosphatase inhibitor SHP099 has been reported as a potent and selective allosteric inhibitor that binds an interface between the N- and C-terminal SH2 domains of SHP2, locking the enzyme in an inactive conformation^69^. This compound and a related analog TNO155 have progressed into clinical studies^70^, suggesting that targeting the allosteric sites of protein phosphatases can be a successful strategy. Therefore, we began our biochemical studies with a protein construct containing both the phosphatase (Ptase) and C2 domains, which includes an allosteric site previously studied as a potential binding site for phosphatidylserine and PI(3,4)P_2_. These lipids have been reported to inhibit and activate respectively the Ptase domain through successive feed-back and forward loops as PI(3,4,5)P_3_ is converted to PI(3,4)P_2_^23, 71, 72^. Fragment-based screening of the SHIP1 Ptase–C2 protein identified multiple ligands bound at this same inter-domain cleft^46^, suggesting small molecules can bind this site. Although these previous studies support the viability of targeting this allosteric site between the Ptase and C2 domains of SHIP1, our efforts to use a traditional, cell-free biochemical assay did not translate meaningfully to cellular activity. Such discrepancies have been observed with other complex enzymes^73, 74^. We believe these negative results support the need to avoid an over simplified view of SHIP1 functioning only as an enzyme rather than a complex, multidomain, multifunctional protein that mediates multiple protein-protein interactions while serving as an interfacial enzyme at membrane surfaces. Therefore, we enabled a Bioluminescence-based Thermal Shift Assay (BiTSA) to study SHIP1 target engagement in intact cells^48, 75, 76^ in conjunction with a signaling assay (pAKT/tAKT) to provide additional evidence that observed cellular pharmacological activities are on-target^77, 78^.

With this strategy in place, we surveyed a broad range of reported inhibitors and focused on one series, the pyridyl-pyrazole-piperidine scaffold represented by compound **9**, because it was unique in its ability to affect desired pharmacology at concentrations without confounding cytotoxicity. Our SAR studies identified compound **32**, which consistently increased the uptake of myelin/membrane debris at concentrations that were not cytotoxic to either BV2 cells or primary murine microglia. Given this desired pharmacology we sought to confirm SHIP1 target engagement by compound **32** with several lines of supporting evidence. Importantly, the rank order SAR observed in BiTSA tracked with reductions in pAKT/tAKT, a relationship consistent with target engagement and expected changes in the plasma membrane PIP population. Indeed, mass-spectral analysis of PIP species in THP-1 cells treated with compound **32** compared to its methyl **41** and acetamide **43** analogs revealed a shift that aligned with corresponding activities in the BiTSA and pAKT/tAKT assay. Furthermore, compound **32** shifted the protein’s binding preferences from monophosphorylated PIPs toward di- and tri-phosphorylated forms, with a marked increase in its association with the product of its phosphatase activity, PI(3,4)P_2_, as measured by the protein lipid overlay (PLO) assay. Finally, compound **32** increased the uptake of fibrillar amyloid, consistent with our prior findings that primary murine microglia from *Inpp5d*^HET^ mice with reduced SHIP1 expression exhibit enhanced phagocytic clearance of amyloid^5^. This phenotype is consistent with the transcriptional and functional signatures of disease-associated microglia that have been reported to confer protection in AD. Moreover, enhanced amyloid uptake represents a mechanistic overlap with anti-amyloid immunotherapies, which facilitate microglia-mediated Aβ clearance and have demonstrated efficacy in slowing cognitive decline in clinical studies. Importantly, compound **32** did not alter the behavior of *Inpp5d^HOM^* knockout microglia, supporting that its activity is dependent on SHIP1, and providing additional evidence that the desired pharmacology compound **32** is mediated by SHIP1.

The unique and desired activity of compound **32** may be due in part to off-target activities. Given the similarity in structure of compound **32** to crizotinib we sought to evaluate its activity against kinases. Rather than relying on a broad kinase screening panel, which can be difficult to interpret with regard to a particular cellular context, we evaluated the most likely kinase targets, the PIP5K family, given the observed shifts in PIP species. We also performed a cellular kinome profiling study using the same BV2 cells in which desired pharmacology is observed. The SAR of PIP5K family inhibition did not align with cellular activity and the in cell kinome profiling failed to identify any kinases that were bound by compound **32**. Although not an exhaustive search for off-targets, which will be the subject of future studies, these results suggest that the activity of compound **32** is not kinase dependent.

We have demonstrated sufficient brain exposure and a central pharmacodynamic effect of SHIP1 modulation in mice. An important consideration in interpreting the activity of compound **32** in mouse studies is its high protein binding, which influences the free fraction available for activity *in vivo*. Fortunately, compound **32** retained its ability to enhance amyloid uptake by primary microglia in a serum shift assay, indicating that its activity is not significantly diminished by protein binding. Furthermore, compound **32** altered microglial transcriptional profiles and IL-1β levels in SAA mouse brain after only 25 hours of exposure. One explanation for these activities despite high protein binding is the amphiphilic nature of compound **32**. Amphiphilic compounds with affinity for both hydrophilic and hydrophobic environments, commonly exhibit high protein binding but they can also partition into cellular membranes. The observation that the effect of compound **32** on the PI pool in THP-1 cells was dependent on the acyl chains of the detected species suggests that membrane partitioning may be important for its activity against SHIP1. Features of the SAR around its lipophilic benzyl tail and the basic amine of the piperidine may reflect modulating membrane partitioning and protein binding, providing an alternative to the traditional "free drug" hypothesis, which assumes only the unbound aqueous fraction is pharmacologically active^79^. These results are similar to the SAR observed for ligands targeting membrane-integral proteins like GPCRs, especially those with hydrophobic endogenous ligands, which interact within the bilayer where localized concentrations, binding kinetics, and affinity can be strongly influenced by the membrane microenvironment^80^. SHIP1 is a membrane-associated protein with several domains that may interact with lipids at the membrane surface such as phosphatidylserine, PI(4,5)P₃, and PI(3,4,5)P₃. Unlike soluble enzyme inhibitors, an amphiphilic compound like **32** that has the potential to partition into lipid bilayers and accumulate near its target may be ideal for effective SHIP1 modulation.

Although we have provided evidence for cellular target engagement and dependence on SHIP1 for the desired pharmacology, the precise binding location and mode of action of compound **32** remain unknown. Additional structural and biophysical studies will be needed to define where and how it interacts with SHIP1, and whether its activity reflects direct occupancy of a putative allosteric site, inhibition of protein interactions with SHIP1, modulation of its lipid interactions at the membrane surface, or some combination of these potential modes of action. While we sought to address the risk that its activity might be due to kinase inhibition, an exhaustive search for off-targets is still needed. Combined with an understanding of its mechanism and proteome-wide activities, further SAR studies will focus on improving potency and reducing potential off-target liabilities. More broadly, our findings highlight the feasibility of targeting SHIP1 in the CNS and support a strategy in which an amphiphilic scaffold can achieve sufficient target engagement of a membrane-associated inositol phosphatases by exploiting lipid microenvironments and measuring activity using cellular protein thermostability and signaling assays, rather than relying on free drug concentrations and cell-free enzyme assays. By demonstrating that SHIP1 modulation enhances microglial uptake of amyloid in a manner similar to a genetic model, this work provides a foundation for therapeutic development aimed at harnessing microglia to reduce amyloid levels and slow the progression of cognitive decline in AD in a complementary manner related to the activity of the amyloid targeting antibodies recently approved for clinical use. Finally, future efforts to evaluate compound **32** in long-term efficacy and safety studies will be essential next steps toward translating SHIP1 inhibition into a viable therapeutic strategy.

## METHODS

### Chemistry

Reagents and solvents were purchased from commercial sources and used without further purification. All reactions involving air- or moisture-sensitive reagents were performed under a nitrogen or argon atmosphere. NMR spectra were recorded on Bruker Avance Neo 400 MHz or DRX500-1 500 MHz instruments. For ^1^H NMR, chemical shifts in ppm relative to the residual solvent peak, multiplicities, coupling constants in Hertz, and numbers of protons are indicated. Reactions were routinely monitored by analytical TLC and/or LC-MS. Analytical TLC was performed on silica gel 60 F_254_ silica gel plates, and 254 nm UV light and/or I_2_ staining were used for visualization. Reaction monitoring LC-MS data were obtained on a Waters Acquity UPLC system equipped with UV (TUV or PDA), mass (SQD2 or QDa), and/or evaporative light scattering detectors. Flash NP or RP chromatography were performed on a Teledyne-ISCO NextGen 300 instrument using prepacked silica gel or C18-functionalized silica gel columns available from Teledyne-ISCO, or on a Teledyne-ISCO Combiflash using prepacked silica gel columns available from Agela or Welch. High-resolution mass spectra were obtained on an Agilent 6550 Q-TOF instrument.

All compounds are >95% purity as determined by HPLC. One of the following specified methods was used to determine test compound purity: Methods **A:** mobile phase A, 0.1% TFA in H_2_O; mobile phase B, ACN; column, Acquity HSS-T3 (1.8 μm, 100 mm x 2.1 mm); flow rate, 0.3 mL/min; detection wavelength, 214 nm; column temperature, 30°C. **B:** mobile phase A, 5 mM NH_4_OAc in H_2_O; mobile phase B, ACN; column, Acquity BEH C18 (1.7 μm, 100 mm x 2.1 mm); flow rate, 0.3 mL/min; detection wavelength, 214 nm; column temperature, 30 °C. **C:** mobile phase A, 0.1% TFA in H_2_O; mobile phase B, MeOH; column, ThermoScientific Hypersil GOLDC18 column (1.8 μm, 100 mm x 2.1 mm); flow rate, 0.5 mL/min; detection wavelength, 254 nm; column temperature, 30 °C.

High-resolution mass spectrometry data is provided for compounds with associated *in vitro* data. NMR and HPLC traces are included in the supplementary information for key compounds with *in vitro* data and for compound **32**.

### *tert-*Butyl 4-(4-(5-hydroxypyridin-3-yl)-1*H*-pyrazol-1-yl)piperidine-1-carboxylate (16)

Potassium carbonate (1.59 g, 2 eq., 11.5 mmol) was added to a solution of 5-bromopyridin-3-ol (**14,** 1 g, 5.75 mmol) and tert-butyl 4-(4-(4,4,5,5-tetramethyl-1,3,2-dioxaborolan-2-yl)-1*H*-pyrazol-1-yl)piperidine-1-carboxylate (**15,** 2.17 g, 5.75 mmol) in toluene (5 mL), ethanol (5 mL) and water (2 mL,), and reaction mixture was purged with nitrogen for 20 minutes. [1,1′-bis(diphenylphosphino)ferrocene]dichloropalladium(II) (234 mg, 0.05 eq., 287 µmol) was then added and reaction mixture was heated at 100°C and stirred for 16h. The reaction mixture was cooled, diluted with water and extracted with EtOAc. The EtOAc extracts were washed with brine, dried over anhydrous sodium sulfate, filtered and concentrated. The crude material was purified by silica gel flash chromatography using 40% EtOAc in hexane as an eluent to afford *tert-*butyl 4-(4-(5-hydroxypyridin-3-yl)-1*H*-pyrazol-1-yl)piperidine-1-carboxylate (0.5 g, 1.39 mmol, 24.2%) as a white solid. ^1^H NMR (400 MHz, DMSO-*d*_6_) δ 9.86 (s, 1H), 8.33 (s, 1H), 8.30 (d, 1H, *J =* 1.6 Hz), 7.96 (d, 1H, *J* = 2.4 Hz), 7.91 (s, 1H), 7.30 (t, 1H, *J =* 2.2 Hz), 4.40-4.33 (m, 1H), 4.05 (d, 2H, *J =* 12 Hz), 2.92 (br s, 2H), 2.03 (d, 2H, *J =* 10.4 Hz), 1.95 (qd, 2H, *J =* 4.0, 12.0 Hz), 1.41 (s, 9H); LRMS (ES+) m/z 345.33 [M+H]^+^; LC-MS purity 98.1% (Method A).

### *tert*-Butyl 4-(4-(5-((2-chlorobenzyl)oxy)pyridin-3-yl)-1*H*-pyrazol-1-yl)piperidine-1-carboxylate (24) (TAD-000029)

Potassium carbonate 321 mg, 2 eq., 2.32 mmol) was added portion wise to a stirred solution of *tert*-butyl 4-(4-(5-hydroxypyridin-3-yl)-1*H*-pyrazol-1-yl)piperidine-1-carboxylate (**16,** 0.4 g, 1.16 mmol) and 1-(bromomethyl)-2-chlorobenzene (263 mg, 1.1 eq., 1.28 mmol) in dimethylformamide (7 mL), and reaction mixture heated at 100°C for 16h. The reaction mixture was diluted with water and extracted with EtOAc. The EtOAc extracts were washed with brine, dried over anhydrous sodium sulfate, filtered and concentrated. The crude material was purified by silica gel flash chromatography using 40% EtOAc in hexane as an eluent to afford *tert*-butyl 4-(4-(5-((2-chlorophenyl)methoxy]pyridin-3-yl)-1*H*-pyrazol-1-yl)piperidine-1-carboxylate (180 mg, 365 µmol, 31.4%) as an off white solid. ^1^H NMR (400 MHz, DMSO-*d*_6_) δ 8.49 (s, 1H), 8.44 (s, 1H), 8.18 (d, 1H, *J =* 2.4 Hz), 8.03 (s, 1H), 7.72 (m, 1H), 7.67-7.65 (m, 1H) 7.55-7.53 (m, 1H), 7.45-7.41 (m, 2H), 5.27 (s, 2H), 4.41-4.35 (m, 1H), 4.05 (d, *J =* 11.6 Hz, 2H), 2.94 (br s, 2H), 2.03 (d, 2H, *J =* 12.8 Hz), 1.79 (qd, 2H, *J =* 4.0, 12.0 Hz), 1.42 (s, 9H); LRMS (ES+) m/z 469.22 [M+H]^+^; LC-MS purity 97.3% (Method A); Calculated [M+H]^+^ for C_25_H_29_ClN_4_O_3_: 469.2006; Found: 469.2007.

The following compounds were made by alkylation of *tert*-butyl 4-(4-(5-hydroxypyridin-3-yl)-1*H*-pyrazol-1-yl)piperidine-1-carboxylate (**16**) with the corresponding benzyl bromide:

### tert-Butyl 4-(4-(5-((2,4-dichlorobenzyl)oxy)pyridin-3-yl)-1*H*-pyrazol-1-yl)piperidine-1-carboxylate (intermediate)

1H NMR (400 MHz, DMSO-*d*6): δ 8.49 (s, 1H), 8.44 (s, 1H), 8.18 (d, 1H, *J =* 2.4 Hz), 8.04 (s, 1H), 7.72-7.50 (m, 4H), 5.36 (s, 2H), 4.39-4.35 (m, 1H), 4.05 (d, 2H, *J =* 11.1 Hz), 2.95-2.93 (m, 2H), 2.50-2.49 (m, 2H), 1.83-1.77 (m, 2H), 1.42 (s, 9H). LRMS (ES+) m/z 503.25 [M+H]^+^; LC-MS purity 97.2% (Method A).

### *tert*-Butyl 4-(4-(5-((2,6-dichlorobenzyl)oxy)pyridin-3-yl)-1*H*-pyrazol-1-yl)piperidine-1-carboxylate (intermediate)

^1^H NMR (400 MHz, DMSO-*d*_6_) δ 8.52 (d, 1H, *J* = 1.6 Hz), 8.45 (s, 1H), 8.20 (d, 1H, *J =* 2.8 Hz), 8.05 (s, 1H), 7.70 (t, 1H, *J* = 2.0 Hz,), 7.60-7.48 (m, 3H), 5.36 (s, 2H), 4.42-4.34 (m, 1H), 4.05 (d, 2H, *J =* 11.6 Hz), 2.94 (br s, 2H), 2.06 (d, 2H, *J =* 10.8 Hz), 1.79 (qd, 2H, *J =* 4.1, 12.1 Hz), 1.42 (s, 9H); LRMS (ES+) m/z 503.30 [M+H]^+^; LC-MS purity 99.0% (Method A).

### *tert*-Butyl 4-(4-(5-((4-chlorobenzyl)oxy)pyridin-3-yl)-1*H*-pyrazol-1-yl)piperidine-1-carboxylate (intermediate)

^1^H NMR (400 MHz, DMSO-*d*_6_) δ 8.47 (d, 1H, *J* = 1.7 Hz), 8.42 (s, 1H), 8.16 (d, 1H, *J =* 2.8 Hz), 8.01 (s, 1H), 7.68-7.67 (m, 1H), 7.53-7.47 (m, 4H), 5.22 (s, 2H), 4.41-4.35 (m, 1H), 4.05 (d, 2H, *J =* 12.1 Hz), 2.93 (br s, 2H), 2.05 (m, 2H), 1.79 (qd, 2H, *J =* 4.3, 12.2 Hz1.42 (s, 9H); LRMS (ES+) m/z 469.22 [M+H]^+^; LC-MS purity 98.4% (Method A).

### *tert*-Butyl 4-(4-(5-((2-methylbenzyl)oxy)pyridin-3-yl)-1*H*-pyrazol-1-yl)piperidine-1-carboxylate (intermediate)

^1^H NMR (400 MHz, DMSO-*d*_6_) δ 8.47 (d, 1H, *J* = 1.6 Hz), 8.42 (s, 1H), 8.18 (d, 1H, *J =* 2.4 Hz), 8.02 (s, 1H), 7.70 (t, 1H, *J* = 2.2 Hz), 7.45 (d, 1H, *J* = 7.2 Hz), 7.29-7.20 (m, 3H), 5.20 (s, 2H), 4.42-4.34 (m, 1H), 4.05 (d, 2H, *J =* 11.6 Hz), 2.95 (br s, 2H), 2.35 (s, 3H), 2.05 (d, 2H, *J =* 10 Hz), 1.79 (qd, 2H, *J =* 4.4, 12.2 Hz), 1.42 (s, 9H); LRMS (ES+) m/z 449.38 [M+H]^+^; LC-MS purity 99.2% (Method A).

### *tert*-Butyl 4-(4-(5-((2-fluorobenzyl)oxy)pyridin-3-yl)-1*H*-pyrazol-1-yl)piperidine-1-carboxylate (intermediate)

^1^H NMR (400 MHz, DMSO-*d*_6_) δ 8.48 (d, 1H, *J* = 1.6 Hz), 8.44 (s, 1H), 8.18 (d, 1H, *J =* 2.4 Hz), 8.03 (s, 1H), 7.72 (t, 1H, *J* = 2.2 Hz), 7.63-7.59 (m, 1H) 7.48-7.42 (m, 1H), 7.30-7.24 (m, 2H), 5.26 (s, 2H), 4.42-4.34 (m, 1H), 4.05 (d, 2H, *J =* 12 Hz), 2.96 (br s, 2H), 2.05 (d, 2H, *J =* 10 Hz), 1.79 (qd, 2H, *J =* 4.2, 12.2 Hz), 1.42 (s, 9H); LRMS (ES+) m/z 453.38 [M+H]^+^; LC-MS purity 99.8% (Method A).

### *tert*-Butyl 4-(4-(5-(benzyloxy)pyridin-3-yl)-1*H*-pyrazol-1-yl)piperidine-1-carboxylate (intermediate)

^1^H NMR (400 MHz, DMSO-*d*_6_) δ 8.46 (d, *J* = 1.6 Hz, 1H), 8.42 (s, 1H), 8.16 (d, 1H, *J =* 2.8 Hz), 8.01 (s, 1H), 7.68 (t, 1H, *J* = 2.2 Hz), 7.50-7.33 (m, 5H), 5.21 (s, 2H), 4.41-4.35 (m, 1H), 4.05 (d, *J =* 10.8 Hz, 2H), 2.94 (br s, 2H), 2.05 (d, 2H, *J =* 10.4 Hz), 1.79 (qd, 2H, *J =* 4.2, 12.1 Hz), 1.42 (s, 9H); LRMS (ES+) m/z 435.40 [M+H]^+^; LC-MS purity 99.9% (Method A).

### 3-((2-Chlorobenzyl)oxy)-5-(1-(piperidin-4-yl)-1*H*-pyrazol-4-yl)pyridine. HCl (32) (TAD-0000032)

4M HCl in dioxane (2 mL) was added dropwise to a solution of tert-butyl 4-(4-(5-((2-chlorobenzyl)oxy)pyridin-3-yl)-1*H*-pyrazol-1-yl)piperidine-1-carboxylate (**24,** 180 mg, 384 µmol) in dichloromethane (5 mL, 78.1 µmol) at 0°C. The reaction mixture allowed to warm to at room temperature and stirred for 3h. The reaction mixture was concentrated and triturated with *n*-pentane and diethyl ether to precipitate 3-((2-chlorobenzyl)oxy)-5-(1-(piperidin-4-yl)-1*H*-pyrazol-4-yl)pyridine hydrochloride salt (91 mg, 222 µmol, 57.9%) as an off white solid. ^1^H NMR (400 MHz, DMSO-*d*_6_) δ 9.08 (m, 1H), 8.88 (m, 1H), 8.65 (s, 1H), 8.50 (s, 1H), 8.35 (s, 1H), 8.16 (s, 1H), 7.98 (s, 1H), 7.69-7.66 (m, 1H), 7.56-7.54 (m, 1H), 7.46-7.40 (m, 2H), 5.28 (s, 2H), 4.54-4.49 (m, 1H), 3.40 (d, 2H, *J* = 13 Hz), 3.14-3.04 (m, 2H), 2.25-2.07 (m, 4H); LRMS (ES+) m/z 369.13 [M+H]^+^; LC-MS purity 98.8% (Method A); Calculated [M+H]^+^ for C_20_H_21_ClN_4_O: 369.1482; Found: 369.1487.

The following compounds were made as HCl salts by deprotection of the Boc group following the method above:

### 3-((2,4-Dichlorobenzyl)oxy)-5-(1-(piperidin-4-yl)-1*H*-pyrazol-4-yl)pyridine. HCl (9) (TAD-0000025)

^1^H NMR (400 MHz, DMSO- *d*_6_) δ 8.92 (m, 1H), 8.73-8.71 (m, 1H), 8.65 (s, 1H), 8.50 (s, 1H), 8.35 (s, 1H), 8.17 (s, 1H), 8.00 (bs, 1H), 7.74-7.69 (m, 2H), 7.54 (dd, 1H*, J* = 8.4 Hz, 2.0 Hz), 5.32 (s, 2H), 4.51 (m, 1H), 3.42-3.39 (m, 2H), 3.15-3.06 (m, 2H), 2.27-2.05 (m, 4H). LRMS (ES+) m/z 403.10 [M+H]^+^; LC-MS purity 97.9% (Method A), Calculated [M+H]^+^ for C_20_H_20_Cl_2_N_4_O: 403.1092; Found: 403.1092.

### 3-((4-Bromobenzyl)oxy)-5-(1-(piperidin-4-yl)-1*H*-pyrazol-4-yl)pyridine. HCl (29) (TAD-0000030)

^1^H NMR (400 MHz, DMSO-*d*_6_) δ 9.05 (br s, 1H), 8.86 ( br s, 1H), 8.68 (s, 1H), 8.54 (s, 1H), 8.37 (s, 1H), 8.18 (s, 1H), 8.07 (s, 1H), 7.64 (d, 2H, *J =* 8.4 Hz), 7.47 (d, 2H, *J =* 8.4 Hz), 5.28 (s, 2H), 4.55-4.48 (m, 1H), 3.40 (d, 2H, *J* = 12.9 Hz), 3.14-3.06 (m, 2H), 2.26-2.10 (m, 4H); LRMS (ES+) m/z 413.09 [M+H]^+^; LC-MS purity 99.3% (Method B);

### 3-((4-Chlorobenzyl)oxy)-5-(1-(piperidin-4-yl)-1*H*-pyrazol-4-yl)pyridine. HCl (30) (TAD-0000031)

^1^H NMR (400 MHz, DMSO-*d*_6_) δ 8.94 (br s, 1H), 8.72 ( br s, 1H), 8.59 (s, 1H), 8.47 (s, 1H), 8.29 (s, 1H), 8.14 (s, 1H), 7.90 (s, 1H), 7.54-7.48 (m, 4H), 5.27 (s, 2H), 4.54-4.49 (m, 1H), 3.40 (d, 2H, *J* = 12.4 Hz), 3.14-3.06 (m, 2H), 2.26-2.09 (m, 4H); LRMS (ES+) m/z 369.16 [M+H]^+^; LC-MS purity 99.4% (Method A); Calculated [M+H]^+^ for C_20_H_21_ClN_4_O: 369.1482; Found: 369.1485.

### 3-((3-Chlorobenzyl)oxy)-5-(1-(piperidin-4-yl)-1*H*-pyrazol-4-yl)pyridine. HCl (31) (TAD-0411745)

^1^H NMR (400 MHz, DMSO-*d*_6_) δ 9.06 (br s, 1H), 8.88 ( br s, 1H), 8.70 (s, 1H), 8.55 (s, 1H), 8.40 (s, 1H), 8.19 (s, 1H), 8.11 (s, 1H), 7.60 (s, 1H), 7.48-7.43 (m, 3H), 5.32 (s, 2H), 4.56-4.49 (m, 1H), 3.61 (bs, 1H), 3.39 (d, 2H, *J* = 12.8 Hz), 3.14-3.06 (m, 2H), 2.32-2.10 (m, 4H); LRMS (ES+) m/z 369.15 [M+H]^+^; LC-MS purity 99.3% (Method A); Calculated [M+H]^+^ for C_20_H_21_ClN_4_O: 369.1482; Found: 369.1487.

### 3-((2,6-Dichlorobenzyl)oxy)-5-(1-(piperidin-4-yl)-1*H*-pyrazol-4-yl)pyridine. TFA (25) (TAD-0410899)

Trifluoroacetic acid (1 mL) was added drop wise to a stirred solution of *tert*-butyl 4-(4-(5-((2,6-dichlorobenzyl)oxy)pyridin-3-yl)-1*H*-pyrazol-1-yl)piperidine-1-carboxylate (230 mg, 457 µmol) in dichloromethane (5 mL) at 0°C and reaction mixture stirred at room temperature for 3h. After completion (TLC and LCMS monitoring), reaction mixture was concentrated and triturated with *n*-pentane and diethyl ether. Submitted to prep HPLC using TFA buffer to afford 3-((2,6-dichlorobenzyl)oxy)-5-(1-(piperidin-4-yl)-1*H*-pyrazol-4-yl)pyridine (130 mg, 238 µmol, 52.8%) as an off white trifluoroacetate salt. ^1^H NMR (400 MHz, DMSO- *d*_6_) δ 8.61 (m, 1H), 8.57 (d, 1H, *J* = 1.6 Hz), 8.43 (s, 1H), 8.37 (m, 1H), 8.26 (d, 1H, *J* = 2.8 Hz), 8.13 (s, 1H), 7.84 (m, 1H), 7.61-7.47 (m, 3H), 5.38 (s, 2H), 4.56-4.48 (m, 1H), 3.44-3.41 (m, 2H), 3.16-3.07 (m, 2H), 2.27-2.08 (m, 4H). LRMS (ES+) m/z 403.20 [M+H]^+^; LC-MS purity 97.4% (Method A); Calculated [M+H]^+^ for C_20_H_20_Cl_2_N_4_O: 403.1092; Found: 403.1093.

The following compounds were made as trifluoroacetate salts by deprotection of the Boc group following the method above:

### 3-(Benzyloxy)-5-(1-(piperidin-4-yl)-1*H*-pyrazol-4-yl)pyridine. TFA (26) (TAD-0410386)

^1^H NMR (400 MHz, DMSO-*d*_6_) δ 8.94 (bs, 1H), 8.72 (m, 1H), 8.49-8.40 (m, 2H), 8.19 (d, 1H, *J* = 2.8 Hz), 8.08 (s, 1H), 7.71 (s, 1H), 7.50-7.33 (m, 5H), 5.22 (s, 2H), 4.54-4.48 (m, 1H), 3.50-3.41 (m, 2H), 3.15-3.07 (m, 2H), 2.25-2.07 (m, 4H); LRMS (ES+) m/z 335.35 [M+H]^+^; LC-MS purity 99.2% (Method A). Calculated [M+H]^+^ for C_20_H_22_FN_4_O: 335.1872; Found: 335.1879.

### 3-((2-Methylbenzyl)oxy)-5-(1-(piperidin-4-yl)-1*H*-pyrazol-4-yl)pyridine. TFA (27) (TAD-0410410)

^1^H NMR (400 MHz, DMSO-*d*_6_) δ 8.66 (m, 1H), 8.50 (s, 1H), 8.40 (m, 2H), 8.21 (d, 1H, *J* = 2.4 Hz), 8.09 (s, 1H), 7.71 (m, 1H), 7.46-7.44 (m, 1H), 7.29-7.20 (m, 3H), 5.21 (s, 2H), 4.54-4.49 (m, 1H), 3.43-3.40 (m, 2H), 3.15-3.07 (m, 2H), 2.26 (s, 3H). 2.23-2.08 (m, 4H); LRMS (ES+) m/z 349.37 [M+H]^+^; LC-MS purity 94.1% (Method A); Calculated [M+H]^+^ for C_21_H_24_N_4_O: 349.2028; Found: 349.203.

### 3-((2-Fluorobenzyl)oxy)-5-(1-(piperidin-4-yl)-1*H*-pyrazol-4-yl)pyridine. TFA (28) (TAD-0410388)

^1^H NMR (400 MHz, DMSO-*d*_6_) δ 8.44 (d, 1H, *J* = 1.2 Hz), 8.33 (s, 1H), 8.15 (d, 1H, *J* = 2.4 Hz8.05 (s, 1H), 7.70-7.69 (m, 1H), 7.58-7.54 (m, 1H), 7.46-7.40 (m, 1H), 7.46-7.40 (m, 1H), 7.25-7.22 (m, 2H), 5.23 (s, 2H), 4.53-4.47 (m, 1H), 3.40 (d, 2H, *J* = 13.2 Hz), 3.14-3.02 (m, 2H), 2.25-2.06 (m, 4H); LRMS (ES+) m/z 353.3 [M+H]^+^; LC-MS purity 96.6% (Method A); Calculated [M+H]^+^ for C_20_H_21_FN_4_O: 353.1778; Found: 353.1781.

### 4-(4-(3-((2-Chlorobenzyl)oxy)phenyl)-1*H*-pyrazol-1-yl)piperidine. TFA (47) (TAD-0410390)

^1^H NMR (400 MHz, DMSO-*d*_6_) δ 8.64 (m, 1H), 8.40 (m, 1H), 8.27 (s, 1H), 7.97 (s, 1H), 7.64-7.62 (m, 1H), 7.55-7.51 (m, 1H), 7.43-7.38 (m, 2H), 7.31-7.20 (m, 3H), 6.87 (dd, 1H, *J* = 2.0, 7.8 Hz), 5.20 (s, 2H), 4.53-4.45 (m, 1H), 3.43-3.40 (m, 2H), 3.15-3.06 (m, 2H), 2.25--2.07 (m, 4H); LRMS (ES+) m/z 368.30 [M+H]^+^; LC-MS purity 97.23% (Method A). Calculated [M+H]^+^ for C_20_H_21_ClN_4_O: 368.1530; Found: 368.1531.

### 4-((2-Chlorobenzyl)oxy)-2-(1-(piperidin-4-yl)-1*H*-pyrazol-4-yl)pyridine. AcOH (48) (TAD-0410623)

This compound was prepared by protection of the Boc group following the method above but submitted to prep HPLC using AcOH buffer to afford 4-((2-chlorobenzyl)oxy)-2-(1-(piperidin-4-yl)-1*H*-pyrazol-4-yl)pyridine (29 mg, 78.6 µmol, 24.6%) as a white acetate salt. ^1^H NMR (400 MHz, DMSO-*d*_6_) δ 8.35 (s, 1H), 8.33 (d, 1H, *J* = 5.6 Hz), 8.02 (s, 1H), 7.66-7.64 (m, 1H), 7.56-7.54(m, 1H), 7.45-7.38 (m, 3H), 6.86 (dd, 1H, *J* = 2.6, 5.8 Hz), 5.28 (s, 2H), 4.24-4.18 (m, 1H), 3.06-3.02 (m, 2H), 2.62-2.50 (m, 2H), 1.99-1.97 (m, 2H), 1.87 (s, 3H), 1.84-1.74 (m, 2H); LRMS (ES+) m/z 369.25 [M+H]^+^; LC-MS purity 99.93% (Method A). Calculated [M+H]^+^ for C_20_H_21_ClN_4_O: 369.1482; Found: 369.1484.

### 2-((2-Chlorobenzyl)oxy)-6-(1-(piperidin-4-yl)-1*H*-pyrazol-4-yl)pyridine. TFA (49) (TAD-0410597)

^1^H NMR (400 MHz, DMSO-*d*_6_) δ 8.64 (m, 1H), 8.34 (m, 1H), 8.30 (s, 1H), 8.04 (s, 1H), 7.71 (t, 2H*, J* = 7.8Hz), 7.62-7.59 (m, 1H), 7.53-7.50 (m, 1H), 7.38-7.36 (m, 3H), 7.29 (d, 1H, *J* = 7.2 Hz), 6.70 (d, 1H, *J* = 8 Hz), 5.51 (s, 2H), 4.58-4.50 (m, 1H), 3.45-3.41 (m, 2H), 3.13-3.05 (m, 2H), 2.32--2.08 (m, 4H); LRMS (ES+) m/z 369.15 [M+H]^+^; LC-MS purity 98.74% (Method A). Calculated [M+H]^+^ for C_20_H_21_ClN_4_O: 369.1482; Found: 369.1484.

### 3-((2-Chlorobenzyl)oxy)-5-(1-(1-methylpiperidin-4-yl)-1*H*-pyrazol-4-yl)pyridine (41) (TAD-0410749)

Lithium aluminum hydride solution 2.0 M in THF (0.6 mL) was added dropwise to a stirred solution of *tert*-butyl 4-(4-(5-((2-chlorobenzyl)oxy)pyridin-3-yl)-1*H*-pyrazol-1-yl)piperidine-1-carboxylate (**24,** 180 mg, 384 µmol) in tetrahydrofuran (5 mL, 61.4 µmol) at 0°C. The reaction mixture was warmed to room temperature and then heated at 50°C for 16h. After completion (TLC & LCMS monitoring), the reaction mixture was quenched with ice water, diluted with water, extracted with 20% MeOH in DCM, washed with water, brine, dried over anhydrous sodium sulphate concentrated. The crude material was purified by flash column chromatography using 20% MeOH in DCM as an eluent, and the desired fractions were concentrated to afford 3-((2-chlorophenyl)methoxy)-5-(1-(1-methylpiperidin-4-yl)-1*H*-pyrazol-4-yl)pyridine (59 mg, 153 µmol, 40%). ^1^H NMR (400 MHz, DMSO-*d*_6_) δ 8.49 (d, 1H, *J* = 1.6 Hz), 8.47 (s, 1H), 8.18 (d, 1H, *J* = 2.4Hz), 8.02 (s, 1H), 7.73-7.1 (m, 1H), 7.67-7.65 (m, 1H), 7.55-7.53 (m, 1H), 7.44-7.41 (m, 2H), 5.27 (s, 2H), 4.16-4.08 (m, 1H), 2.85 (d, 2H, *J* = 11.2 Hz), 2.21 (s, 3H), 2.08-1.92 (m, 6H); LRMS (ES+) m/z 383.20 [M+H]^+^; LC-MS purity 99.7% (Method A); Calculated [M+H]^+^ for C_21_H_23_ClN_4_O: 383.1639; Found: 383.1640.

### 3-((2-Chlorobenzyl)oxy)-5-(1-(1-(3,3-dimethylbutyl)piperidin-4-yl)-1*H*-pyrazol-4-yl)pyridine (42) (TAD-0410679)

To a stirred solution of 3-[(2-chlorophenyl)methoxy]-5-[1-(piperidin-4-yl)-1*H*-pyrazol-4-yl]pyridine (150 mg, 407 µmol) and 1-bromo-3,3-dimethylbutane (67.1 mg, 407 µmol) in dimethylformamide (3 mL, 38.7 µmol), was added potassium carbonate (169 mg, 3 eq., 1.22 µmol) portion wise and reaction mixture heated at 100°C for 16h. After completion (TLC and LCMS monitoring). Reaction mixture was diluted with water, extracted with EtOAc, washed with brine, dried over anhydrous sodium sulfate, filtered and concentrated. Crude was purified by flash chromatography using 40% EtOAc in hexane as an eluent, desired fractions were concentrated to afford 3-((2-chlorobenzyl)oxy)-5-(1-(1-(3,3-dimethylbutyl)piperidin-4-yl)-1*H*-pyrazol-4-yl)pyridine (15 mg, 33.1 µmol, 8.1%) as an off white solid. ^1^H NMR (400 MHz, DMSO-*d*_6_) δ 8.49 (d, 1H, *J* = 1.2 Hz), 8.43 (s, 1H), 8.18 (d, 1H, *J* = 2.4Hz), 8.03 (s, 1H), 7.73-7.2 (m, 1H), 7.67-7.65 (m, 1H), 7.55-7.53 (m, 1H), 7.45-7.41 (m, 2H), 5.27 (s, 2H), 4.15 (m, 1H), 2.99-2.96 (m, 3H), 2.10-1.92 (m, 6H), 1.39-1.33 (m, 2H), 0.897 (s, 9H); LRMS (ES+) m/z 453.35 [M+H]^+^; LC-MS purity 98% (Method A); Calculated [M+H]^+^ for C_26_H_33_ClN_4_O: 453.2421; Found: 453.2419.

### 1-(4-(4-(5-((2-Chlorobenzyl)oxy)pyridin-3-yl)-1*H*-pyrazol-1-yl)piperidin-1-yl)ethan-1-one (43) (TAD-0409937)

To the stirred solution of 3-((2-chlorophenyl)methoxy)-5-(1-(piperidin-4-yl)-1*H*-pyrazol-4-yl)pyridine (0.5 g, 1.36 µmol) in dichloromethane (5 mL) was added dropwise triethylamine (588 µL, 3 eq., 4.07 µmol) at 0°C followed by acetyl chloride (128 mg, 1.2 eq., 1.63 µmol). Reaction mixture was stirred at RT for 16h. After completion (TLC & LCMS monitoring), reaction mixture was diluted with chilled water, extracted with DCM. Organic layer was washed with water, brine, dried over anhydrous sodium sulfate, concentrated to afford crude. Crude was purified by flash column chromatography using 50% EtOAc in *n*-heptane, desired fractions were concentrated to afford 1-(4-(4-(5-((2-chlorobenzyl)oxy)pyridin-3-yl)-1*H*-pyrazol-1-yl)piperidin-1-yl)ethan-1-one (166 mg, 392 µmol, 28.9%) as brown solid. ^1^H NMR (400 MHz, DMSO-*d*_6_) δ 8.49 (m, 1H), 8.44 (s, 1H), 8.19 (d, 1H, *J* = 2.4Hz), 8.04 (s, 1H), 7.73 (m, 1H), 7.67-7.65 (m, 1H), 7.55-7.53 (m, 1H), 7.45-7.39 (m, 2H), 5.27 (s, 2H), 4.48-4.42 (m, 2H), 3.94-3.91 (m, 1H), 3.34-3.19 (m, 1H), 2.77-2.71 (m, 1H); 2.12-2.05 (m, 2H) 1.92 (s, 3H), 1.92-1.71 (m, 2H); LRMS (ES+) m/z 411.15 [M+H]^+^; LC-MS purity 97.7% (Method A), Calculated [M+H]^+^ for C_22_H_23_ClN_4_O_2_: 411.1588; Found: 411.1586.

The following compound was synthesized following the method described above:

### 1-(4-(4-(5-((2-Chlorobenzyl)oxy)pyridin-3-yl)-1*H*-pyrazol-1-yl)piperidin-1-yl)-3,3-dimethylbutan-1-one (44) (TAD-0410391)

^1^H NMR (400 MHz, DMSO-*d*_6_) δ 8.49 (br s, 1H), 8.43 (s, 1H), 8.18 (d, 1H, *J* = 2.8 Hz), 8.03 (s, 1H), 7.72 (m, 1H), 7.67-7.65 (m, 1H), 7.55-7.53 (m, 1H), 7.43-7.41 (m, 2H), 5.27 (s, 2H), 4.57-4.54(m, 1H),4.44-4.41 (m, 1H), 4.11-4.08 (m, 1H), 3.20-3.17 (m, 1H), 2.75-2.72 (m, 1H); 2.3-2.24 (m, 2H), 2.08 (m, 2H), 1.78 (m, 2H). 1.00 (s, 9H); LRMS (ES+) m/z 467.37 [M+H]^+^; LC-MS purity 97.36% (Method A); Calculated [M+H]^+^ for C_26_H_31_ClN_4_O_2_: 467.2214; Found: 467.2214.

### Methyl 4-(4-(5-((2-chlorobenzyl)oxy)pyridin-3-yl)-1*H*-pyrazol-1-yl)piperidine-1-carboxylate (45) (TAD-0410392)

To the stirred solution of 3-((2-chlorophenyl)methoxy)-5-(1-(piperidin-4-yl)-1*H*-pyrazol-4-yl)pyridine (150 mg, 407 µmol) in dichloromethane (3 mL, 46.9 µmol) was added triethylamine (170 µL, 3 eq., 1.22 µmol) followed by methyl chloroformate (57.6 mg, 1.5 eq., 610 µmol) at 0°C. Reaction mixture was stirred at RT for 16h. After completion TLC & LCMS monitoring, reaction mixture was diluted with water, extracted with DCM. Organic layer was washed with brine, dried over anhydrous Na_2_S0_4_, concentrated. Crude was purified using 10% MeOH in DCM as an eluent, desired fractions were concentrated to afford methyl 4-(4-(5-((2-chlorobenzyl)oxy)pyridin-3-yl)-1*H*-pyrazol-1-yl)piperidine-1-carboxylate (53 mg, 118 µmol, 29.0%) as yellowish semi solid. ^1^H NMR (400 MHz, DMSO-*d*_6_) δ 8.49 (s, 1H), 8.44 (s, 1H), 8.18 (s, 1H), 8.03 (s, 1H), 7.72 (s, 1H), 7.65 (m, 1H), 7.53 (m, 1H), 7.43-7.40 (m, 2H), 5.27 (s, 2H), 4.44-4.41 (m, 1H), 4.08-4.07 (m, 2H), 3.62 (s, 3H), 3.01 (m, 2H), 2.08-2.05 (m, 2H), 1.86-1.81 (m, 2H); LRMS (ES+) m/z 427.30 [M+H]^+^; LC-MS purity 95.1% (Method A); Calculated [M+H]^+^ for C_22_H_23_ClN_4_O_3_: 427.1537; Found: 427.1537.

### Benzyl 4-(4-(5-((2-chlorobenzyl)oxy)pyridin-3-yl)-1*H*-pyrazol-1-yl)piperidine-1-carboxylate (46) (TAD-0410616)

To a stirred solution of 3-((2-chlorobenzyl)oxy)-5-(1-(piperidin-4-yl)-1*H*-pyrazol-4-yl)pyridine hydrochloride salt (75 mg, 170 µmol) in tetrahydrofuran (2 mL, 24.6 µmol) was added benzyl chloroformate (29 µL, 1.2 eq., 204 µmol) and *N,N*-diisopropylethylamine (149 µL, 5 eq., 856 µmol) at room temperature. The reaction mixture was stirred for 1.5 hours at which time LCMS indicated the reaction was complete. The reaction mixture was diluted with the water (2 mL) and extracted with ethyl acetate (3 times X 2 mL). The combined organic layers were dried over anhydrous sodium sulfate, filtered and concentrated under reduced pressure to obtain the crude compound. The crude material was purified by flash silica gel column chromatography using 0-85% EtOAc in *n*-hexanes to afford the desired product benzyl 4-(4-(5-((2-chlorobenzyl)oxy)pyridin-3-yl)-1*H*-pyrazol-1-yl)piperidine-1-carboxylate (32 mg, 63.6 µmol, 37%) as a yellow oil. ^1^H NMR (400 MHz, DMSO-*d*_6_) δ 8.53 (s, 1H), 8.47 (s, 1H), 8.24 (s, 1H), 8.06 (s, 1H), 7.79 (s, 1H), 7.70-7.65 (m, 1H), 7.58-7.53 (m, 1H), 7.47-7.41 (m, 2H), 7.41-7.36 (m, 4H), 7.36-7.30 (m, 1H), 5.29 (s, 2H), 5.11 (s, 2H), 4.48-4.39 (m, 1H), 4.13 (d, 2H, *J* = 13.0 Hz), 3.16-2.93 (m, 2H), 2.09 (d, 2H, *J* = 12.0 Hz), 1.91-1.79 (m, 2H). LRMS (ES+) m/z 503 [M+H]^+^; LC-MS purity 94.80% (Method C). Calculated [M+H]^+^ for C_28_H_27_ClN_4_O_3_: 503.1580; Found: 503.1847.

### *tert*-Butyl (*S*)-4-(4-(5-(1-(2-chlorophenyl)ethoxy)pyridin-3-yl)-1*H*-pyrazol-1-yl)piperidine-1-carboxylate (intermediate)

To a stirred solution of *tert*-butyl 4-(4-(5-hydroxypyridin-3-yl)-1*H*-pyrazol-1-yl)piperidine-1-carboxylate (**16,** 0.5 g, 1.52 µmol) and (*R*)-1-(*o*-chlorophenyl)-1-ethanol (238 mg, 1.52 µmol) in tetrahydrofuran (20 mL, 246 µmol), was added triphenylphosphine (799 mg, 2 eq., 3.04 µmol) portion wise and reaction mixture stirred at room temperature for 1h. Then DEAD (530 mg, 2 eq., 3.04 µmol) was added in stock solution of tetrahydrofuran dropwise and reaction mixture stirred at room temperature for 16h. After completion (TLC and LCMS monitoring). Reaction mixture was diluted with water, extracted with EtOAc, washed with brine, dried over anhydrous sodium sulfate, filtered and concentrated. Crude was purified by flash chromatography using 70% EtOAc in hexane as an eluent, desired fractions were concentrated to afford *tert*-butyl (*S*)-4-(4-(5-(1-(2-chlorophenyl)ethoxy)pyridin-3-yl)-1*H*-pyrazol-1-yl)piperidine-1-carboxylate (**33,**180 mg, 193 µmol, 12.2%) as a sticky brown solid. ^1^H NMR (400 MHz, DMSO-*d*_6_) δ 8.40 (s, 1H), 8.39 (m, 1H) 8.35 (s, 1H), 7.94 (d, 1H, *J* = 2.4 Hz), 7.92 (s, 1H), 7.55-7.48 (m, 3H), 7.37-7.30 (m, 2H), 5.88 (q, 1H, *J* = 6.4Hz), 4.36 (m, 1H), 4.05-4.02 (m, 2H), 2.93-2.92 (m, 2H), 2.04-2.01 (m, 2H), 1.83-1.79 (m, 2H), 1.63 (d, 3H, *J* = 6.4 Hz), 1.62 (s, 9H); LRMS (ES+) m/z 483.0 [M+H]^+^; LC-MS purity 99.7% (Method A), 90.2% ee.

### (S)-3-(1-(2-Chlorophenyl)ethoxy)-5-(1-(piperidin-4-yl)-1*H*-pyrazol-4-yl)pyridine. TFA (37) (TAD-0411448)

Trifluoroacetic acid (1 mL) drop wise to a stirred solution of *tert*-butyl (*S*)-4-(4-(5-(1-(2-chlorophenyl)ethoxy)pyridin-3-yl)-1*H*-pyrazol-1-yl)piperidine-1-carboxylate (180 mg, 373 µmol) in dichloromethane (4 mL) at 0°C and reaction mixture stirred at room temperature for 3h. After completion (TLC and LCMS monitoring), reaction mixture was concentrated and triturated with *n*-pentane and diethyl ether. Submitted to prep HPLC using TFA buffer to afford (*S*)-3-(1-(2-chlorophenyl)ethoxy)-5-(1-(piperidin-4-yl)-1*H*-pyrazol-4-yl)pyridine (45 mg, 89.7 µmol, 24.1%) as an off white trifluoroacetate salt.^1^H NMR (400 MHz, DMSO-*d*_6_) δ 8.62 (m, 1H), 8.45 (d, 1H, *J* = 1.6 Hz), 8.39 (m, 1H) 8.33 (s, 1H), 8.00 (s, 1H), 7.99 (d, 1H, *J* = 2.4 Hz), 7.59-7.58 (m, 1H), 7.55-7.48 (m, 2H), 7.38-7.30 (m, 2H), 5.89 (q, 1H, *J* = 6.4Hz), 4.53-4.48 (m, 1H), 3.43-3.40 (m, 2H), 3.14-3.06 (m, 2H), 2.23-2.06 (m, 4H), 1.63 (d, 3H, *J* = 6.4 LRMS (ES+) m/z 383.3 [M+H]^+^; LC-MS purity 99.4% (Method A). Calculated [M+H]^+^ for C_21_H_23_ClN_4_O: 383.1639; Found: 383.1642.

The following compounds were synthesized by a Mitsunobu reaction followed by TFA deprotection of the Boc intermediate in a similar fashion to compound **37**.

### (*R*)-3-(1-(2-Chlorophenyl)ethoxy)-5-(1-(piperidin-4-yl)-1*H*-pyrazol-4-yl)pyridine. TFA (38) (TAD-0411473)

^1^H NMR (400 MHz, DMSO-*d*_6_) δ 8.64 (m 1H), 8.45 (d, 1H, *J* = 1.6Hz), 8.39 (m, 1H) 8.34 (s, 1H), 8.00 (m, 2H), 7.62 (m, 1H), 7.55-7.49 (m, 2H), 7.38-7.31 (n, 2H), 5.90 (q, 1H, *J* = 6.4 Hz), 4.53-4.48 (m, 1H), 3.43-3.41 (m, 2H), 3.14-3.05 (m, 2H), 2.24-2.05 (m, 4H), 1.63 (d, 3H, *J* = 6.4 Hz); LRMS (ES+) m/z 383.3 [M+H]^+^; LC-MS purity 98.5% (Method A). Calculated [M+H]^+^ for C_21_H_23_ClN_4_O: 383.1639; Found: 383.1643.

### (*R*)-3-(1-(4-Chlorophenyl)ethoxy)-5-(1-(piperidin-4-yl)-1*H*-pyrazol-4-yl)pyridine (40) (TAD-0000082)

The crude TFA salt was further purified by prep HPLC to afford (*R*)-3-(1-(4-chlorophenyl)ethoxy)-5-(1-(piperidin-4-yl)-1*H*-pyrazol-4-yl)pyridine as an off-white free base. ^1^H NMR (400 MHz, DMSO-*d*_6_) δ 8.70 (br s, 1H), 8.41 (d, 1H, *J* = 1.6 Hz), 8.06 (d, 1H, *J* = 2.4 Hz), 8.02 (s, 1H), 7.59-7.58 (m, 1H), 7.49-7.41 (m, 4H), 5.72 (q, 1H, *J* = 6.3Hz), 4.52-4.46 (m, 1H), 3.43-3.40 (m, 2H). 3.10 (m, 2H), 2.21-2.06 (m, 4H), 1.58 (d, 3H, *J* = 6.4 Hz); LRMS (ES+) m/z 383.24 [M+H]^+^; LC-MS purity 98.2% (Method A). Calculated [M+H]^+^ for C_21_H_23_ClN_4_O: 383.1639; Found: 383.1639.

### (*S*)-3-(1-(4-Chlorophenyl)ethoxy)-5-(1-(piperidin-4-yl)-1*H*-pyrazol-4-yl)pyridine (39) (TAD-0422576)

The crude TFA salt was purified by prep HPLC to afford (*S*)-3-(1-(4-chlorophenyl)ethoxy)-5-(1-(piperidin-4-yl)-1*H*-pyrazol-4-yl)pyridine as an off-white free base (30mg, 76umol, 36.7%). ^1^H NMR (400 MHz, DMSO-*d*_6_) δ 8.38 (d, 1H, *J* = 1.7 Hz), 8.31 (s, 1H) 8.03 (d, 1H, *J* = 2.7 Hz), 7.93 (s, 1H), 7.57-7.56 (m, 1H), 7.49-7.41 (m, 4H), 5.71 (q, 1H, *J* = 6.3Hz), 4.21-4.13 (m, 1H), 3.05-3.01 (m, 2H), 2.57-2.54 (m, 2H), 1.97-1.72 (m, 4H), 1.57 (d, 3H, *J* = 6.4 Hz); LRMS (ES+) m/z 383.15 [M+H]^+^; LC-MS purity 99.9% (Method A).

### 2-((2-Chlorobenzyl)oxy)-4-iodopyridine (52)

Sodium hydride (295 mg, 1.5 eq., 7.36 µmol) was added portion wise to a stirred solution of (2-chlorophenyl)methanol (**50,** 0.7 g, 4.91 µmol) in DMF (8 mL) at 0°C and reaction mixture stirred at 0°C for 30 min. 2-chloro-4-iodopyridine (**51**, 1.41 g, 1.2 eq., 5.89 µmol) was then added and reaction mixture was stirred at RT for 16h. After completion (TLC and LCMS monitoring), reaction mixture was diluted with water, extracted with EtOAc, washed with brine, dried over anhydrous sodium sulfate, filtered and concentrated. The crude material was purified by flash chromatography using 50% EtOAc in hexane as an eluent, desired fractions were concentrated to afford 2-((2-chlorobenzyl)oxy)-4-iodopyridine (250 mg, 716 µmol, 14.6%) as white solid. ^1^H NMR (400 MHz, DMSO-*d*_6_) δ 7.93 (d, 1H, *J* = 5.7Hz), 7.57-7.55 (m, 1H), 7.52-7.50 (m, 1H), 7.42-7.37 (m, 4H), 5.41 (s, 2H); LRMS (ES+) m/z 345.70 [M+H]^+^.

### *tert*-Butyl 4-(4-(2-((2-chlorobenzyl)oxy)pyridin-4-yl)-1*H*-pyrazol-1-yl)piperidine-1-carboxylate

Potassium carbonate dipotassium carbonate (160 mg, 2 eq., 1.16 µmol) was added to a stirred solution of 2-((2-chlorobenzyl)oxy)-4-iodopyridine (**52,** 0.2 g, 579 µmol) and *tert*-butyl 4-(4-(4,4,5,5-tetramethyl-1,3,2-dioxaborolan-2-yl)-1*H*-pyrazol-1-yl)piperidine-1-carboxylate (**15,** 218 mg, 579 µmol) in toluene (5.65 mL, 47.7 µmol), ethanol (5.65 mL, 96.7 µmol) and water (0.2 mL) and mixture was purged with nitrogen for 20 minutes. [1,1’-Bis(diphenylphosphino)ferrocene]palladium(II) dichloride (23.6 mg, 0.05 eq., 28.9 µmol) was then added and reaction mixture was heated at 90°C and stirred for 16h. After completion of the reaction (TLC and LCMS monitoring), the reaction mixture was concentrated. The crude was purified by flash chromatography using 30% EtOAc in hexane as an eluent, desired fractions were concentrated to afford *tert*-butyl 4-(4-(2-((2-chlorobenzyl)oxy)pyridin-4-yl)-1*H*-pyrazol-1-yl)piperidine-1-carboxylate (0.2 g, 422 µmol, 72.9%) as an light brown semi solid. ^1^H NMR (400 MHz, DMSO-*d*_6_) δ 8.51 (s, 1H), 8.11-8.08 (m, 2H), 7.58-7.56 (m, 1H), 7.52-7.50 (m, 1H), 7.40-7.36 (m, 2H), 7.24 (dd, 1H, *J* = 1.4, 5.4 Hz), 7.15 (s, 1H), 5.43 (s, 2H), 4.41-4.34 (m, 1H), 4.04 (m, 2H), 2.92 (br s, 2H), 2.04 (m, 2H), 1.79 (qd, 2H, *J =* 4.2, 12.2 Hz), 1.42 (s, 9H); LRMS (ES+) m/z 469.40 [M+H]^+^; LC-MS purity 99.4% (Method A).

### 2-((2-Chlorobenzyl)oxy)-4-(1-(piperidin-4-yl)-1*H*-pyrazol-4-yl)pyridine (53) (TAD-0422881)

Oxalyl chloride (68.6 µL, 3 eq., 0.8 µmol) was added to a stirred solution of *tert*-butyl 4-(4-(2-((2-chlorobenzyl)oxy)pyridin-4-yl)-1*H*-pyrazol-1-yl)piperidine-1-carboxylate (125 mg, 267 µmol) in methanol (5.98 mL, 148 µmol) at 0°C. The reaction mixture was stirred at RT for 3h. After completion (TLC & LCMS monitoring), the reaction mixture was concentrated, washed with pentane and diethyl ether, dried and concentrated. The crude material was purified by prep HPLC to afford 2-((2-chlorobenzyl)oxy)-4-(1-(piperidin-4-yl)-1*H*-pyrazol-4-yl)pyridine (59 mg, 158 µmol, 59.4%) as brown semi solid. ^1^H NMR (400 MHz, DMSO-*d*_6_) δ 8.35 (s, 1H), 8.33 (d, 1H, *J* = 5.8 Hz), 8.02 (s, 1H), 7.66-7.64 (m, 1H), 7.56-7.54 (m, 1H), 7.45-7.38 (m, 3H), 6.86 (dd, 1H, *J* = 2.6, 5.8 Hz), 5.22 (s, 2H), 4.24-4.18 (m, 1H), 3.05-3.02 (m, 2H), 2.59-2.56 (m, 2H), 1.99-1.96 (m, 2H), 1.84-1.74 (m, 2H); LRMS (ES+) m/z 369.25 [M+H]^+^; LC-MS purity 99.9% (Method A). Calculated [M+H]^+^ for C_20_H_21_ClN_4_O: 369.1482; Found: 369.1487.

### 3-Bromo-5-((2-chlorobenzyl)oxy)pyridine (54)

2-chlorobenzylbromide (2.46 mL, 1.1 eq., 19 µmol) was added dropwise to a stirred solution of 5-bromo-3-pyridinol (**14,** 3 g, 17.2 µmol) in dimethylformamide (37.5 mL, 484 µmol) and cesium carbonate (11.2 g, 2 eq., 34.5 µmol) 0°C and reaction mixture was stirred at RT for 16h. After completion of the reaction (TLC monitoring), reaction mixture was diluted with ice cold water, and extracted with EtOAc, washed with water, brine, dried over anhydrous sodium sulfate, filtered and concentrated. The crude material was purified by flash chromatography using 0-30% EtOAc in *n*-heptane as an eluent, desired fractions were concentrated to afford 3-bromo-5-((2-chlorobenzyl)oxy)pyridine (3.2 g, 10.7 µmol, 62.2%) as a brown solid. ^1^H NMR (400 MHz, DMSO-*d*_6_) δ 8.33 (m, 2H), 7.53-7.47 (m, 2H), 7.45-7.41 (m, 1H), 7.34-7.29 (m, 2H), 5.20 (s, 2H).

### 3-((2-Chlorobenzyl)oxy)-5-((trimethylsilyl)ethynyl)pyridine

Triethylamine (2.82 mL, 3 eq., 20.1 µmol) and ethynyltrimethylsilane (2.86 mL, 3 eq., 20.1 µmol) were added to a stirred solution of 3-bromo-5-((2-chlorobenzyl)oxy)pyridine (**54,** 2 g, 6.7 µmol) in toluene (40 mL, 376 µmol) and reaction mixture was purged with argon for 15 minutes. Copper iodide (319 mg, 0.25 eq., 1.67 µmol) and palladium—triphenylphosphine (1/4) (387 mg, 0.05 eq., 335 µmol) were then added to the reaction mixture at 0°C which was then heated at 80°C for 16h. After completion of the reaction (TLC and LCMS monitoring), reaction mixture was diluted with ice cold water, and extracted with EtOAc, washed with water, brine, dried over anhydrous sodium sulfate, filtered and concentrated. The crude material was purified by flash chromatography using 0-20% EtOAc in *n*-heptane as an eluent, desired fractions were concentrated to afford 3-((2-chlorobenzyl)oxy)-5-((trimethylsilyl)ethynyl)pyridine (1.5 g, 4.75 µmol) as light brown solid. ^1^H NMR (400 MHz, DMSO-*d*_6_) δ 8.34 (m, 2H), 7.55-7.53 (m, 1H), 7.47-7.32 (m, 4H), 5.21 (s, 2H), 0.25 (s, 9H).

### 3-((2-Chlorobenzyl)oxy)-5-ethynylpyridine (59)

Potassium carbonate (1.71 g, 3 eq., 12.3 µmol) was added to a stirred solution of 3-((2-chlorobenzyl)oxy)-5-((trimethylsilyl)ethynyl)pyridine (1.3 g, 4.12 µmol) in methanol (26 mL, 642 µmol) under a N_2_ atmosphere and the reaction mixture stirred at RT for 16h. After completion of the reaction (TLC and LCMS monitoring), reaction mixture was diluted with ice cold water, and extracted with EtOAc, washed with water, brine, dried over anhydrous sodium sulfate, filtered and concentrated. The crude material was purified by flash chromatography using 0-30% EtOAc in *n*-heptane as an eluent, desired fractions were concentrated to afford 3-((2-chlorobenzyl)oxy)-5-ethynylpyridine (1g, 2.87 µmol, 69.8%) as a brown solid. ^1^H NMR (400 MHz, DMSO-*d*_6_) δ 8.38 (s, 1H), 8.34 (s, 1H), 7.55-7.52 (m, 1H), 7.48-7.31 (m, 4H), 5.22 (s, 2H), 3.22 (s, 1H).

### *tert*-Butyl 4-(4-(5-((2-chlorobenzyl)oxy)pyridin-3-yl)-1*H*-1,2,3-triazol-1-yl)piperidine-1-carboxylate

Sodium ascorbate (203 mg, 0.25 eq., 1.03 µmol) and copper sulfate pentahydrate (205 mg, 0.2 eq., 821 µmol) were added to a stirred solution of 3-((2-chlorobenzyl)oxy)-5-ethynylpyridine (**59,** 1 g, 4.1 µmol) and tert-butyl 4-azido-1-piperidinecarboxylate (**60,** 1.11 g, 1.2 eq., 4.92 µmol) in t-butanol (20 mL, 209 µmol) and water (20 mL, 1.11 mol) under a N_2_ atmosphere. The reaction mixture was stirred at RT for 16h. After completion of the reaction (TLC and LCMS monitoring), reaction mixture was diluted with ice cold water, and extracted with EtOAc, washed with water, brine, dried over anhydrous sodium sulfate, filtered and concentrated. The crude material was purified by flash chromatography using 0-100% EtOAc in *n*-heptane as an eluent, desired fractions were concentrated to afford *tert*-butyl 4-(4-(5-((2-chlorobenzyl)oxy)pyridin-3-yl)-1*H*-1,2,3-triazol-1-yl)piperidine-1-carboxylate (1.1 g, 2.22 µmol, 54.2%) as an off white solid. ^1^H NMR (400 MHz, DMSO-*d*_6_) δ 8.88 (s, 1H), 8.69 (s, 1H, *J* = 1.6Hz), 8.35 (d, 1H, *J =* 2.8 Hz), 7.90-7.89 (m, 1H), 7.67-7.65 (m, 1H), 7.55-7.54 (m, 1H), 7.45-7.42 (m, 2H), 5.31 (s, 2H), 4.84-4.76 (m, 1H), 4.07 (m, 2H), 2.99 (br s, 2H), 2.15-2.12 (m, 2H), 1.79 (qd, 2H, *J =* 4.2, 12.1 Hz), 1.43 (s, 9H); LRMS (ES+) m/z 470.3 [M+H]^+^; LC-MS purity 99.4% (Method A).

### 3-((2-Chlorobenzyl)oxy)-5-(1-(piperidin-4-yl)-1*H*-1,2,3-triazol-4-yl)pyridine. TFA (62) (TAD-0411690)

*Tert*-butyl 4-(4-(5-((2-chlorobenzyl)oxy)pyridin-3-yl)-1*H*-1,2,3-triazol-1-yl)piperidine-1-carboxylate was deprotected as described earlier to afford 3-((2-chlorobenzyl)oxy)-5-(1-(piperidin-4-yl)-1H-1,2,3-triazol-4-yl)pyridine as a light brown trifluoroacetate salt (130mg, 269 µmol, 50.5%) ^1^H NMR (400 MHz, DMSO-*d*_6_) δ 8.83 (s, 1H), 8.79 (m, 1H), 8.73 (d, 1H, *J* = 1.6 Hz), 8.54 (m, 1H), 8.37 (d, 1H, *J* = 2.8 Hz), 7.93-7.92 (m, 1H), 7.68-7.64 (m, 1H), 7.56-7.53 (m, 1H), 7.46-7.43 (m, 2H), 5.32 (s, 2H), 4.92-4.87 (m, 1H), 3.48-3.44 (m, 2H), 3.18-3.15 (m, 2H), 2.24-2.13 (m, 4H); ^19^F NMR (400 MHz, DMSO-*d*_6_) δ -73.49; LRMS (ES+) m/z 370.30 [M+H]^+^; LC-MS purity 99.53% (Method A). Calculated [M+H]^+^ for C_19_H_20_ClN_5_O: 370.1435; Found: 370.1435.

### *tert*-Butyl 4-(1-(5-((2-chlorobenzyl)oxy)pyridin-3-yl)-1*H*-pyrazol-4-yl)piperidine-1-carboxylate (intermediate) (TAD-0410527)

To a reaction vessel was added tert-butyl 4-(IH-pyrazo)-4-yl)piperidine-l-carboxylate (100 mg, 398 µmol), 3-bromo-5-((2-chlorobenzyl)oxy)pyridine (238 mg, 796 µmol), copper(II) iodide (11.4 mg, 59.7 µmol), potassium carbonate (137 mg, 995 µmol), potassium phosphate (135 mg, 637 µmol), acetonitrile (4 ml, 76.5 µmol), and (1*R*,2*R*)-*N*^1^,*N*^2^-dimethylcyclohexane-1,2-diamine-dimethylcyclohexane-1,2-diamine (62.7 µM, 398 µmol). The vessel was purged with nitrogen, sealed and heated 120 °C for 5 hrs. After the reaction was complete (monitored by LCMS), the mixture was cooled to room temperature and diluted with saturated aqueous ammonium chloride (3 mL). This mixture was extracted with ethyl acetate (3 x 3 mL). The combined extracts were washed with water (3 x 1 mL), dried over anhydrous sodium sulfate, filtered and concentrated. The residue was purified by flash chromatography using a gradient of 0% to 40% ethyl acetate in hexane as an eluent. Desired fractions were concentrated to afford *tert*-butyl 4-(1-(5-((2-chlorobenzyl)oxy)pyridin-3-yl)-1*H*-pyrazol-4-yl)piperidine-1-carboxylate (169 mg, 360 µmol, 90.6%) as a yellow oil. ^1^H NMR (400 MHz, DMSO-*d*_6_) δ 8.73 (s, 1H), 8.51 (s, 1H), 8.31 (s, 1H), 7.93 (s, 1H), 7.76 (s, 1H), 7.68 (d, 1H, *J* = 6.4 Hz, 1H), 7.56 (d, 1H, *J* = 7.8 Hz, 1H), 7.47-7.40 (m, 2H), 5.33 (s, 2H), 4.01 (d, 2H, *J* = 9.7 Hz), 2.96-2.76 (m, 2H), 2.73 (t, 1H, *J* = 11.6 Hz), 1.91 (d, 2H, *J* = 13.1 Hz), 1.50-1.37 (m, 11H). LRMS (ES+) m/z 469 [M+H]^+^; LC-MS purity 94.71% (Method C).

### 3-((2-Chlorobenzyl)oxy)-5-(4-(piperidin-4-yl)-1*H*-pyrazol-1-yl)pyridine (TAD-410530) (57)

To a stirred solution of *tert*-butyl 4-(1-(5-((2-chlorobenzyl)oxy)pyridin-3-yl)-1*H*-pyrazol-4-yl)piperidine-1-carboxylate (150 mg, 320 µmol) in dichloromethane (2 mL, 31.4 µmol) was added 4Nhydrochloric acid in dioxane (0.5 mL, 2mmol) at room temperature. After 2 hours, additional 4 N hydrochloric acid in dioxane (0.5 ml, 2 mmol) was added and the reaction mixture was stirred overnight. After completion (monitored by LCMS) the solution was concentrated in vacuo and washed with methyl tert-butyl ether (2 x 2 mL). The remaining solids were dried *in vacuo* to afford the HCl salt of 3-((2-chlorobenzyl)oxy)-5-(4-(piperidin-4-yl)-1*H*-pyrazol-1-yl)pyridine (118 mg, 267 µmol) as a white solid. ^1^H NMR (400 MHz, DMSO-*d*_6_) δ 9.22-9.12 (m, 1H), 9.05-8.93 (m, 1H), 8.82 (s, 1H), 8.60 (s, 1H), 8.41 (s, 1H), 8.11 (s, 1H), 7.80 (s, 1H), 7.71-7.66 (m, 1H), 7.60-7.54 (m, 1H), 7.49-7.40 (m, 2H), 5.38 (s, 2H), 3.30 (d, 2H, *J* = 12 Hz), 3.00 (q, 2H, *J* = 11.2 Hz,), 2.93-2.84 (m, 1H), 2.12 (d, 2H, *J* = 13.4 Hz\), 1.79 (q, 2H, *J* = 13.2 Hz), LRMS (ES+) m/z 369 [M+H]^+^; LC-MS purity 100.0% (Method C). Calculated [M+H]^+^ for C_20_H_21_ClN_4_O: 369.1482; Found: 369.1484.

### *tert*-Butyl 4-(1-(5-((2-chlorobenzyl)oxy)pyridin-3-yl)-1*H*-pyrazol-3-yl)piperidine-1-carboxylate (intermediate) (TAD-0410406)

To a reaction vessel was added tert-butyl 4-(1*H*-pyrazol-3-yl)piperidine-l-carboxylate (210 mg, 836 µmol), 3-bromo-5-(2-chlorobenzyloxy)pyridine (498 mg, 1.mmol), copper(II) iodide (27 mg, 83.6 µmol), potassium carbonate (289 mg. 2.09 mmol), potassium phosphate (284 mg, 1.34 mmol), acetonitrile (9 mL, 172 µmol), and (1R,2R)-N1,N2-dimethylcyclohexane-1,2-diamine (119 mg, 836 µmol). The vessel was purged with nitrogen, sealed and heated 120 °C for 4 hours. After the reaction was complete (monitored by LCMS), the mixture was cooled to room temperature and diluted with saturated aqueous ammonium chloride (5 mL). This mixture was extracted with ethyl acetate (3 x 5 mL). The combined extracts were washed with saturated aqueous ammonium chloride (2 mL), dried over anhydrous sodium sulfate, filtered and concentrated. The residue was purified by flash chromatography using a gradient of 0% to 50% ethyl acetate in hexane as an eluent. Desired fractions were concentrated to afford tert-butyl 4-(1-(5-((2-chlorobenzyl)oxy)pyridin-3-yl)-1*H*-pyrazol-3-yl)piperidine-1-carboxylate (341 mg. 727 µmol, 87.0%) as clear colorless oil. ^1^H NMR (400 MHz, DMSO-*d*_6_) δ 8.73 (s, 1H), 8.53 (s, 1H), 8.30 (s, 1H), 7.90 (s, 1H), 7.70-7.65 (m, 1H), 7.57-7.53 (m, 1H), 7.46-7.39 (m, 2H), 6.51 (s, 1H), 5.33 (s, 2H), 4.01 (d, 2H, *J* = 10.2 Hz), 3.02-2.73 (m, 3H), 1.91 (d, 2H, *J* = 12.01.59-1.47 (m, 2H), 1.42 (s, 9H); LRMS (ES+) m/z 469 [M+H]^+^; LC-MS purity 90.97% (Method C).

### 3-((2-Chlorobenzyl)oxy)-5-(3-(piperidin-4-yl)-1*H*-pyrazol-1-yl)pyridine (58) (TAD-0410407)

To a stirred solution of tert-butyl 4-(1-(5-((2-chlorobenzyloxy)pyridin-3-yl-1*H*-pyrazol-3-yl)piperidine-1-carboxylate (280 mg, 597 µmol) in dichloromethane (3 mL, 47.1 µmol) was added 4N hydrochloric acid in dioxane (3 mL, 12 mmol) at room temperature. After 1 hr, the reaction was complete (monitored by LCMS) and white solids settled to the bottom of the vessel. The supernatant was removed, and the remaining solids were washed with dichloromethane (3 x 5 mL). The remaining solids were dried *in vacuo* to afford 3-((2-chlorobenzyl)oxy)-5-3(piperidin-4-yl-1*H*-pyrazol-1-ypyridine (259 mg, 586 µmol, 98.2%) as a white solid. ^1^H NMR (400 MHz, DMSO-*d*_6_) δ 9.26-9.17 (m, 1H), 9.05-8.95 (m, 1H), 8.80 (s, 1H), 8.65 (s, 1H), 8.43 (s, 1H), 8.11 (s, 1H), 7.72-7.66 (m, 1H), 7.59-7.54 (m, 1H), 7.48-7.40 (m, 2H), 6.54 (s, 1H), 5.38 (s, 2H), 3.31 (d, 2H, *J* = 11.5 Hz), 3.08-2.97 (m, 3H), 2.14 (d, 2H, *J* = 13.6 Hz), 1.97-1.85 (m, 2H). LRMS (ES+) m/z 369 [M+H]^+^; LC-MS purity 100.0% (Method C). Calculated [M+H]^+^ for C_20_H_21_ClN_4_O: 369.1482; Found: 369.1484.

### 3-((2-Chlorobenzyl)oxy)-5-(1-(piperidin-4-yl)-1*H*-pyrazol-3-yl)pyridine. TFA (63) (TAD-0410839)

^1^H NMR (400 MHz, DMSO-*d*_6_) δ 8.67 (d, 1H, *J* = 1.6 Hz), 8.64 (m, 1H), 8.39 (m, 1H). 8.34 (d, 1H, *J* = 2.8 Hz), 7.90 (d, 1H, *J* = 2.4 Hz), 7.83-7.82 (m, 1H), 7.68-7.65 (m, 1H), 7.56-7.53 (m, 1H), 7.45-7.40 (m, 2H), 6.92 (d, 1H, *J* = 2.4 Hz), 5.31 (s, 2H), 4.59-4.54 (m, 1H), 3.46-3.42 (m, 2H), 3.05-3.06 (m, 2H), 2.26-2.14 (m, 4H); LRMS (ES+) m/z 369.25 [M+H]^+^; LC-MS purity 98.69% (Method A). Calculated [M+H]^+^ for C_20_H_21_ClN_4_O: 369.1482; Found: 369.1485.

### Enzymatic inhibition

The SHIP1 and SHIP1 proteins were prepared and characterized as previously described^7^. Briefly, the 2D human SHIP1^395-898^ Ptase-C2 domain protein and 5D human SHIP1^1-899^ and mouse SHIP1^1-861^ multidomain protein constructs were expressed and purified from baculovirus P2 viral stocks. The 2D human SHIP2^420-878^ Ptase-C2 domain was expressed and purified from *E.coli*. The enzymes were preincubated with compounds in assay buffer (50 mM HEPES, 150 mM NaCl, 2 mM MgCl_2_) for 20 min before adding substrate. For the human 2D proteins, the final reaction concentration of the PI(3,4,5)P3-diC8 substrate (Cayman Chemical, 10007764) was 52 µM, while the concentrations of the enzymes were 10 nM for SHIP1 and 50 nM for SHIP2. Reactions were quenched after 10 min with Malachite BioMol Green (Enzo, BML-AK111-1000) and then incubated for 30 min at room temperature and absorbance (620 nm) measured. For the human and mouse 5D proteins, the final reaction concentrations for the substrate and the enzymes were 40 µM and 10 nM correspondingly. Reactions were quenched after 2 min and then incubated for 30 min at room temperature and absorbance measured. Inhibitory potency (IC_50_) values were calculated by fitting absorbance versus inhibitor concentration.

### Bioluminescence-based Thermal Shift Assay (BiTSA)

We employed a HiBit tag on full length SHIP1 and established a stable clone of HMC3 cells expressing HiBit-*INPP5D*, similar to the reported BiTSA^51^. An expression construct encoding full-length human SHIP1 encoded by *INPP5D*, tagged at the N-terminus with the HiBiT peptide (11 amino acid sequence from Promega), was generated by subcloning the open reading frame into the pcDNA3.1(+) expression plasmid under the control of CMV promoter and NeoR NDA for selection. The resulting plasmid was sequence-verified and used to transfect human microglial HMC3 cells via Fugene 6 (Promega). Following antibiotic selection with 400 µg/ml Geneticin (Gibco) and serial cell dilution in 24 well plates, individual clones resistant to the selection were picked up for subculture to generate enough cells screened for stable integration and expression of the HiBiT-tagged SHIP1 protein suitable for use with luminescence-based detection with the Nano-Glo® HiBiT Lytic Detection System (Promega). A clonal population exhibiting robust and consistent expression of HiBiT-*INPP5D* was isolated and expanded in the DMEM GlutaMax media with 10% FBS, 1X Penicillin/ Streptomycin and 300 µg/ml Geneticin for passage the cell culture while keeping the moderate selection pressure to maintain the stable HiBit-*INPP5D* expression, frozen cell stocks were generated and stored in liquid nitrogen.

In screening mode, 10,000 cells in 20 µL/well were plated into white 96 well PCR plates (AB-3396/B, Thermo Scientific), then treated with a single concentration of compound (100 μM) or fragments (200 μM) at 37 °C for 60 min and then exposed to 3-min isothermal heating at 44.2 °C the experimentally determined Tm of HiBit-SHIP1 (n=28), and recovering to 25 °C for 3 min in a PCR machine (Eppendorf Mastercycler X50a). The cell plate was processed with the one step Nano-Glo^®^ HiBiT Lytic Detection System Protocol (Promega) and scanned with Synergy H1 plate reader (BioteK) for luminescence. The single point screening was routinely run with 3 repeats for each compound in an assay. The raw luminescence/well is normalized to the mean luminescence/well of untreated controls (typically 8 wells/plate) from the same plate and expressed as % control. The average % control value of 3 repeats were calculated and rank ordered. The mean and SD of untreated control wells were also calculated. The active compound was determined with its % control value difference larger than mean ± 3xSD of the untreated controls. When the difference is higher than the control values,the compound stabilizes SHIP1, whereas differences lower than control values indicate SHIP1 destabilization. In dose-response mode, cells were treated with decreasing concentrations from 100 µM (compounds) or 200 μM (fragments) with 1:3 serial dilutions to generate 8-point dose-dependent data from each column, and duplicates were run in each assay. Raw luminescence/well in each plate were normalized with the average value of untreated control wells in the same plate and expressed as % control. The % control values were used to determine AC_50_ calculated using a four-parameter logistic curve regression model. Percent luminescence remaining at the highest concentration is also reported for the compounds when the difference from the DMSO control is greater than mean ± 3xSD. Otherwise, an AC_50_ was not calculated.

### pAKT/tAKT assay

THP-1 cells were cultured in THP-1 medium, RPMI 1640 (Wisent #350-007 CL) with 10% FBS (Wisent, #090-150) + 1 % Penicillin/Streptomycin (Wisent, # 450-201), at a density of 0.8-1.0 × 10⁶ cells/mL, collected, resuspended, counted and adjusted to 0.5 × 10⁶ cells/mL in a final volume of 30 mL, centrifuged at 200 xg for 5 minutes, and resuspended in 30 mL of fresh THP-1 medium. Cells were stimulated with 20 ng/mL recombinant human IL-4 (PeproTech® ThermoFisher, #200-04-50UG, 0.1 mg/mL stock; 6 μL per 30 mL culture) and incubated at 37 °C, 5% CO₂ in T175 cm^2^ flask for 24 hours. Following stimulation, cells were harvested into 50 mL conical tubes, counted for density and viability, and pelleted by centrifugation at 200 xg for 5 minutes. Supernatants were discarded, and cells were resuspended in appropriate volume to meet 0.5 × 10⁶ cells/mL in fresh THP-1 medium without phenol red and biotin and supplemented with 10% FBS. Cultures were returned to T175 cm^2^ flask and incubated for an additional 24 hours under the same conditions to allow recovery. THP-1 cells were then seeded at 5,000 cells/well in AlphaPlate 384 shallow-well plates (ProxiPlate; Revvity #6008350) and equilibrated for 2 hours at 37 °C, 5% CO₂ in RPMI without biotin or phenol red. Cells were then treated with compound in 2-fold serial dilutions starting at 100 μM. Each condition was tested in duplicate (n = 2), and treatments were carried out for 15 minutes at 37 °C, 5% CO₂. Phosphorylated and total AKT (pAKT/tAKT) levels were quantified using the Revity Alpha SureFire Ultra Multiplex PhosphoAKT (S473 or T308) kit (MPSU-PTAKT) according to the manufacturer’s instructions for the 1-plate assay protocol. Plates were incubated at 22 °C with both Acceptor and Donor Mix reagents before reading. Alpha signal detection was performed using a Tecan Spark plate reader using the AlphaPlex protocol (Excitation: 680 nm; Emission: 615 nm for pAKT1/2/3 and 545 nm for tAKT). The pAKT(S473)/tAKT or pAKT(T308)/tAKT ratio was calculated for each well, and data were normalized to % inhibition based on DMSO-treated and maximal control wells. IC_50_ values were determined using a four-parameter variable slope curve fit in GraphPad Prism.

### Lipidomic profiling

Phosphoinositides were measured using targeted liquid chromatography tandem mass spectrometry. THP-1 pellets were lyophilized overnight before extraction. Lipids were extracted in Eppendorf tubes using a single-phase extraction method with 2:2:1 butanol:methanol:water with 1 mM of ammonium formate and 50 µM of medronic acid. Into each sample, 40 pmol of phosphatidylinositol phosphate internal standards were also added PI(3’,5’)P2 16:0/16:0-d5 (Avanti Research) and PI(5’)P 16:0/16:0-d5 (Avanti Research). Samples were vortexed and sonicated in a sonicator bath for 1 hour before centrifugation at 13,000 x g for 10 minutes. The supernatant was transferred into glass vials for mass spectrometry analysis.

Phosphatidylinositol phosphates were measured using a BEH Premier C8 VanGuard-FIT column (2.1 x 100mm, 1.7µm, Waters) operated at 55 °C. Solvents comprise of A) 1:1 water:acetonitrile with 20mM ammonium bicarbonate and 5 µM medronic acid and B) 1:1:8 water:acetonitrile:isopropanol with 10mM ammonium bicarbonate and 2.5 µM medronic acid. Targeted analysis was conducted using an Agilent 6495C. Mass spectrometer conditions for phosphatidylinositol phosphates were as follows: gas temperature, 250 °C; gas flow rate, 15 L/min; nebulizer, 20 psi; sheath gas temperature, 320 °C; capillary voltage, 4500 V and sheath gas flow, 12 L/min. Both phosphatidylinositol monophosphate and diphosphate classes were measured in positive mode as [M+H]+ ions with the neutral loss of the phospho-inositol and the additional phosphates as the product ions. Relative concentrations were calculated by normalizing the species against the corresponding internal standards, PI[5’]P 16:0/16:0-d5 and PI[3’,5’]P 16:0/16:0-d5 (Avanti Research).

### Protein Lipid Overlay Assay

Recombinant SHIP1 5-domain protein was expressed and purified as previously described. For assays containing compound 32, the molecule was present throughout all incubation and washing steps after the blocking step at stated concentrations. Commercially available PIP arrays (Echelon Biosciences, P-6100) were incubated with blocking solution (PBS + 5% BSA) overnight at 4 °C. The solution was decanted and recombinant SHIP1 5D protein (10 nM, PBST + 3% BSA) was incubated for 1 hour at room temperature. The membrane was washed 3x with PBST + 3% BSA followed by addition of primary antibody (Invitrogen SHIP1 Monoclonal Antibody, SHIP-02, Thermo Fisher #MA1-10450) in PBST + 3% BSA. This was allowed to incubate at room temperature for 1 hour. The primary antibody solution was discarded, and membrane was washed 3x for 5 minutes with PBST + 3% BSA. Secondary antibody was then allowed to incubate with the membrane for 1 hour followed by two 5-minute washes with PBST + 3% BSA followed by a final five-minute wash with PBS. The membrane was then imaged using a Licor Odessey CLx.

### Primary microglial culture

Microglia were isolated as described previously^60-62^. Briefly, cortical and hippocampal tissue from the neonatal C57BL/6J (*Inpp5d^WT^*; The Jackson Laboratory, JAX #:000664) mice (P0-P3), B6.129S6(C)-*Inpp5d*^tm1.1Wgk^/J (*Inpp5d^HET^*; The Jackson Laboratory, JAX #:028269) mice (P0-P3) or *Inpp5d*^HOM^ (bred in house) mice (P0-P3) were homogenized in Dulbecco’s Modified Eagle Medium (DMEM; Gibco, 10566016), sequentially filtered through 250 and 100 μm meshes, and cultured in the Advanced DMEM/F12 media (Gibco, 12634028) supplemented with 10% fetal bovine serum (Gibco, 16000044), 1x GlutaMax (Gibco, 35050061), and 1x Penicillin/Streptomycin (Gibco, 15140122). At 21 days *in vitro* (DIV), the cultures were subjected to mild trypsinization using 0.0625% Trypsin-EDTA (Gibco, 25200072) in DMEM for 30 minutes to detach an intact layer of astrocytes. The microglia attached to the bottom was used for the amyloid uptake and treatment studies.

### High-content imaging assay

The microglial phagocytosis/cell health high-content assay has been described in detail elswhere^63^. In brief, BV2, HMC3, or primary microglia plated in 384 well cell imaging plate (Falcon 353962, Corning) at 400 cells/45ul/well for BV2, 600 cells/45ul/well for HMC3 and 2000 cells cells/45ul/well for primary microglia. Cells were treated with compounds by adding 5 ul/well of 10x of final concentration compound for 24 hrs and then seeded with adding 5ul/well of 50 ug/ml pHrodo-myelin/membrane debris. After 20 hrs, nuclear staining solution (10ug/ml Hoechst 33342 in media) was added to cell plates with 20ul/well, and the plates were incubated for another 30 min and then scanned with an ArrayScan XTI high-content analysis reader (Thermo Scientific), and the associated image data was analyzed with Thermo Scientific HCS Studio. Since the assay is mix-and-read without liquid change, the imaging cell count is reliable. Mean total phagocytosis spot intensity per cell, total cell counts per well and mean average nuclear intensity per cell for cell health were measured. Cellular potencies for each endpoint (EC_50_ for phagocytosis, IC_50_ for cell count) were calculated using a four-parameter logistic curve regression model.

### Amyloid uptake assay

Primary microglial cultures were incubated with HiLyte Fluor 488-labeled fAβ₁₋₄₂ (aggregated at 37 °C for 5 days; 0.5 μM; AnaSpec, AS-65627) in serum-free Advanced DMEM/F12 medium for 30 minutes, unless otherwise specified. Cultured microglia were then fixed with 4% paraformaldehyde overnight. Next day, cells were washed, and permeabilized with PBST. Slides were blocked with 5% normal donkey serum and sodium azide (0.03%) for 1 hour at RT. An anti-Iba1 antibody (goat, 1:1000; Novus Biologicals, NB100-1028) diluted in blocking solution, and cells were incubated overnight at 4°C. After washing with PBST, cells were incubated with an Alexa Fluor 546-conjugated anti-goat secondary antibody (1:1000; Invitrogen, A11056) in blocking solution for 1 hour at RT. Nuclei were stained with 4′,6-diamidino-2-phenylindole (DAPI). Images were acquired using a fluorescence microscope, and image analysis was performed with ImageJ2 (NIH, version 1.53t). For inhibitor studies, microglial cultures were pre-treated with vehicle (0.1% DMSO) or Compounds 22, 24, or 28 at concentrations of 0.3, 1, or 3 µM for 6 hours, followed by a 30-minute fAβ exposure. Four biological replicates were performed per condition. Four biological replicates were performed for each condition. For each replicate, eight fields of view were acquired, and the resulting images were quantitatively analyzed. Each data point therefore represents the mean value derived from these eight independently scanned and analyzed images within a single biological replicate.

### Lactate dehydrogenase (LDH) cytotoxicity assay

LDH released into culture supernatant was used as a measure of cytotoxicity^81^. The LDH Cytotoxicity Assay Kit (Abcam, ab65393) was used according to the manufacturer’s instructions. In brief, 10 µL of culture supernatant was combined with 100 µL of LDH reaction mix and incubated at 37 °C for 30 minutes. Absorbance was measured at 450 nm and 650 nm using a microplate reader.

### Animal Studies

All animal studies conform to the Guide for the Care and Use of Laboratory Animals published by the National Institutes of Health (NIH Publication No. 85-23, revised 2011) and were reviewed and approved by the respective Institutional Animal Care and Use Committees (IACUC) at The University of Pittsburgh (PITT) and Indiana University (IU) prior to study initiation.

### Pharmacokinetics Study

Adult male wildtype mice (C57BL/6J JAX#:000664) from The Jackson Laboratory were received at approximately 8 weeks of age and acclimated for at least 1 week to the PITT animal facility prior to the experiments. Mice were group housed (n=3-4 per cage) with *ad libitum* access to food and water. The housing room consists of ventilated caging and automated watering system with room temperature controlled at a setting of 72±2 °F and humidity at 50±10%. The testing facility was on a 12:12 L:D schedule (lights on at 7:00 AM). The study was conducted in non-fasted subjects with compounds administered at the beginning of the light cycle. Mice (n=3) were dosed with compound **32** (100 mg/kg) via oral gavage (20 ml/kg formulated at 5 mg/mL dose volume) in a vehicle solution of 1% hydroxypropyl methylcellulose (HPMC)/Tween 80 (0.25%)/purified water^82, 83^. Serial blood samples (50 µL) were collected via the tail sampling procedure in non-anesthetized mice into heparinized capillary tubes. Blood was transferred into Eppendorf tubes maintained on wet ice and plasma was processed within 20-60 min post blood collection by centrifugation at 14500 rpm for 10 minutes at 4 °C. Plasma aliquots were frozen on dry ice and stored at -80 °C until analysis. For terminal tissue collection, mice were subjected to isoflurane anesthesia to the surgical plan of anesthesia followed by decapitation and brain removal. Brains were briefly rinsed in cold PBS and snap frozen on dry ice followed by storage at - 80 °C until processing and analysis. The left hemisphere without cerebellum was analyzed for brain exposure levels of compound **32** at the 24-hour terminal timepoint.

### Bioanalytical procedures

The method to quantify compound **32** from mouse plasma and brain was developed using temazepam as the internal standard, liquid-liquid extraction, and HPLC-MS/MS QTRAP (Sciex 6500+ QTRA) with an electrospray ionization probe run in positive mode. Liquid-liquid extraction of plasma (20µL) was used with the addition phosphate buffered saline (pH 7.4, 50 µL) and methyl tert-butyl ether (2mL). Phosphate buffered saline (pH 7.4) was added to brain tissue to a total volume of 0.5 mL prior to homogenization using Benchmark Bead Blaster. Following addition of internal standard, 3 mL methyl tert-butyl ether was added to 0.5 mL homogenate. The mobile phase was run as gradient using 0.1% formic acid in water (Solvent A) and acetonitrile (Solvent B) on a Phenomenex Onyx Monolithic 100X4.6 mm column with a flow rate of 500 µL/min. The gradient elution began with 80% A and 20% B transitioning to 5% A and 95% B at 5 minutes, which was held until 7 minutes. Compound **32** elutes at 4.51 min. The multiple reaction monitoring (MRM) Q1/Q3 (m/z) transitions for compound **32** were 369.2/125.0 and for temazepam 301.0/255.1. The lower limit of quantification of compound **32** was 1.0 ng/mL in plasma and 0.4 ng/g in brain.

### Pharmacokinetic Analysis

A naïve pooled approach was used to calculate pharmacokinetic parameters of Compound **32** in C57BL/6J mice. Mean plasma concentrations (n=3-6 mice per timepoint) were calculated and noncompartmental analysis performed in PKanalix (v2024R1, Simulations Plus). Area under the concentration-time curve (AUC) was calculated using the linear trapezoidal rule with linear interpolation following extravascular dosing.

### Pharmacodynamic Study

Male and female wild type (C57BL/6J JAX #:000664) and hAbeta^SAA^ knock-in mice (APP-SAA KI JAX #:034711) from The Jackson Laboratory were aged to 14 months of age while being housed and maintained in a 12/12-h light/dark cycle with food (Purina Lab Diet, 5K52) and water *ad libitum* in the IU School of Medicine animal facility prior to the experiments. Compound **32** was prepared in a vehicle solution consisting of 1% hydroxypropyl methylcellulose (HPMC; Sigma, H7509), 0.25% Tween 80 (Sigma, P1754), and injectable water (Millipore, 4.86505.0500). Compound **32** (150 mg/kg; 10 mL/kg body weight) was administered via oral gavage three times at 12-hour intervals. Mice were briefly restrained manually, and a sterile, disposable, flexible gavage needle (18-gauge, 1.5-inch, 2 mm round tip; BG 9928, Braintree Scientific, Inc.) was gently inserted into the esophagus and advanced into the stomach. The volume was injected in one single step.

One hour after the last administration, the animals were anesthetized, and a cardiac puncture was performed to collect blood using heparin tubes (Microvette® 500 Lithium Heparin LH, 20.1345.100). The animals were euthanized and transcardially perfused with cold PBS. Brains were rapidly extracted and hemisected. Blood plasma and one hemisphere were snap-frozen, followed by storage at -80 °C, for compound quantification in the Clinical Pharmacology Analytical Core (CPAC). The other hemisphere was also snap-frozen and stored at -80 °C until it was processed for RNA and protein isolation.

### Tissue processing, RNA Isolation and Protein extraction

Cerebellum from the hemibrains was dissected and banked. The rest of the tissue was homogenized in nine hundred microliters of tissue protein extraction reagent (T-PER) (Thermo Fisher Scientific, 78510) containing protease and phosphatase inhibitor cocktail (Thermo Fisher Scientific, 78440). One aliquot of three hundred microliters was sampled from the tissue homogenates and mixed in an equal volume of RNA STAT-60 (Tel-Test Inc., CS-502). RNA was isolated using the PureLink RNA mini kit (Invitrogen, 12183020) and PureLink DNase (Invitrogen, 12185010) according to the manufacturer’s protocol. RNA quality and quantity were determined using the Nanodrop 2000 spectrophotometer (Thermo Fisher Scientific) and stored at -80 °C until processing and analysis. The rest of the homogenate was centrifuged at 1000 g x 10 min at 4°C. The supernatant was stored at -80 °C for the quantitative cytokine assay.

### Nanostring nCounter assay

Two hundred nanograms of brain RNA from 14-month-old B6 and control or treated SAA mice were used for gene expression profiling with the nCounter mouse Glial and Neuroinflammation Profiling panels (NanoString Technologies) following the manufacturer’s directions.

Nanostring nCounter data from two panels, Glia panel (Glia) and Inflammation panel (Infl), were used as input to our differential gene expression (DGE) pipeline. Separate DGE analyses were performed for each panel independently to simplify the workflow and make the results more interpretable. For each panel (*i.e.*, Glia and Inflammation) we converted the RCC files from the Nanostring nCounter platform to a human readable CSV file using the nanostring package. The list variable was then converted to a matrix and saved as a CSV file. These CSVs were loaded into a NanoStringSet object then normalized using the estimated normalization factors from the NanoStringDiff package. Based the normalized NanoStingSet object, a negative binomial generalized linear model was fit to perform the final DGE estimates. Differentially expressed genes (DEGs) were identified based on the contrasts between SAA, B6, and SAA treated with compound **32** groups. In total six different sets of DEGs were identified: Glia SAA vs. B6, Glia SAA+cmpd32 vs. B6, Glia SAA+ cmpd32 vs. SAA, Infl SAA vs. B6, Infl SAA+cmpd32 vs. B6, and Infl SAA+cmpd32 vs. SAA. These DEG sets were visualized using heatmaps (pheatmap), and volcano plots (EnhanncedVolcano).

### Meso Scale Discovery (MSD)-based quantitative cytokine assay

The MSD quantitative cytokine assay was performed as described previously^60^. Tissue homogenates were centrifuged at 800 x g 10 min at 4 °C. The supernatant was utilized for analysis. Samples were assayed in duplicate using the MSD V-PLEX mouse proinflammatory Panel I (Meso Scale Diagnostics, K15048D) assay.

### Statistical analysis

Data are presented as mean ± SEM. Statistical analyses were performed using unpaired t-tests, one-way ANOVA, or two-way ANOVA, as appropriate. Data were assessed for normality, and non-parametric tests were applied when assumptions of normality were not met. Significant main effects or interactions were followed by post hoc multiple comparisons using Tukey’s test for parametric data or Dunn’s test post hoc for non-parametric data. A p-value < 0.05 was considered statistically significant. All analyses were conducted using GraphPad Prism (version 9.3.1; GraphPad Software).

## SUPPORTING INFORMATION

Supplementary materials containing Figures S1-S7 and Table S1. The mass spectrometry proteomics data "Kinome profiling of BV2 cells treated with crizotinib or compound 32" have been deposited to the ProteomeXchange Consortium via the PRIDE^84^ partner repository with the dataset identifier PXD071931.

## CONTRIBUTIONS

C.D.J. and C.R.-B. contributed equally to this work. P.A.H., C.D.J., D.E.B., and W.B.C. designed molecules, oversaw and performed chemical syntheses, and analyzed data. A.D.M., A.K.H., K.S., and R.D.I. oversaw biochemical studies, prepared and characterized proteins, performed *in vitro* studies, and analyzed data. S.C., J.L.D., A.L.O., C.R.-B., J.J.-W., E.R.M., and M.M. oversaw, designed and performed *in vitro* cellular assays and analyzed data. S.P.A., C.J.B. and D.M.S. oversaw and performed cellular kinome profiling studies, and analyzed data. R.K.-D., P.J.M., A.N.F., M.v.B.-M., J.C.D. and K.H. oversaw, designed, and performed lipidomic studies, and analyzed data. S.J.S.R., S-P.W, S.D. and H.H. oversaw, designed, and conducted *in vivo* pharmacokinetic studies. L.D.S. and S.K.Q. oversaw bioanalytical studies and analyzed data. A.L.O., I.H.C.-F., C.R.-B. oversaw, designed and performed *in vivo* pharmacodynamic studies, and analyzed data. T.S.J. analyzed RNA transcriptional data and performed statistical analyses. B.T.L. and A.D.P. conceived the project and secured funding. T.I.R. secured funding, supervised the project, designed studies, analyzed data, and wrote the manuscript with input from all authors.

## ACKNOWLEDGMENTS

We thank our advisors at the Harrington Discovery Institute Therapeutic Development Center for their expert guidance and helpful suggestions especially Peter Bernstein. The logistical and project management support of Ariel Bontrager at Purdue University, Becky Klein at the Indiana Biosciences Research Institute, and Ankita Ankita Sharma at Dalriada are gratefully acknowledged. We thank Jerry Di Salvo at Evotec for overseeing, and Sarah Souza for performing, enzyme assays. We thank Jens Tiefenbach and his team at Dalriada for running cellular assays. We thank Avi Benitah and Cidney Hart for running western blot analyses. The authors are grateful for the resources and support of the University of Pittsburgh Preclinical Phenotyping Core (PPC). Mass spectrometry was provided by the Clinical Pharmacology Analytical Core (CPAC) at Indiana University School of Medicine. All intellectual property rights, including copyrights and trademarks, relating to CETSA^®^ are owned by Pelago Bioscience AB. BioRender was used to design figures.

## FUNDING

This work was primarily supported by NIH National Institute on Aging (NIA) grants and supplements that are components of the Accelerating Medicines Partnership for AD (AMP-AD), the TaRget Enablement to Accelerate Therapy Development for Alzheimer’s Disease (TREAT-AD) Centers, and the Alzheimer’s Drug Development Program: U54AG065181 (Palkowitz, Lamb, Richardson); U01AG088021 (Richardson, Palkowitz, Mesecar, Lamb, Oblak); R01AG081322 (Kaddurah-Daouk, Arnold, Meikle); RDI was supported by T32AG071444 (Landreth); CB was supported by CA272370 (Kelley, Clapp). SPA acknowledges support from the Indiana University Precision Health Initiative. Mass spectrometry for bioanalytical was provided by the Clinical Pharmacology Analytical Core at Indiana University School of Medicine; a core facility supported by the IU Simon Cancer Center Support Grant P30 CA082709. KH and PJM are supported by National Health and Medical Research Council (NHMRC) investigator grants (1197190 and 2009965). TIR acknowledges support from Alzheimer’s Drug Discovery Foundation (RHDl-202305-2025123).

## CONFLICTS OF INTEREST

All funding provided to the institution and individual authors has been disclosed in the funding sources. CDJ, DEB, WBC, and TIR have patents pending or planned related to this work. JLD, BTL, ADP and TIR are founders and consultants for Monument Biosciences and own stock. CDJ, DEB, WBC, PAH, SJSR and ADM own stock in Monument Biosciences. JLD is an inventor on patents or patent applications assigned to Eli Lilly and Company relating to the assays, methods, reagents and / or compositions of matter for P-tau assays and Aβ targeting therapeutics. JLD has/is served/serving as a consultant or on advisory boards for Eisai, Abbvie, Genotix Biotechnologies Inc, Gates Ventures, Syndeio Biosciences, Dolby Family Ventures, Karuna Therapeutics, Alzheimer’s Disease Drug Discovery Foundation, AlzPath Inc., Cognito Therapeutics, Inc., Eli Lilly and Company, Prevail Therapeutics, Neurogen Biomarking, Spear Bio, Rush University, University of Kentucky, Tymora Analytical Operations, MindImmune Therapeutics, Inc, Early is Good, and Quanterix. JLD has received research support from ADx Neurosciences, Fujirebio, Roche Diagnostics and Eli Lilly and Company in the past two years. JLD has received speaker fees from Eli Lilly and Company and LabCorp. JLD is a founder of Dage Scientific LLC. JLD has stock or stock options in Eli Lilly and Company, Genotix Biotechnologies, MindImmune Therapeutics Inc., AlzPath Inc., and Neurogen Biomarking. BTL receives licensing fees from Ionis Pharmaceuticals, participates on the Advisory Board and receives consulting fees from NervGen Inc. and The Cleveland Clinic, has a leadership or fiduciary role at the Alzheimer’s Association and Cure Alzheimer’s Fund, has received travel support from Alzheimer’s Association and Cure Alzheimer’s Fund, and Department of Defense. ADP is president and CEO of the Indiana Biosciences Research Institute (IBRI). SJSR has received travel support from the Alzheimer’s Association and is a paid consultant of Hager Biosciences. TIR is a consultant for Candenza and has stock options. No other authors have conflict of interests to disclose.

