## Supplementary Materials for "Optimization and Characterization of SHIP1 Ligands for Cellular Target Engagement and Activity in Alzheimer’s Disease Models"

1. Lgenia, Fortville, Indiana 46040, United States; 2. Indiana University School of Medicine, Indianapolis, Indiana 46202, United States; 3. Stark Neurosciences Research Institute, Indiana University School of Medicine, Indianapolis, Indiana 46202, United States; 4. Purdue University, West Lafayette, Indiana 47907, United States; 5. Baker Heart and Diabetes Institute, Melbourne, Victoria 3004, Australia; 6. University of Pittsburgh School of Medicine, Pittsburgh, Pennsylvania 15213, United States; 7. La Trobe University, Melbourne, Victoria 3086, Australia; 8. Indiana Biosciences Research Institute, Indianapolis, Indiana 46202, United States; 9. Duke Institute for Brain Sciences, Duke University, Durham, North Carolina 27708, United States

**TABLE OF CONTENTS**

Figure S1. SHIP1 expression in THP-1 cells ........ S2

Figure S2. Effect of compound **32** on phosphatidylinositol (PI) species ........ S2

Figure S3. SHIP1 expression in BV2 cells and primary murine microglia ........ S3

Figure S4. High content imaging myelin/membrane debris uptake and cell health ........ S4-5

Figure S5. Time course of compound **32** enhancement of microglial uptake of $A\beta_{1-42}$ fibrils ........ S6

Figure S6. Cellular kinome profiling ........ S7

Figure S7. Meso Scale Discovery (MSD) Mouse Proinflammatory Panel ........ S8

Table S1. Phosphate 5-kinase (PIP5K) family activity ........ S9

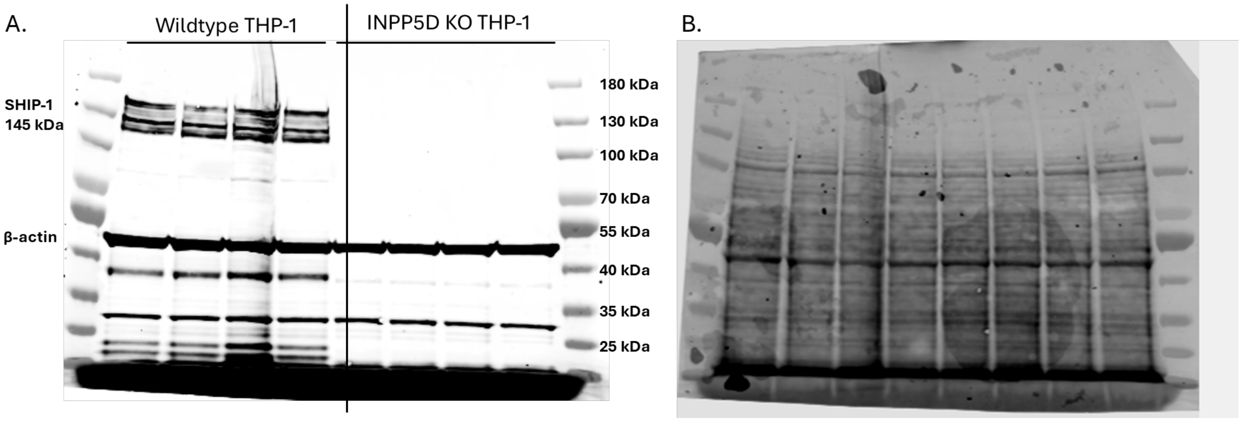


**Figure S1 | SHIP1 protein expression in THP-1 cells.** Multiple bands may reflect distinct INPP5D/SHIP1 isoforms or truncated species from alternative splicing, proteolytic cleavage, or post-translational modifications as RefSeq and Ensembl annotate numerous INPP5D splice variants, and UniProt lists SHIP1 isoforms generated by alternative splicing. Wild type THP-1 cells (ATCC TIB-202) were maintained in RPMI-1640 medium supplemented with 10% fetal bovine serum and 0.05 mM 2-mercaptoethanol. The THP1 INPP5D (SHIP-1) knockout line (YKO-HT0445, Ubigen Biosciences, Guangzhou, China) was generated using CRISPR gene editing. Lysates were collected in RIPA lysis buffer (Thermo Fisher Scientific 89901) supplemented with protease and phosphatase inhibitors (Thermo Fisher Scientific 1861281). **A.** Western blot probed with anti-SHIP1 (1:1000, Thermo Fisher Scientific MA1-10450) and anti-β-actin (1:1000, Cell Signaling Technology 13E5). Immunoreactivity visualized using the Odyssey CLx system (LI-COR). **B.** Ponceau S stain of the same blot demonstrating total protein loading.

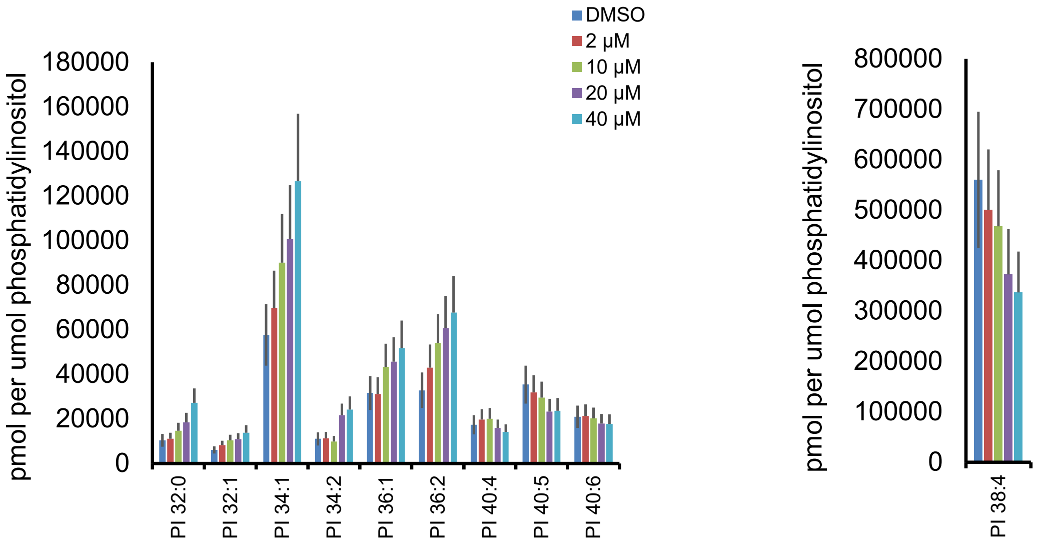


**Figure S2 | Effect of compound 32 on the phosphatidylinositol (PI) pool.** Shorter chained PIs, particularly PI 34:1, were increased while the longer chain and more abundant species PI 38:4 was reduced.

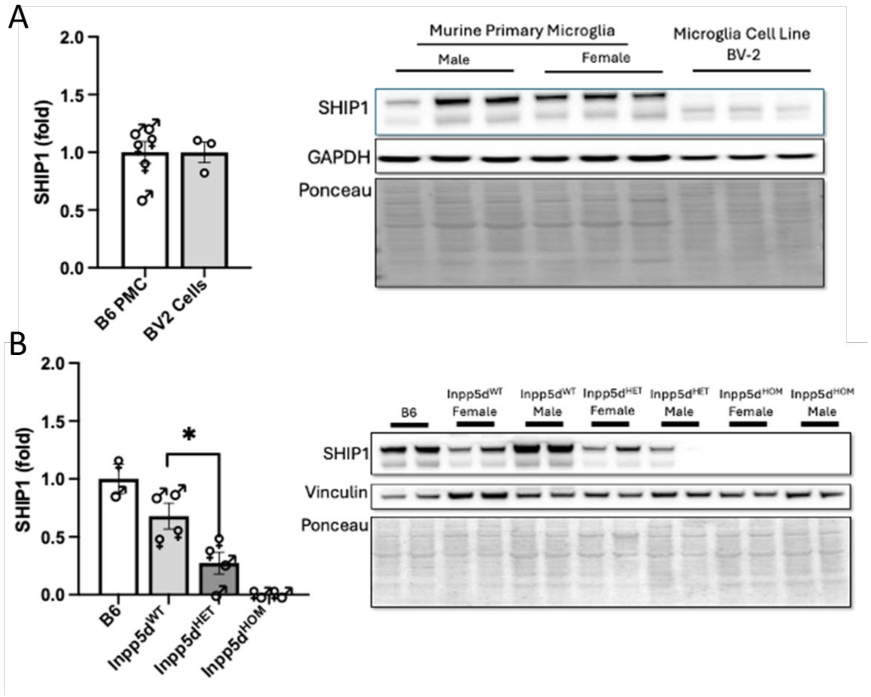


**Figure S3 | SHIP1 expression in BV2 cells and primary murine microglia.** Immunoblot analysis of SHIP1 protein levels using the Thermo Fisher Scientific MA1-10450 antibody shows multiple bands in mouse microglia and BV2 cells, analogous to those observed in THP-1 cells (**Figure S1**), which may reflect SHIP1 isoforms or truncated species. **(A)** Comparative analysis of SHIP1 levels in B6-derived primary microglia (B6 PMC) versus immortalized BV2 cells. Both bands quantified together and normalized to GAPDH. Data are presented as mean ± SEM. **(B)** SHIP1 expression in primary microglia isolated from female (♀) and male (♂) mice of the following genotypes: B6, *Inpp5d*$^{WT}$, *Inpp5d*$^{HET}$, and *Inpp5d*$^{HOM}$. Both bands quantified together and normalized to Vinculin. A significant reduction in SHIP1 expression is observed in *Inpp5d*$^{HET}$ microglia compared to controls. Statistical significance was assessed using Kruskal–Wallis by Dunn's multiple comparisons test or Mann–Whitney tests, as appropriate (*$p < 0.05$).

Primary microglia lysates from B6 WT, Inpp5d$^{HET}$ and Inpp5d$^{HOM}$ mice were collected using the Mammalian Protein Extraction Reagent (M-PER; Thermo Fisher Scientific, 78501) with Protease and Phosphatase Inhibitor Cocktail (Thermo Fisher Scientific, 78447). The protein concentrations were then measured with a bicinchoninic acid (BCA) protein assay kit (Thermo Fisher Scientific, 23225) from tissue lysates. Western blot analysis was performed using the following antibodies: anti-Ship1 (1:1000, Thermo Fisher MA1-10450), anti-GAPDH (1:1000; Abcam, Ab70699), anti-Vinculin (1:1000, Millipore Sigma, V9131). The immunoreactivity was visualized using the iBright® Imagining System (Thermo Fisher Scientific). Densitometry analysis was performed using ImageJ2 (NIH, version 1.53t).

**A**

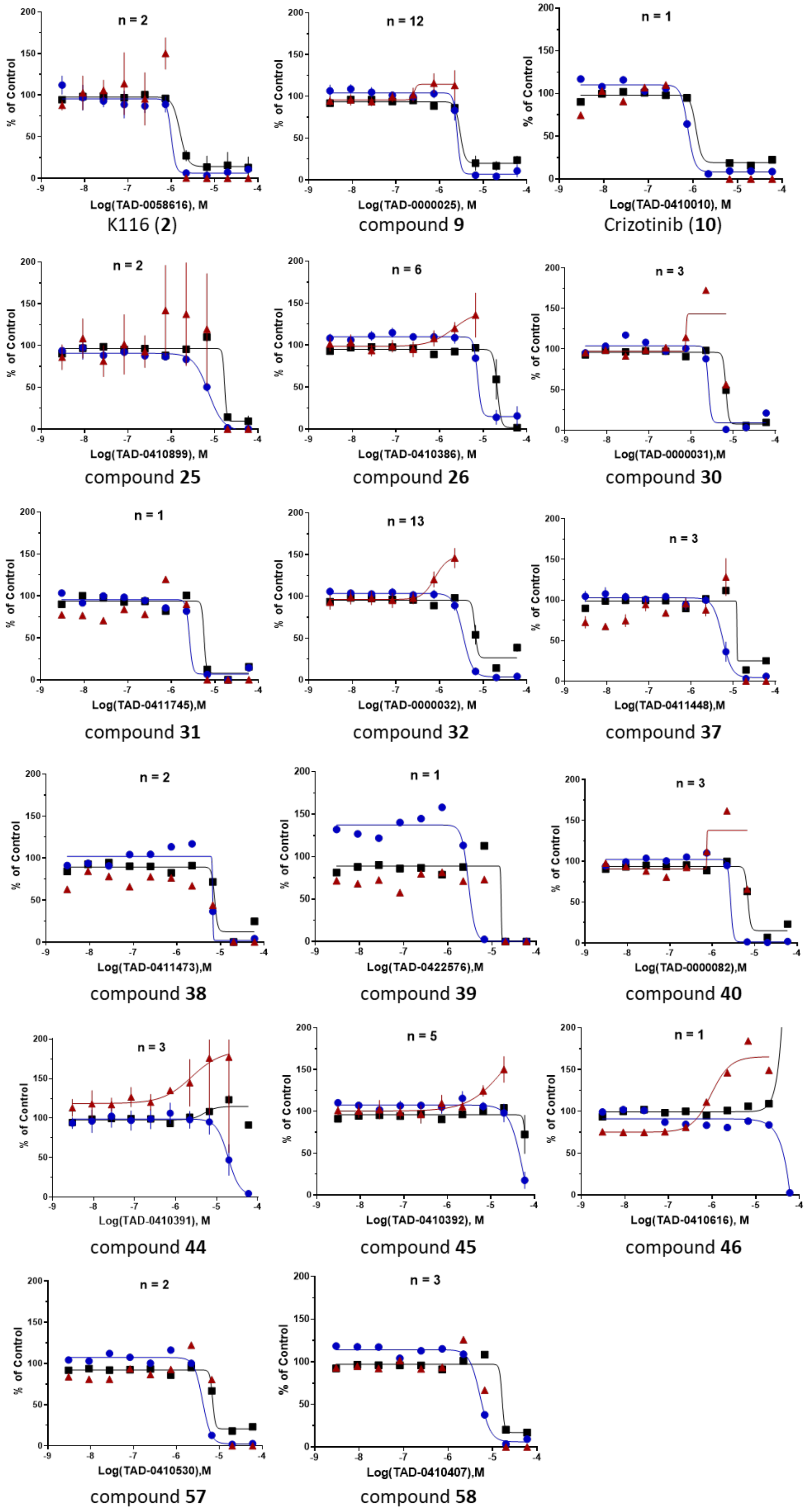

n = 2
% of Control
Log(TAD-0058616), M
K116 (2)
n = 12
% of Control
Log(TAD-0000025), M
compound 9
n = 1
% of Control
Log(TAD-0410010), M
Crizotinib (10)
n = 2
% of Control
Log(TAD-0410899), M
compound 25
n = 6
% of Control
Log(TAD-0410386), M
compound 26
n = 3
% of Control
Log(TAD-0000031),M
compound 30
n = 1
% of Control
Log(TAD-0411745),M
compound 31
n = 13
% of Control
Log(TAD-0000032), M
compound 32
n = 3
% of Control
Log(TAD-0411448),M
compound 37
n = 2
% of Control
Log(TAD-0411473),M
compound 38
n = 1
% of Control
Log(TAD-0422576),M
compound 39
n = 3
% of Control
Log(TAD-0000082),M
compound 40
n = 3
% of Control
Log(TAD-0410391), M
compound 44
n = 5
% of Control
Log(TAD-0410392), M
compound 45
n = 1
% of Control
Log(TAD-0410616), M
compound 46
n = 2
% of Control
Log(TAD-0410530),M
compound 57
n = 3
% of Control
Log(TAD-0410407),M
compound 58

B

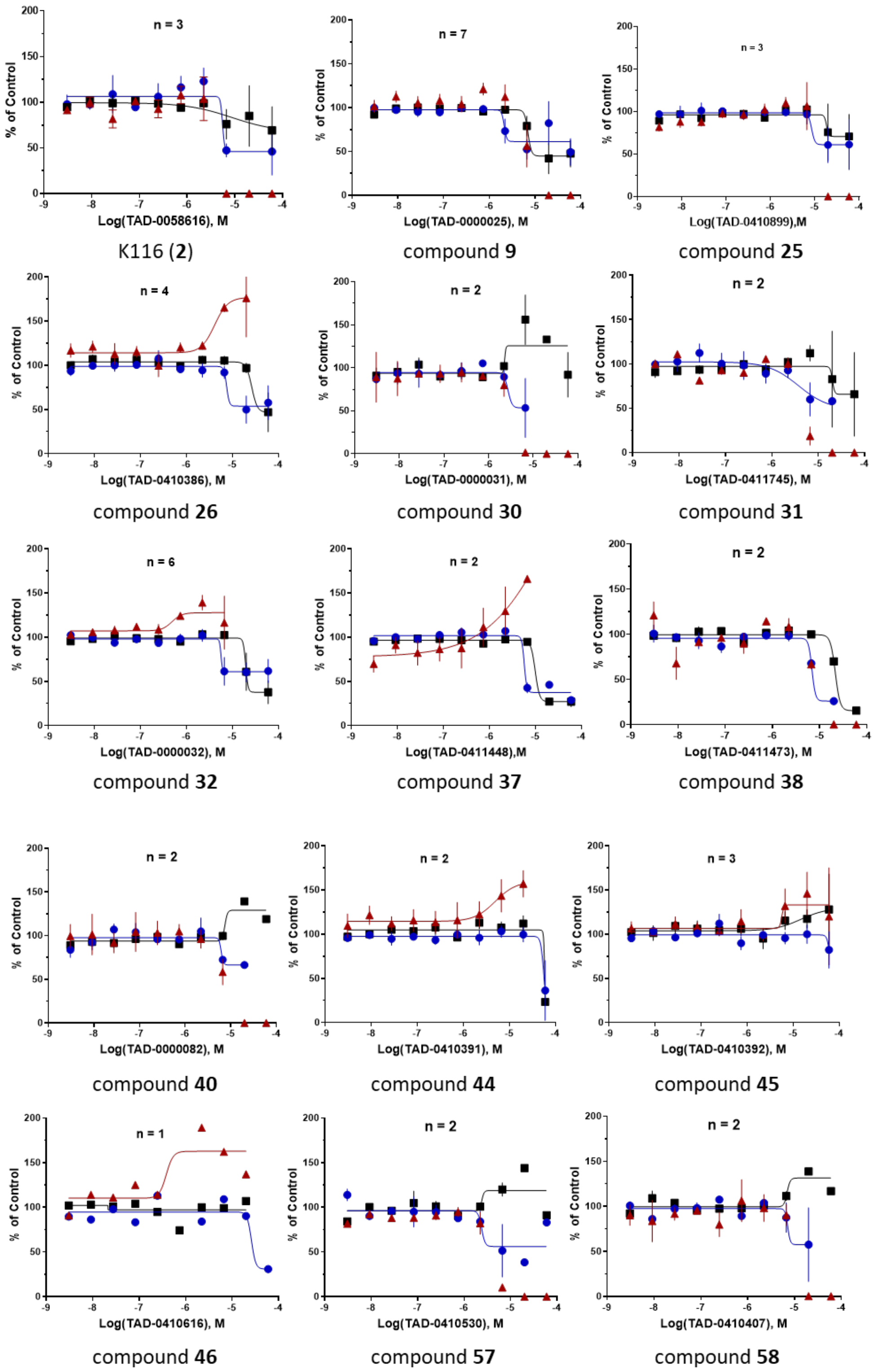


**Figure S4 | Evaluation of pHrodo-myelin/membrane debris uptake and cell health.** Cell count (●), nuclear intensity (■), and pHrodo-myelin/membrane debris uptake (▲). n represents number of experiments. **A.** BV-2 cells**. B.** Primary murine microglia.

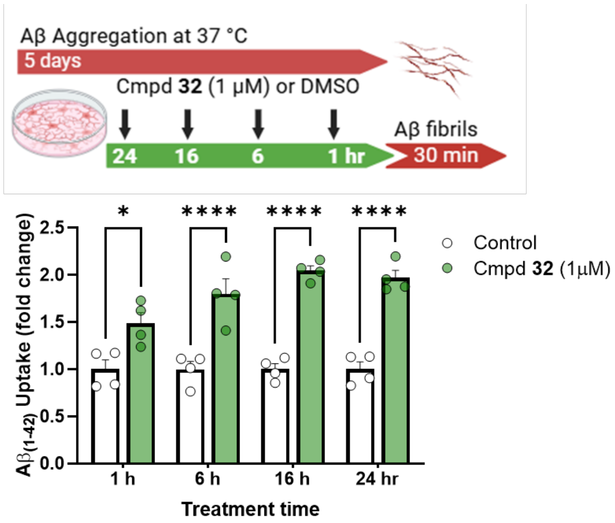


**Figure S5 | Time course of compound 32 enhancement of microglial uptake of $A\beta_{1-42}$ fibrils.** Alexa Fluor-labeled $A\beta_{1-42}$ peptides were aggregated at 37 °C for 5 days to form fibrils. Wild-type primary microglia were plated, allowed to adhere, and pretreated with 1 µM Compound **32** or vehicle for 1, 6, 16, or 24 hours. Pre-formed Aβ fibrils (0.5 µM) were then added to the culture medium. After 30 minutes, cells were washed, and internalized Aβ was quantified by immunofluorescence imaging. Bar graph shows Aβ fibril uptake (fold change vs. control) at each pretreatment time point. Data are mean ± SEM (n = 4 per group). Statistical significance was assessed using two-way ANOVA, the significant interaction by treatment and time of exposure was examined by Tukey's multiple comparisons test.*$p < 0.05$, ****$p < 0.0001$.

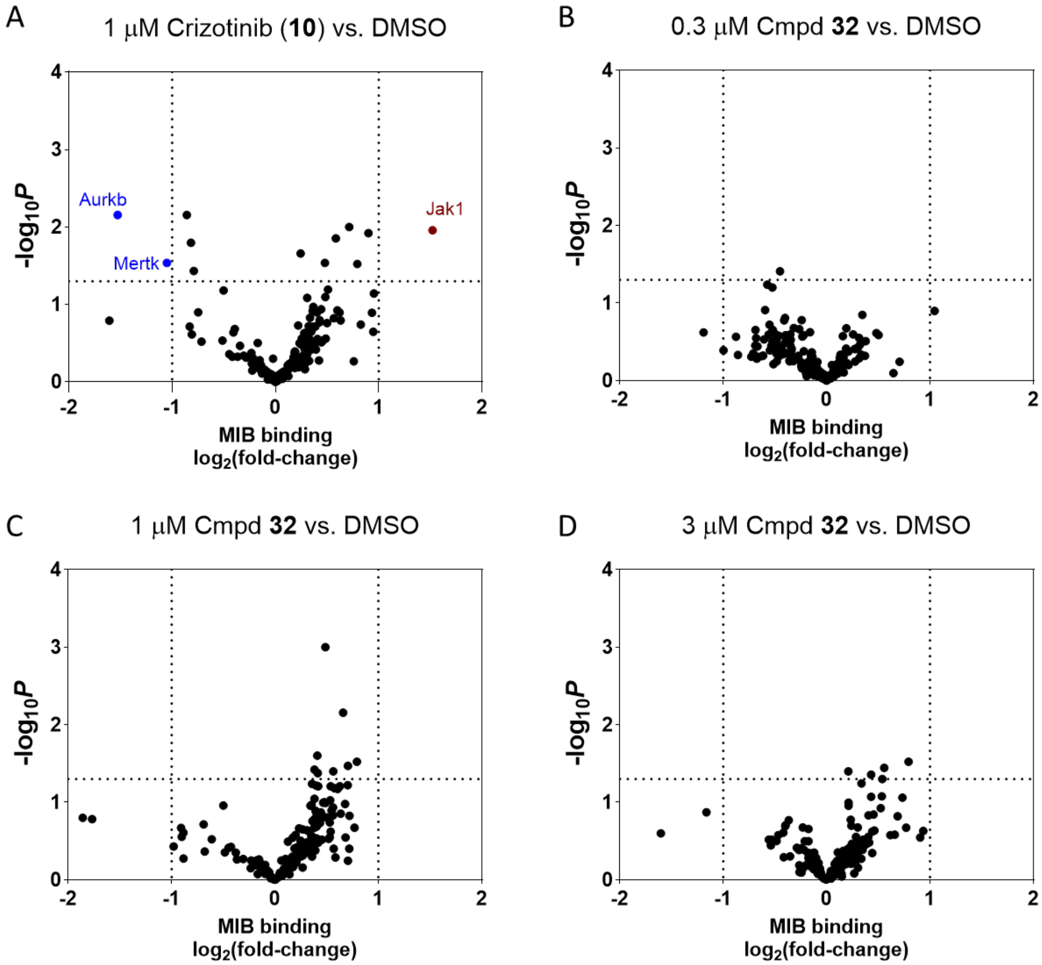


**Figure S6 | Cellular kinome profiling.** Volcano plots illustrating statistically significant changes in kinase binding to Multiplexed Inhibitor Beads (MIB). Kinases with increased or decreased MIB binding are colored in red and blue, respectively. Dashed lines indicate thresholds of statistical significance ($p < 0.05$) and $\log_2$ fold change (>1 or <−1). Data represent n = 3 biological replicates per condition. (A) Crizotinib (1 µM) treatment resulted in decreased MIB binding of Aurkb and Mertk, demonstrating engagement with these kinases as expected. Jak1 showed an increase in binding, potentially reflecting an adaptive or compensatory signaling response. (B-D) No significant changes in MIB binding following treatment with compound **32** at 0.3 µM, 1 µM, and 3 µM for 1 hour, indicating that compound **32** does not engage kinases expressed in BV2 cells.

BV2 cells were cultured in 75 $cm^2$ flasks at a density of $1 \times 10^6$ cells per flask in a serum-free medium. Cells were treated with either crizotinib (1 µM), compound **32** (0.32 µM, 1 µM, or 3 µM), or 0.1% DMSO as a vehicle control for 1 hour. Following treatment, cells were washed with 1X PBS, gently scraped, pelleted by centrifugation (500 × g, 15 minutes, 4°C), and flash frozen. Cell pellets were lysed in MIB (Multiplexed Inhibitor Beads) lysis buffer (50 mM HEPES, 150 mM NaCl, 0.5% Triton X-100, 1 mM EDTA, 1 mM EGTA, pH 7.5) supplemented with protease and phosphatase inhibitors. Lysates were clarified by centrifugation, and protein concentration was determined using the Bradford assay. Equal amounts of protein were loaded onto MIB columns containing a mixture of seven covalently linked type I kinase inhibitors on ECH-Sepharose resin, pre-equilibrated with a high-salt buffer (1 M NaCl). Bound proteins were eluted using 0.5% SDS, 1% β-mercaptoethanol, and 100 mM Tris-HCl (pH 6.8) at 95°C, reduced with DTT, alkylated with iodoacetamide, and concentrated using Amicon Ultra-4 filters (10-kDa cutoff). Proteins were then precipitated with methanol–chloroform, dried, and resuspended in 50 mM HEPES (pH 8.0), followed by overnight digestion with trypsin at 37°C. Peptides were extracted with ethyl acetate to remove residual detergent, dried, purified using C18 spin columns, and reconstituted in 0.1% formic acid for LC-MS/MS proteomic analysis conducted using a label-free quantification (LFQ) strategy[1]. Samples were individually analyzed on a Thermo Orbitrap Exploris 480 mass spectrometer using a data-dependent acquisition method. Peptide identification and protein inference were performed using MaxQuant software, matching acquired MS/MS spectra to the UniProt mouse reference database (TaxID 10090). Relative LFQ intensity values were calculated for identified proteins (e.g., Aurkb, Mertk), and the dataset was filtered to retain only annotated kinases with at least two razor+unique peptides. LFQ intensity values for each kinase were $\log_2$-transformed, and missing values were imputed from a normal distribution. Fold change was calculated based on the average $\log_2$ intensity of treatment groups (n = 3) compared to control groups (n = 3). Statistical significance was evaluated using an unpaired, two-tailed Student's t-test. Kinases with a $\log_2$ fold change greater than 1 or less than –1 and a P-value < 0.05 were considered significantly altered. These data were visualized using a volcano plot, with dashed lines marking the statistical thresholds, allowing for rapid identification of kinases exhibiting significant changes in MIB binding relative to the control following treatment. Data has been uploaded to PRIDE with accession number TBD. The mass spectrometry proteomics data have been deposited to the ProteomeXchange Consortium[2] via the PRIDE partner repository (https://www.ebi.ac.uk/pride/) with the dataset identifier TBD.

1. Adebayo AK, Bhat-Nakshatri P, Davis C, et al. Oxygen tension-dependent variability in the cancer cell kinome impacts signaling pathways and response to targeted therapies. *iScience*. Jun 21 2024;27(6):110068. doi:10.1016/j.isci.2024.110068
2. Perez-Riverol Y, Bai J, Bandla C, et al. The PRIDE database resources in 2022: a hub for mass spectrometry-based proteomics evidences. *Nucleic Acids Res*. Jan 7 2022;50(D1):D543-D552. doi:10.1093/nar/gkab1038

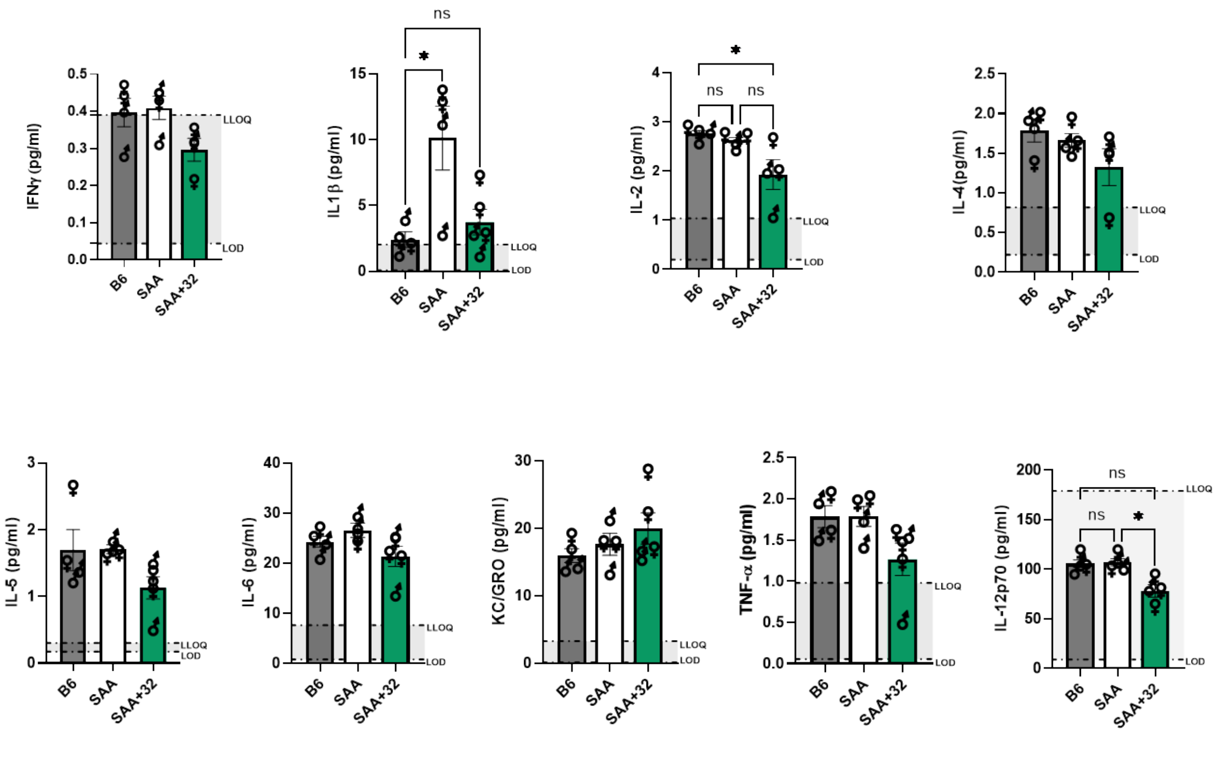


**Figure S7 | Proinflammatory Cytokine Panel Results.** Brain lysates from the acute PD study in wild-type (B6) and SAA mice treated with vehicle (SAA) or Compound **32** (SAA+32) were analyzed using the MSD Cytokine Assay and are reported as pg protein per mL of sample. The dotted line represents the lower limit of quantification (LLOQ), and limit of detection (LOD) for each analyte, as defined by the manufacturer. Nine out of ten cytokines were detected above the LOD. Concentrations below the LOD were considered undetectable and excluded from statistical analysis. Data are presented as mean ± SEM (n = 4–5 per group). Statistical analysis was performed using the Kruskal–Wallis test followed by Dunn's multiple comparisons test (*p < 0.05).

**Table S1. Phosphate 5-kinase (PIP5K) family activity**

| | | | % Enzyme Activity (relative to DMSO controls) | | | | | |
|---|---|---|---|---|---|---|---|---|
| | | Kinase: | PIP5K1A | | PIP5K1B | | PIP5K1C | |
| | | Substrate: | PI(4)P:PS | | PI(4)P | | PI(4)P:PS | |
| Compound | Molecule Name | Testing Concentration (µM) | Data 1 | Data 2 | Data 1 | Data 2 | Data 1 | Data 2 |
| 43 | TAD-0409937 | 10 | 88.76 | 90.12 | 51.84 | 51.03 | 87.19 | 86.76 |
| 43 | TAD-0409937 | 1 | 85.08 | 86.13 | 86.53 | 79.70 | 87.08 | 86.43 |
| 43 | TAD-0409937 | 0.1 | 102.17 | 106.09 | 101.46 | 118.65 | 119.01 | 117.57 |
| 32 | TAD-0000032 | 10 | 86.58 | 87.39 | 58.40 | 55.59 | 91.11 | 90.53 |
| 32 | TAD-0000032 | 1 | 87.36 | 89.77 | 99.52 | 96.84 | 88.88 | 87.65 |
| 32 | TAD-0000032 | 0.1 | 101.96 | 106.06 | 106.98 | 103.91 | 104.20 | 98.66 |
| 41 | TAD-0410749 | 10 | 84.83 | 84.33 | 57.60 | 53.98 | 93.70 | 87.40 |
| 41 | TAD-0410749 | 1 | 85.74 | 87.05 | 115.46 | 103.94 | 101.10 | 99.99 |
| 41 | TAD-0410749 | 0.1 | 105.74 | 108.92 | 103.15 | 97.62 | 100.53 | 100.13 |
| | PIK-93 | Control Compound IC50* (M): | 1.33E-04 | | ND | | 2.23E-05 | |
| | APY0201 | | ND | | 2.23E-06 | | ND | |

ND *Indicates compound not tested against enzyme

**PIP5K Lipid Kinase Panel Activity Assay.** Kinase assays were performed using Reaction Biology's Lipid kinase platform. Briefly, 5µL of 1X kinase in reaction buffer is delivered to background wells. 5µL of 1X kinase with substrate mixture (PIP5K1A substrate: PI(4)P:PS, SignalChem, Cat# P427-59, PIP5K1B substrate: PI(4)P, Cayman, Cat# 10007757, PIP5K1C substrate: PI(4)P:PS, SignalChem, Cat# P427-59) is delivered to all remaining assay wells (Reaction plate: Corning 3572, Non-Treated 384 well plates). Compounds prepared in DMSO or DMSO control are delivered into kinase reaction mixture by Acoustic technology (Echo550; nanoliter range) and are pre-incubated with kinase mixture for 20 mins. After preincubation, 50X ATP (10 µM final concentration) is delivered into the reaction mixture to initiate the reaction and the assay is incubated for 60 min at 30 °C. The kinase reaction is quenched by adding 5 µL/well ADP-Glo reagent (ADP-GloTM Kit: Promega Cat# V9103), and incubated for 40 min. 10 µL/well ADP-Glo Detection reagent is added and incubated for 30 mins at RT in dark. Luminescence signal is measured by EnVision (Revvity). The background subtracted luminescence signal is converted into µM ADP production based on ADP standard curves from each experiment. Enzyme activity in each assay well (at various compound concentrations) relative to that of DMSO wells were calculated.
